# Host-Derived Lipids Sustain Membrane Biogenesis in an Obligate Endosymbiont

**DOI:** 10.64898/2026.11.04.764378

**Authors:** Kathrine Xin Yee Tan, Dolma Michellod, Makoto Arita, Aya Hori, Tomonari Nozaki, Naoki Matsuda, Maki Kondo, Shuji Shigenobu

## Abstract

Intracellular symbionts often undergo extreme genome reduction and lose pathways considered essential in free-living bacteria, including membrane phospholipid biosynthesis. Yet how such symbionts maintain their membranes remains poorly understood. The pea aphid *Acyrthosiphon pisum* harbors the obligate endosymbiont *Buchnera aphidicola* within specialized host cells called bacteriocytes, providing a tractable system to address this question. Here we integrated untargeted lipidomics, transcriptomics, and functional assays to investigate lipid metabolism across the symbiosis. Although *Buchnera* lacks nearly all canonical genes for phospholipid biosynthesis, purified *Buchnera* contained all major phospholipid classes detected in the host, including phosphatidylcholine (PC), phosphatidylserine (PS), phosphatidylethanolamine (PE), phosphatidylglycerol (PG), and cardiolipin (CL). This repertoire differs markedly from conventional bacterial lipidomes, combining canonical bacterial phospholipids with PC, PS, and cholesterol sulfate, features more characteristic of the eukaryotic host environment. Every lipid species detected in *Buchnera* was also present in aphid tissues, consistent with extensive host contribution to the symbiont lipidome. Yet the *Buchnera* lipidome was compositionally distinct, with characteristic differences in the relative abundance of individual lipid species. Bacteriocytes were transcriptionally specialized for lipid precursor metabolism, including upregulation of phospholipid precursor synthesis and acetyl-CoA transport pathways. RNA interference targeting host lipid precursor genes reduced *Buchnera* abundance and disrupted symbiont morphology, linking host lipid metabolism to *Buchnera* maintenance. We also found that sterol availability influences *Buchnera* abundance. Together, these findings support a model in which host lipid metabolism sustains membrane biogenesis in a highly reduced endosymbiont and identify membrane lipids as an important dimension of metabolic integration in obligate intracellular symbioses.

**Significance Statement:** Obligate intracellular symbionts often lose genes considered essential for independent life, raising the question of how basic cellular functions are maintained. Remarkably, the highly reduced aphid endosymbiont *Buchnera aphidicola*, despite lacking nearly all genes for phospholipid biosynthesis, contains a diverse membrane lipid repertoire shared with its host. Host cells harboring *Buchnera* are specialized for lipid precursor metabolism, and perturbing these host pathways compromises symbiont abundance and cellular integrity. These findings reveal a previously underappreciated level of metabolic integration in which host lipid metabolism supports symbiont membrane biogenesis. Because similar losses of lipid biosynthetic capacity occur widely among intracellular symbionts, host-supported membrane biogenesis may represent a general feature of endosymbiosis evolution.

## Introduction

Symbiotic associations between insects and bacteria are widespread in nature and often involve metabolic interdependencies in which nutrient exchange enables adaptation to specialized ecological niches [1], [2], [3], [4]. Among them, the association between the pea aphid, *Acyrthosiphon pisum*, and its obligate endosymbiont *Buchnera aphidicola* stands as a paradigm of symbiotic coevolution. Aphids feed on plant phloem sap, a nutritionally unbalanced diet deficient in essential amino acids, and rely on *Buchnera* to supplement these deficiencies [4], [5], [6]. *Buchnera* resides intracellularly within specialized host cells called bacteriocytes, where each bacterial cell is encapsulated by a host-derived symbiosomal membrane. Bacteriocytes are organized into an organ-like structure, the bacteriome, which consists of two bilateral halves extending along the body and joining above the hindgut. *Buchnera* is vertically transmitted from mother to offspring [7]. The association is strictly obligate and mutualistic, with neither partner able to complete normal reproduction in the absence of the other [7], [8], [9]. Over its long evolutionary history, *Buchnera* has undergone extensive genome reduction, loosing many genes considered essential in free-living bacteria while retaining those for its obligate intracellular and mutualistic lifestyle [6], [10], [11], [12], [13]. Genomic and physiological studies have revealed extensive metabolic complementation between the partners: the host supplies substrates and metabolic precursors, whereas *Buchnera* retains biosynthetic pathways that produce essential amino acids and other nutrients that are limited in the aphid diet [5], [14]. Beyond these well-characterized nutritional interactions, however, highly reduced endosymbionts have been proposed to function as integrated components of host cellular systems, with functions lost from the symbiont genome effectively compensated by the host [5], [14], [15]. How such integration extends to lipid metabolism, whose products serve diverse functions as energy reserves, signaling molecules, and structural components of cellular membranes, remains largely unresolved.

In Gram-negative bacteria, phospholipids such as phosphatidylethanolamine (PE), phosphatidylglycerol (PG), and cardiolipin (CL) are major components of the cell envelop and are crucial for membrane integrity and function[16], [17], [18]. Genomic evidence points to a striking asymmetry in lipid metabolism between the two partners. Whereas the aphid host retains the major pathways required for fatty acid and phospholipid biosynthesis, *Buchnera* has lost nearly all genes required for phospholipid biosynthesis, retaining only a cardiolipin synthase (*clsA*) homolog [6], [19], [20], [21]. Despite this extreme genetic reduction in biosynthetic capacity, *Buchnera* cells exhibit a bacterial inner and outer membrane structure, and each bacterial cell is further encapsulated by a layer of host-derived symbiosomal membrane [22] [23]. Thus, *Buchnera* retains structurally intact bacterial membranes despite near-complete phospholipid auxotrophy, raising what we recently termed a “membrane paradox” [20]. Whether the host supplies membrane lipids or their precursors to *Buchnera*, and how this metabolic asymmetry is reflected in the symbiont lipid composition, have remained unresolved.

To address these questions, including “membrane paradox”, we here combined untargeted lipidomics of *Buchnera* and aphid tissues with bacteriocyte transcriptomic profiling and functional assays. We show that *Buchnera* retains a broad, host-shared lipid repertoire despite its severely reduced lipid biosynthetic capacity, while bacteriocytes are transcriptionally specialized for lipid precursor metabolism. Functional assays using RNA interference (RNAi) and cholesterol supplementation further link host lipid metabolism to *Buchnera* maintenance in vivo. Together, these findings support a model in which host lipid metabolism contributes to *Buchnera* membrane biogenesis and identify membrane lipid metabolism as an important dimension of metabolic integration in obligate intracellular symbioses.

## Results

### Untargeted lipidomics reveals a largely conserved lipid repertoire across *Buchnera* and aphid tissues

We performed LC–MS/MS-based untargeted lipidomics of the pea aphid (whole-body, appendage, and bacteriocyte samples) and purified *Buchnera* to compare lipid composition across host tissues and the endosymbiont. Untargeted lipidomic profiling identified a total of 355 lipid species [figshare: LC-MS/MS lipidomics dataset]. Among these, 334 lipid species were shared among aphid tissues and *Buchnera* samples, indicating a broadly conserved lipid repertoire between host tissues and the endosymbiont (Fig. S3). Notably, all lipid species identified in *Buchnera* were also detected in the aphid lipidome. Cholesterol sulfate, a eukaryotic sterol derivative, was also detected in the purified *Buchnera* fraction.

A class-based heatmap of mean lipid relative abundance showed substantial overlap in lipid classes across all sample types, while also revealing marked differences in their relative abundance (Fig. 1A). Principal component analysis (PCA) further separated samples according to biological sample type, with *Buchnera* forming a distinct cluster from host tissues (PC1: 48.4 % and PC2: 24.5 %; Fig. 1B). This separation was supported by PERMANOVA (R² = 0.892, *p* = 0.001), confirming significant differences in overall lipidomic profile among sample types. Together with the extensive overlap in detected lipid species (Fig. S3A), these results indicate that lipidomic divergence is driven primarily by differences in relative abundance of a shared lipid repertoire rather than qualitative differences in lipid composition (i.e., the presence or absence of specific lipid species].

**Fig. 1.**
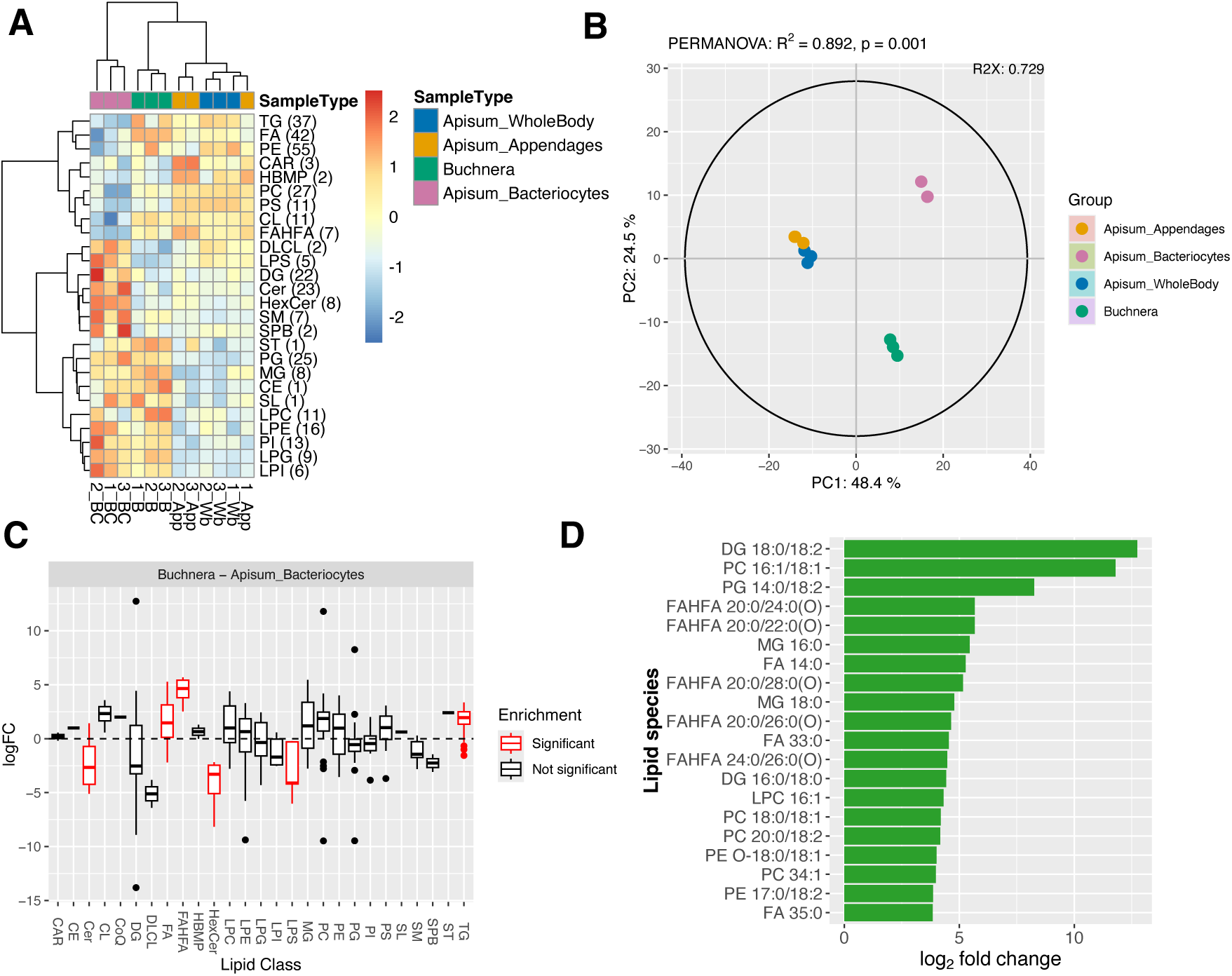
Comparative lipidome analysis revealed conserved lipid repertoire across *Buchnera* and aphid host. **(A)** Heatmap of lipid class relative abundance across *A. pisum* whole body (WB), appendages (App), bacteriocytes (BC) and isolated *Buchnera* cells (B). Each row represents a lipid class while numbers in parentheses indicate the count of lipid species per class. Each column represents a biological replicate. Abundances were averaged within each class, scaled by row (*z*-score) and hierarchically clustered using Ward’s D^2^ method with Euclidean distance. *Buchnera* (green) shares an overall class composition with host tissues but displays distinct relative abundances. **(B)** PCA showing distinct clustering of sample types (PERMANOVA, *R^2^* = 0.892, *p* = 0.001), with PC1 and PC2 explaining 48.8 % and 24.5 % of total variance, respectively. *Buchnera* separates clearly from host-derived samples, indicating a compositionally distinct lipidome. **(C)** LSEA comparing *Buchnera* and aphid bacteriocytes identifies lipid classes significantly enriched or depleted (red) in *Buhcnera* based on log2FC (FDR < 0.05). The dashed line indicates enrichment (log2 FC > 0) or depletion (log2 FC < 0). **(D)** Top 20 lipid species strongly enriched in *Buchnera* relative to bacteriocytes, ranked by log2 FC. Several enriched species belong to phospholipid and fatty acid ester subclasses.Lipid-class abbreviations: CAR, acylcarnitine; CE, cholesteryl ester; Cer, ceramide; CL, cardiolipin; DG, diacylglycerol; DLCL, dilysocardiolipin; FA, fatty acid; FAHFA, fatty acid ester of hydroxyl fatty acid; HBMP, hemibismonoacylglycerophosphate; HexCer, hexosylceramide; LPC, lysophosphatidylcholine; LPE, lysophosphatidylethanolamine; LPG, lysophosphatidylglycerol; LPI, lysophosphatidylinositol; LPS, lysophosphatidylserine; MG, monoacylglycerol; PC, phosphatidylcholine; PE, phosphatidylethanolamine; PG, phosphatidylglycerol; PI, phosphatidylinositol; PS, phosphatidylserine; SM, sphingomyelin; SPB, sphingoid base; TG, triacylglycerol, ST, steroid conjugate (cholesterol sulfate), CoQ, Coenzyme Q, SL, sulfonolipid.

To identify the lipid class features underlying this divergence, we compared *Buchnera* with bacteriocytes, the endosymbiont intracellular niche, using differential abundance analysis and Lipid Set Enrichment Analysis (LSEA) (FDR < 0.05, |log_2_FC| > 1) (Fig. 1C; see SI Appendix, Fig. S4-6). Fatty acids (FA), fatty acid ester of hydroxyl fatty acid (FAHFA) and triacylglycerols (TG) were significantly enriched in *Buchnera*, whereas ceramides (Cer), hexosylceramide (HexCer) and lysophosphatidylserine (LPS) were depleted (Fig. 1C-D). In addition, *Buchnera* exhibited enrichment of fully saturated species and depletion of 17-carbon acyl chains (Fig. S3C,D). Detailed analyses of individual lipid species, acyl-chain composition and pairwise tissue comparisons are provided in Figs. S3–S6. Collectively, these results show that the major class-level differences between *Buchnera* and bacteriocytes are associated primarily with non-phospholipid lipids.

### Major phospholipid classes are detected in *Buchnera* despite the loss of phospholipid biosynthetic genes

To investigate membrane lipids in the aphid-*Buchnera* symbiosis, we next focused on glycerophospholipids, the primary structural lipids of cell membranes. Genome analysis revealed that *Buchnera* lacks nearly all genes required for phospholipid biosynthesis (SI Appendix, Fig. S7A) [6], [24], [25]. In addition, *Buchnera* lacks key genes of the fatty acid synthesis (FAS II) pathway, indicating that it is incapable of de novo membrane lipid synthesis (SI Appendix, Fig. S7B). Despite lacking the genetic capacity for de novo phospholipid synthesis, purified *Buchnera* fractions contained all major phospholipid classes, including phosphatidylcholine (PC), phosphatidylserine (PS), phosphatidylethanolamine (PE), phosphatidylglycerol (PG), and cardiolipin (CL) (Fig. 2A).

**Fig. 2.**
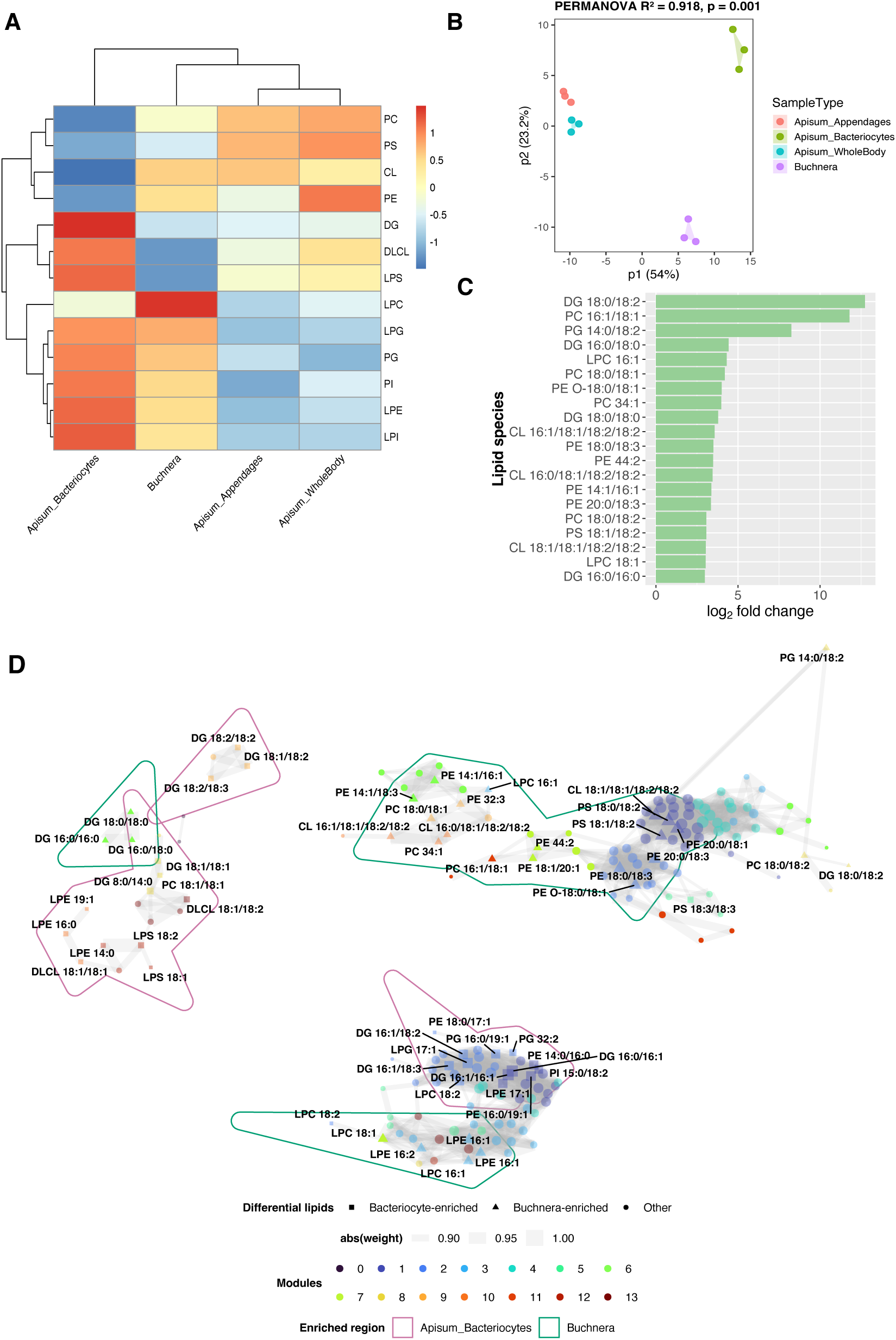
*Buchnera* retained major phospholipids, which exhibited distinct distribution pattern. **(A)** Heatmap of phospholipid class abundance across *A. pisum* whole body (Wb), appendages (App), bacteriocytes (BC), and isolated *Buchnera* cells (B). Each row represents a phospholipid class and each column a biological sample type. Abundances were summed per class and z-score by row. **(B)** PCA of all phospholipid species shows clear separation among tissue types (PERMANOVA, *R²* = 0.918, *p* = 0.001). **(C)** Top 20 phospholipid species significantly enriched in *Buchnera* relative to bacteriocytes (log₂ FC> 1, adj. *p* < 0.05). **(D)** Correlation network of phospholipid species (Spearman ρ ≥ 0.85, *p* < 0.05). Nodes represent individual lipid species and are coloured according to their correlation module identified by hierachical clustering. Node size is proportional to network connectivity. Triangles and squares denote the top 30 lipid species significantly enriched in *Buchnera* and bacteriocytes, respectively. Edge width is proportional to the strength of the Spearman correlation. Coloured hulls outline clusters of differentially enriched lipids identified by DBSCAN clustering in the network layout, highlighting spatial groupings of *Buchnera*-enriched (green) and bacteriocyte-enriched (pink) lipid species. The largely distinct distribution of the two enrichment groups suggests that lipids enriched in *Buchnera* and bacteriocytes occupy different regions of the correlation network, reflecting tissue-specific patterns of lipid co-variation. Lipid-class abbreviations: PA, phosphatidic acid; DG, diacylglycerol; PC, phosphatidylcholine; PE, phosphatidylethanolamine; PG, phosphatidylglycerol; PI, phosphatidylinositol; PS, phosphatidylserine; LPC, lysophosphatidylcholine; LPE, lysophosphatidylethanolamine; LPG, lysophosphatidylglycerol; LPI, lysophosphatidylinositol; LPS, lysophosphatidylserine; CL, cardiolipin; DLCL, dilysocardiolipin.

Lipidomic profiling of phospholipids further revealed that phospholipid class abundance varied markedly across host tissues and *Buchnera* (Fig. 2A). PCA clearly separated samples by tissue type (PERMANOVA, R² = 0.918, *p* = 0.001) (Fig. 2B), underscoring the strong tissue specificity of the phospholipid landscape. At the molecular species level, differential abundance analysis between *Buchnera* and bacteriocytes identified several phospholipid species significantly enriched in *Buchnera*, including DG 18:0/18:2, PC 16:1/18:1 and PG 14:0/18:2 (Fig. 2C). Together, these results indicate that although host tissues and *Buchnera* share a similar phospholipid repertoire, they differ in the relative abundance of individual phospholipid species.

To examine how individual phospholipid species are coordinated across tissues, we constructed a correlation-based network (Fig. 2D), using phospholipid species exhibiting strong positive correlations across tissues (Spearman’s *r* ≥ 0.85, *p* < 0.05). The resulting network comprised several interconnected modules rather than random distribution. Overlaying the top 30 phospholipids enriched in *Buchnera* and bacteriocytes, identified by differential abundance analysis (supplementary Dataset S1), revealed distinct but partially connected clusters corresponding to each compartment. For instance, *Buchnera*-enriched phospholipids formed a compact cluster containing CL- and PG-rich species, whereas bacteriocyte-enriched phospholipids occupied a neighbouring but distinct region enriched in DG- and PE-containing species (Fig. 2D). A narrow overlapping zone connected these two clusters, suggesting coordination between host and symbiont despite compartment-specific phospholipid enrichment.

### Bacteriocytes are transcriptionally specialized for lipid precursor synthesis

To investigate how the host supports *Buchnera* membrane phospholipid homeostasis despite its near-complete loss of phospholipid biosynthetic genes, we analyzed the transcriptome of aphid bacteriocytes, the specialized host cells that harbor the endosymbiont. We generated transcriptomes from purified bacteriocytes and whole-body samples of the three pea aphid morphs—viviparous females, oviparous females, and males—by deep RNA sequencing. PCA of the resulting transcriptomes showed clear separation by tissue type (PERMANOVA, R² = 0.79, *p* = 0.001) (Fig. 3A), indicating that bacteriocytes possess a transcriptional program distinct from that of other host tissues.

**Fig. 3.**
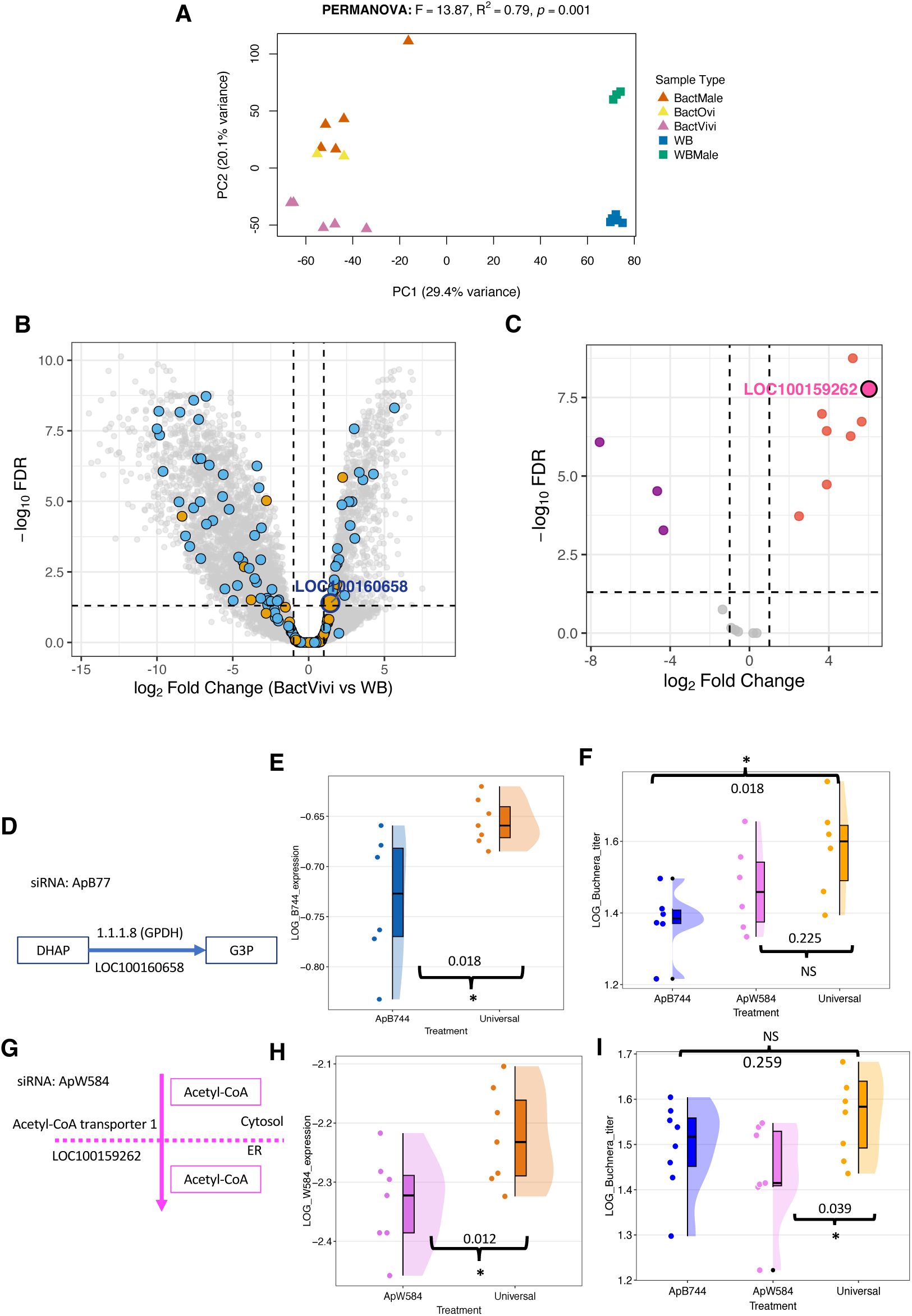
Bacteriocyte transcriptome reveals enrichment of lipid precursor synthesis **(A)** PCA of aphid transcriptomes from bacteriocytes (BactOvi, BactVivi, BactMale) and whole body samples (WB, WBMale) shows distinct clustering by tissue type (PERMANOVA, *R²* = 0.79, *p* = 0.001). **(B)** Differential expression analysis (BactVivi vs WB) identifies genes enriched for lipid metabolic processes (GO:0006629; blue dots) and phospholipid metabolism (GO:0006644; yellow dots), including glycerol-3-phosphate dehydrogenase (GPDH; LOC100160658), which synthesizes the phospholipid precursor sn-glycerol-3-phosphate. **(C)** Volcano plot showing upregulation of acetyl-CoA transporter 1 (*AT-1*; LOC100159262) in bacteriocytes (upregulated: pink dots; downregulated: purple dots). (D–I) Functional validation by siRNA interference targeting *GPDH* (ApB744) and AT-1 (ApW584). **(D)** illustrates the targeted steps: GPDH catalyzes DHAP → G3P. **(E)** At 12 µM siRNA for 24 h, GPDH expression was significantly lower than the Universal control (ApB744 = –0.733 ± 0.067 [mean ± SD], *n* = 6; Universal = – 0.655 ± 0.023, *n* = 7; Welch’s one-tailed *t*-test, *t*(6.0139) = –2.693, *p* = 0.018). **(F)** *GPDH* knockdown at 20 µM 50 h post-treatment significantly reduced *Buchnera* titer (1.38 ± 0.091 (ApB744), 1.47 ± 0.123 (ApW584), and 1.58 ± 0.135 [mean ± SD], (Universal; *n* = 6) (ANOVA, *F*(2, 15) = 4.393, *p* = 0.032; post hoc test: Dunnett’s test). **(G)** AT-1 transfers acetyl-CoA from the cytosol to the endoplasmic reticulum. **(H)** *AT-1* expression was significantly reduced under RNAi treatment (ApW584 = –2.34 ± 0.080 [mean ± SD]; Universal = –2.22 ± 0.083, *n* = 7; one-tailed Student’s *t*-test, *t*(12) = –2.560*, p* = 0.012). **(I)** *Buchnera* titers declined significantly following *AT-1* knockdown at 12 µM (30h post-treatment). Mean titers were 1.49 ± 0.099 (mean ± SD, *n* = 8) for ApB744, 1.44 ± 0.114 (*n* = 7) for ApW584, and 1.57 ± 0.095 (*n* = 8) for the Universal control (ANOVA, *F*(2, 20) = 3.19, *p* = 0.063; post hoc test: Dunnett’s test).These results indicate that both host genes are essential for sustaining symbiont proliferation through lipid-precursor supply. Each data point represents one biological replicate consisting of three pooled aphid nymphs. Asterisks indicate statistical significance*: p* < 0.05 (*); NS, not significant.

Differential expression analysis comparing bacteriocytes and whole-body samples of viviparous female identified 999 upregulated and 2,633 downregulated genes (FDR < 0.05, |log_₂_FC| > 1). Gene Ontology (GO) enrichment analysis with GOseq indicated that bacteriocytes are trascriptionally specialized for metabolite transport and membrane-associated functions, with transporter activity, transmembrane transport, and membrane components among the most significantly enriched categories (SI Appendix, Fig. S8). Regarding lipid-related processes, semantically non-redundant GO enrichment analysis using GOfuncR identified lipid metabolic process (GO:0006629) and phospholipid metabolic process (GO:0006644) as significantly enriched categories (Fig. 3B; supplementary Dataset S1).

To place these transcriptional differences in the context of phospholipid metabolism, we integrated bacteriocyte transcriptomic data with the *Buchnera* lipidome using the KEGG glycerophospholipid pathway (api00564) as a common reference framework (Fig. 5; see also SI Appendix, Fig. S11 and Supplementary Dataset S1). Pathway mapping showed that genes upregulated in bacteriocytes are concentrated in the upstream portion of the glycerophospholipid biosynthetic pathway, from dihydroxyacetone phosphate (DHAP) through glycerol-3-phosphate (G3P), phosphatidic acid (PA), CDP-diacylglycerol (CDP-DAG), and diacylglycerol (DG). In particular, *GPDH (LOC100160658*), *GPAT1 (LOC100166666), Ppap2a (LOC100164616),* and *CDS (LOC100163059*), which catalyze sequential steps in phospholipid precursor synthesis, were significantly upregulated, whereas phospholipase-related genes such as *GIIIspla2 (LOC100569429)* were downregulated (Fig. 5). These expression patterns indicate coordinated transcriptional activation of host pathways responsible for phospholipid precursor synthesis.

We next focused on the acetyl-CoA transporter (*AT-1*) gene family, which mediates the transport of acetyl-CoA from the cytosol to the endoplasmic reticulum (ER). Based on eggNOG orthology annotations, the *A. pisum* genome encodes 25 *AT-1* paralogs, representing a marked expansion of this gene family compared with other insects [26], [27]. According to edgeR analysis (Supplementary Dataset S1), transcripts from 17 paralogs were detected, several of which were strongly upregulated in bacteriocytes, most notably *LOC100159262* (Fig. 3C). These findings suggest that the expanded *AT-1* gene family has undergone functional diversification associated with acetyl-CoA trafficking in bacteriocytes, potentially associated with lipid precursor metabolism.

### Host lipid precursor pathways are required for *Buchnera* maintenance

To test whether host lipid precursor pathways identified by RNA-Seq contribute to *Buchnera* maintenance, we performed siRNA-mediated knockdown of two host genes: glycerol-3-phosphate dehydrogenase (GPDH; LOC100160658), which catalyzes the conversion of dihydroxyacetone phosphate (DHAP) to glycerol-3-phosphate (G3P; Fig. 3D), and AT-1 (LOC100159262), which transports acetyl-CoA into the ER (Fig. 3G). Both transcripts were significantly reduced within 24 h following siRNA treatment (Fig. 3E, H).

Knockdown of *AT-1* using 12 µM siRNA led to a significant decrease in *Buchnera* titer after 30 h (Fig. 3I). In contrast, *GPDH* knockdown produced a comparable reduction only at a higher siRNA concentration (20 µM) and prolonged incubation (50 h; Fig. 3F).

Super-resolution imaging further revealed morphological abnormalities of *Buchnera* cells following knockdown of both genes (SI Appendix, Fig. S9A–G). *AT-1* knockdown (ApW584) produced marked morphological distortions, including irregular cell outlines and uneven DAPI staining, whereas *GPDH* knockdown (ApB744) caused milder but similar effects. In contrast, aphids treated with Universal siRNA controls retained uniformly spherical *Buchnera* cells. These results provide functional evidence that host lipid precursor pathways contribute to *Buchnera* maintenance.

### Cholesterol promotes *Buchnera* proliferation in vivo

In addition to phospholipid precursors, our lipidomic analysis detected cholesterol sulfate (ST 27:1;O;S), a eukaryotic sterol derivative, in purified *Buchnera* fractions (Fig. 1A). Because sterol is a membrane component characteristic of eukaryotes and generally absent from bacteria, this observation prompted us to examine whether host-derived sterols also influence *Buchnera* maintenance. We therefore supplemented aphids with cholesterol and quantified *Buchnera* abundance in vivo.

Cholesterol supplementation did not affect *Buchnera* titers after 24 h (Fig. 4A). By 48 h, however, a significant increase in *Buchnera* titer was observed in the cholesterol-treated group relative to negative controls (Fig. 4B). Super-resolution imaging revealed no obvious changes in bacteriocyte organization following cholesterol supplementation (SI Appendix, Fig. S9H–K). These results demonstrate that cholesterol availability promotes *Buchnera* proliferation in vivo.

**Fig. 4.**
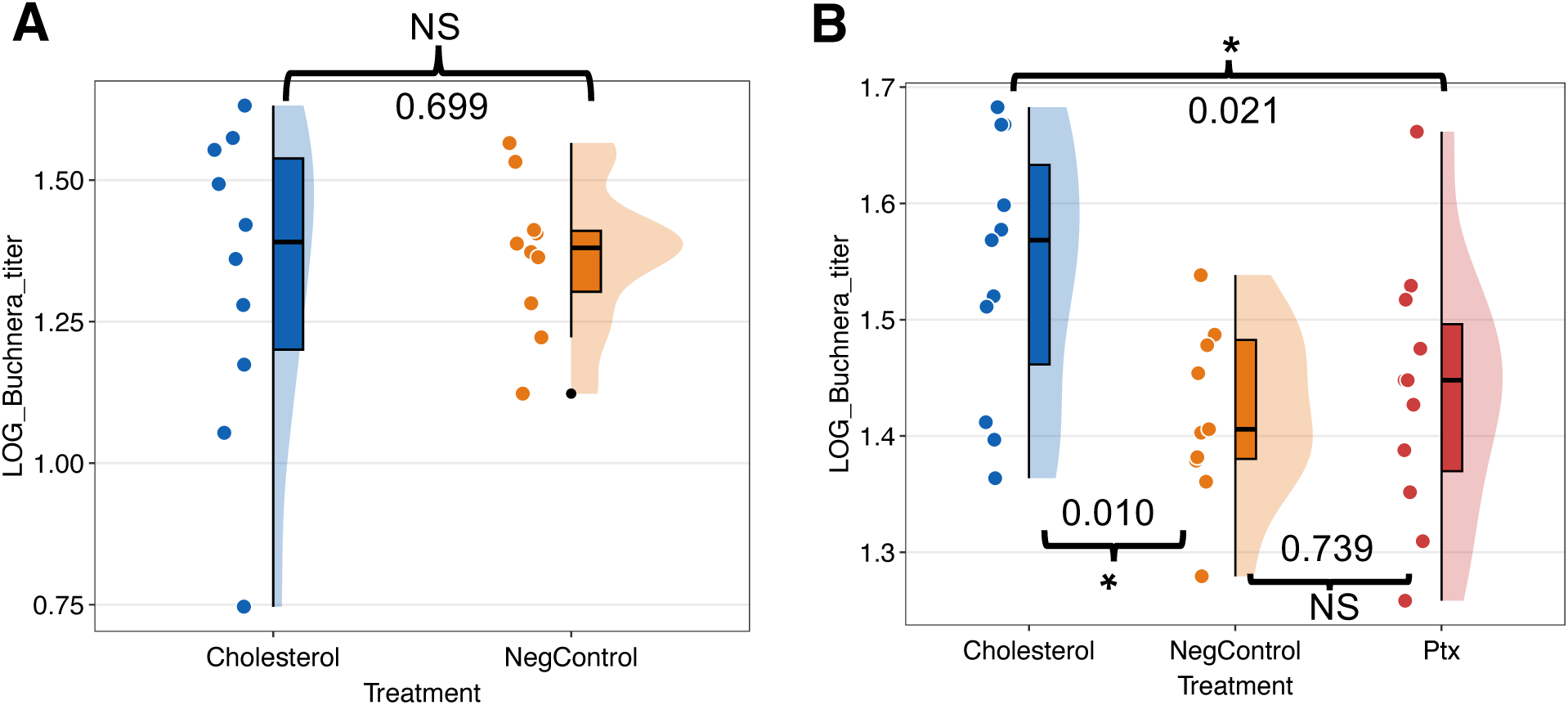
Cholesterol supplementation promotes *Buchnera* proliferation over time. **(A)** Cholesterol supplementation did not affect *Buchnera* titer at 24 h post-treatment (Cholesterol = 1.33 ± 0.275 [mean ± SD], NegControl = 1.37 ± 0.133, *n* = 10; two-tailed Student’s *t*-test, *t*(18) = –0.393, p = 0.699). (**B)** Significant titer increment was observed at 48 h compared with negative controls (NegControl and Ptx) (Cholesterol = 1.54 ± 0.113 [mean ± SD], NegControl = 1.42 ± 0.074, Ptx = 1.44 ± 0.112, *n* = 11; ANOVA, *F*(2, 30) = 4.55, *p* = 0.019; post hoc LSD test)t). Each dot represents one biological replicate comprised of a single treated aphid nymphs. Asterisks indicate statistical significance: *p* < 0.05 (*); NS, not significant.

## Discussion

Extremely reduced bacterial endosymbionts challenge conventional notions of cellular autonomy, because they have lost genes traditionally considered essential for core processes, including membrane biogenesis. Conceptual syntheses have proposed that such symbionts function as integrated components of host cellular systems, with missing functions effectively outsourced to the host [5], [15], [21], [28], [29]. Although metabolic complementation in amino acid and vitamin metabolism is well established in insect–bacterial symbioses, how hosts compensate for the loss of symbiont membrane lipid biosynthesis remains largely unresolved [5], [6], [15], [20], [28]. This problem is particularly striking in the aphid–*Buchnera* association. *Buchnera* has lost most canonical genes required for phospholipid biosynthesis, de novo fatty acid synthesis, unsaturated fatty acid production, and fatty acid degradation, while retaining only a minimal set of lipid-related functions [6]. We recently framed this host-symbiont asymmetry in lipid biosynthetic capacity as a “membrane paradox”—the maintenance of intact bacterial membranes despite near-complete phospholipid auxotrophy—and proposed a host-dominated organization of lipid metabolism in this symbiosis [20]. Here, by integrating lipidomics, transcriptomics, and functional perturbations, we provide experimental support for this model and show that host lipid metabolism contributes to symbiont membrane biogenesis and maintenance.

### *Buchnera* harbors a diverse, noncanonical lipidome despite severe loss of lipid biosynthetic capacity

A striking finding of this study is the apparent mismatch between the extremely reduced lipid biosynthetic capacity of *Buchnera* and its actual phospholipid composition. Although *Buchnera* lacks nearly all canonical genes required for phospholipid biosynthesis and retains only *clsA*, purified *Buchnera* fractions contained all major phospholipid classes detected in the host, including PC, PS, PE, PG, and CL (Fig. 2A). Thus, the loss of endogenous biosynthetic pathways has not resulted in a corresponding loss of membrane phospholipid diversity. This decoupling between biosynthetic capacity and lipid content strongly supports the idea that *Buchnera* depends on host-derived lipids or lipid precursors for membrane biogenesis.

The lipidomic profile of *Buchnera* markedly diverges from those of free-living bacteria. In *Escherichia coli*, a related Gram-negative bacterium, membranes are composed predominantly of PE (70-80%), with smaller proportions of PG (15 - 20 %) and CL (5-10 %) [16], [18], [30], [31]. In contrast, our lipidome profiling revealed that *Buchnera* contains a broader mixture of phospholipid classes, PC and PS, in addition to the canonical bacterial phospholipids PE, PG and CL. Several molecular species, including PC 16:1/18:1, PS 18:1/18:2, PG 14:0/18:2, and CL 16:1/18:1/18:2/18:2, together with DG 18:0/18:2, were significantly enriched in *Buchnera* relative to bacteriocytes (Fig. 2C). *Buchnera* lipidome was also enriched in fully saturated lipid acyl chains and contained cholesterol sulfate, a eukaryotic sterol derivative that is unusual in bacterial lipidomes because sterols are generally absent from bacterial membranes [18], [30], [32]. Thus, *Buchnera*, despite being a member of the Gammaproteobacteria, does not simply retain a conventional bacterial lipid profile. Rather, its lipid composition combines canonical bacterial phospholipids with lipid features more characteristic of the eukaryotic host environment.

### The *Buchnera* lipidome is host-shared but compositionally distinct

Comparative lipidomics revealed that *Buchnera* and aphid tissues share a common lipid repertoire, while differing markedly in the relative abundance of individual lipid species (Fig. 1). Indeed, nearly all detected lipid species were shared between host tissues and *Buchnera*, and every lipid species detected in *Buchnera* was also present in the aphid lipidome (Fig. 1A). Thus, the *Buchnera* lipidome is not defined by a fundamentally distinct set of lipids, but rather by differential abundance within a host-shared repertoire. At the same time, the clear separation of *Buchnera* from host tissues in multivariate analyses (Fig. 2B) and the compartment-specific enrichment of individual lipid species indicate that its lipid composition does not merely mirror that of the surrounding bacteriocyte (Fig. 2D). Rather, *Buchnera* exhibits a compositionally distinct lipid profile, characterized by differential enrichment of several phospholipid classes and molecular species, including PC, PS, PG, and CL (Fig. 2A–B). Earlier work using thin-layer chromatography coupled with gas–liquid chromatography provided the first evidence for presence of PE and PC in *Buchnera* [32]. Our high-resolution LC–MS/MS analysis substantially extends these observations and demonstrates that the *Buchnera* lipidome is broadly embedded within the host lipid repertoire while maintaining a distinct compositional profile. Together, these findings are consistent with extensive host dependence for membrane lipid acquisition, while suggesting that lipid accumulation in *Buchnera* is non-uniform and shaped by the intracellular symbiotic context.

### Retention of *clsA* and detection of cardiolipin mark a residual bacterial lipid signature in *Buchnera*

Despite the strong contribution of the eukaryotic host environment to the *Buchnera* lipidome, one recognizable phospholipid biosynthesis gene is retained in its highly reduced genome: *clsA*. ClsA is annotated as a cardiolipin synthase, and cardiolipin, a canonical bacterial membrane lipid, was readily detected in the *Buchnera* lipidome (Fig. 2A). The aphid genome encodes several enzymes associated with cardiolipin metabolism, primarily for mitochondrial membrane, including a probable CMP-forming cardiolipin synthase (LOC100161189), a tafazzin homolog (LOC100165481), and a lysocardiolipin acyltransferase 1-like protein (LOC115032990), but none of those genes was differentially expressed in viviparous bacteriocytes [19], [33], [34]. In contrast, our previous functional study showed that disruption of *cls* impairs *Buchnera* membrane integrity, causes severe morphological abnormalities and mislocalization, and reduces symbiont abundance [35]. Thus, host dependence does not appear to simply replace the bacterial membrane lipid system. Rather, *Buchnera* seems to maintain a mosaic-like membrane composition in which extensive host-derived lipid provisioning coexists with a minimal but functionally important bacterial lipid biosynthetic capacity centered on cardiolipin, a bacterial phospholipid also characteristic of mitochondrial membranes. This unusual lipid composition may have physiological consequences. The enrichment of saturated lipid species, together with the presence of a sterol derivative, could influence membrane physical properties, including membrane rigidity, whereas bacterial phospholipids such as CL and PG have established roles in membrane organization and have been implicated in cell division, including Z-ring assembly [17], [36], [37], [38], [39], [40]. These properties provide plausible links between lipid composition and *Buchnera* membrane integrity and proliferation, although their direct biophysical and cellular effects remain to be tested.

### Host transcriptional specialization supports symbiont lipid metabolism

RNA-Seq analysis revealed pronounced transcriptional specialization of bacteriocytes, highlighting host-side organization of lipid precursor metabolism relevant to *Buchnera* membrane biogenesis (Fig 5). Genes involved in metabolite transport, membrane-associated functions, and lipid metabolism were enriched in bacteriocytes, and pathway-level integration revealed coordinated upregulation of several enzymes acting in the upstream steps of phospholipid precursor synthesis, including GPDH, GPAT1, Ppap2a, and CDS. This pattern is particularly notable given the extreme metabolic reduction of lipid metabolism in *Buchnera* [20]. The symbiont retains only a single phospholipid synthesis-related gene, *cls*, while genes required for upstream precursor produccion are almost absent. *Buchnera* lacks a recognizable glycerol-3-phosphate dehydrogenase (*glpD*) required to transform DHAP into G3P, a key precursor for phospholipid synthesis [6], [19], [41]. Thus, the bacteriocyte expression profile is consistent with the idea that bacteriocytes are metabolically configured to generate and handle phospholipid precursors that *Buchnera* can no longer synthesize autonomously. In addition, several *AT-1* paralogs are strongly upregulated in bacteriocytes (Fig. 3C, Dataset 01), suggesting specialization of host acetyl-CoA handling in this symbiont-housing cell type. This pattern may reflect regulatory diversification following gene family expansion, although its precise connection to symbiont lipid supply remains to be established [42], [43]. Together, these findings support a host-dominated organization of lipid metabolism, in which bacteriocytes are transcriptionally specialized for upstream lipid metabolism, whereas *Buchnera* has relinquished most of its own phospholipid biosynthetic machinery. This organization differs from the clearly partitioned metabolic complementation described for amino acid biosynthesis in the aphid–*Buchnera* symbiosis and instead supports a predominantly host-dominated organization of lipid metabolism.

**Fig. 5.**
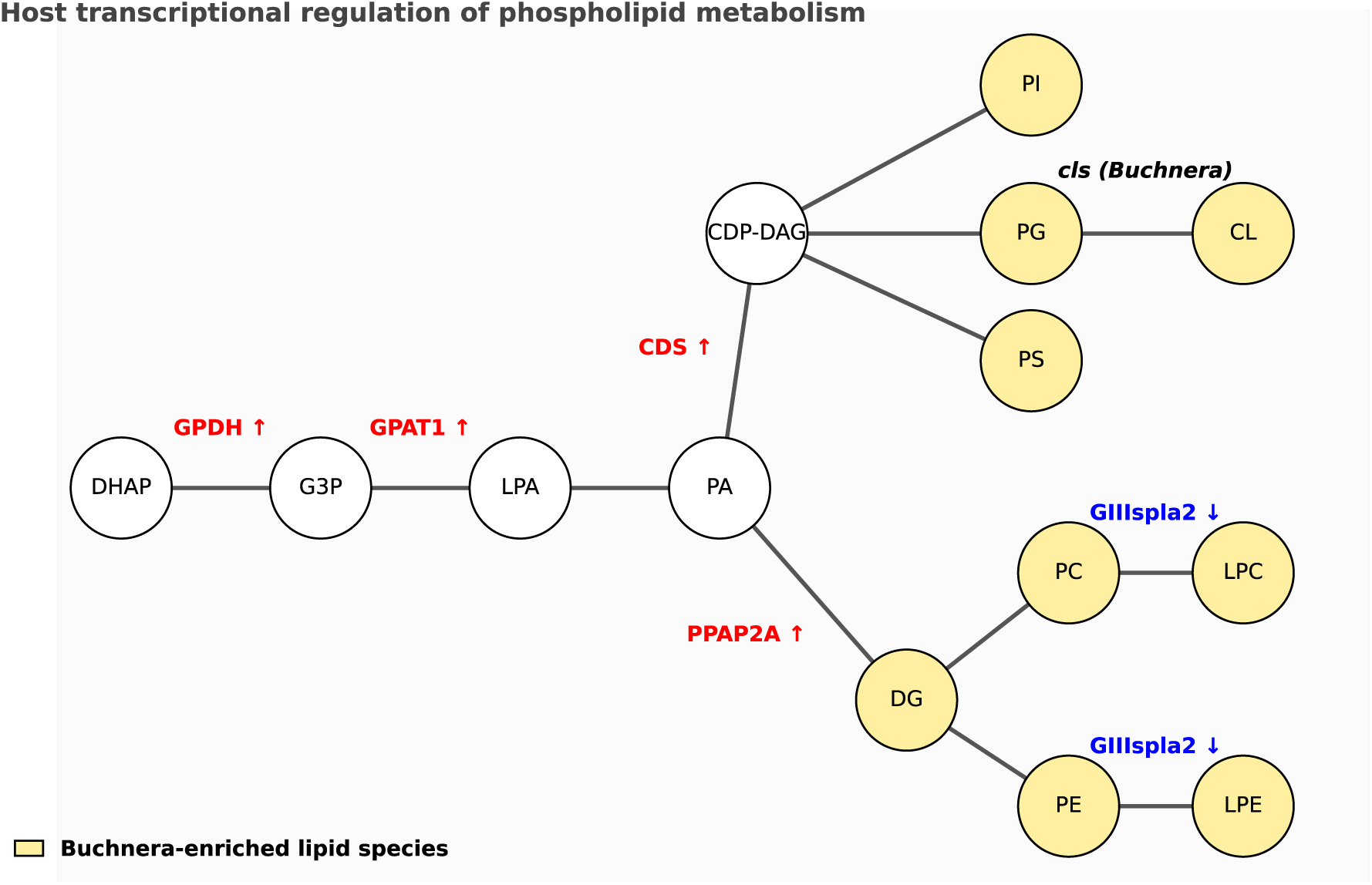
Integration of host transcriptomic and *Buchnera* lipidomic data links bacteriocyte phospholipid metabolism with *Buchnera* lipid profile Differentially expressed genes (DEGs) in viviparous *A. pisum* bacteriocytes compared with whole body tissues were integrated with *Buchnera* phospholipid profiles and mapped onto the glycerophospholipid metabolism pathway (KEGG: api00564). Genes upregulated in bacteriocytes are shown in red and downregulated genes in blue. The *Buchnera*-encoded *cls* gene (cardiolipin synthase) is indicated in italic black. The phospholipid biosynthetic pathway is shown schematically, with representative lipid classes indicated. Yellow highlighting denotes lipid classes for which one or more individual lipid species were significantly enriched in isolated *Buchnera* cells relative to bacteriocytes. Upstream host intermediates are shown in white nodes. The diagram shows that bacteriocytes transcriptionally elevate key steps leading from DHAP → G3P → PA and into CDP-DAG, DG and major phospholipids, while *Buchnera* retains only a late step (PG → CL via *cls*). Most phospholipids in *Buchnera* (PC, PE, PG, PI, PS) therefore appear to derive from host metabolism rather than *de novo* synthesis in the symbiont. Upstream precursors (e.g., DHAP, G3P, LPA, PA, CDP-DAG) were not detected in the lipidomic analysis. Lipid abbreviations: DHAP, dihydroxyacetone phosphate; G3P, glycerol-3-phosphate; LPA, lysophosphatidic acid; PA, phosphatidic acid; CDP-DAG, cytidine diphosphate-diacylglycerol; PI, phosphatidylinositol; PS, phosphatidylserine; PE, phosphatidylethanolamine; PG, phosphatidylglycerol; PC, phosphatidylcholine; CL, cardiolipin; LPC, lysophosphatidylcholine; LPE, lysophosphatidylethanolamine; DG, diacylglycerol.

### Functional perturbation links host lipid precursor metabolism to *Buchnera* maintenance

Importantly, the RNAi experiments provide functional evidence that host lipid precursor metabolism contributes to *Buchnera* maintenance. Knockdown of *AT-1* reduced *Buchnera* titer within 30 h and caused pronounced morphological abnormalities, whereas *GPDH* knockdown produced a comparable, although weaker and more condition-dependent, effect at a higher siRNA concentration and after longer treatment. These phenotypes extend the transcriptomic evidence beyond correlation and functionally link host pathways involved in acetyl-CoA trafficking and G3P production to *Buchnera* titer and cellular integrity. Although indirect effects of host metabolic perturbation cannot be excluded, the concordant effects of targeting two distinct components of host lipid precursor metabolism support an important role for host metabolic activity in sustaining *Buchnera*. Because membrane lipid synthesis is closely linked to bacterial growth and cell division [17], these findings raise the possibility that control over lipid precursor availability represents one of several host mechanisms regulating intracellular symbiont population dynamics. Indeed, insect hosts are thought to exert tight control over the density of obligate intracellular symbionts through multiple mechanisms [44] and lipid availability has been shown to limit endosymbiont proliferation in the *Drosophila–Spiroplasma* association [45]. However, unlike *Spiroplasma*, which resides in the host hemolymph and can directly assimilate circulating lipids in host, *Buchnera* is encapsulated in host-derived membrane compartments. Lipid availability to *Buchnera* may therefore depend on regulated provisioning across the symbiotic interface. Our results therefore raise the possibility that regulation of lipid availability represents an additional component of host control over symbiont abundance.

### Host sterol availability influences *Buchnera* abundance

The detection of cholesterol sulfate in purified *Buchnera* fractions provides another striking indication of host-associated lipid acquisition at the symbiotic interface. Sterols are generally absent from bacterial membranes, and neither partner can synthesize sterols de novo: aphids depend primarily on dietary sterols, whereas *Buchnera* lacks recognizable sterol biosynthetic genes [6], [20], [46]. Thus, the presence of a sterol derivative in the *Buchnera* fraction is most consistent with acquisition through the host environment. Importantly, cholesterol supplementation increased *Buchnera* titer after 48 h, providing an independent physiological line of evidence that sterol availability can influence symbiont maintenance. Exogenous cholesterol incorporation has been reported in bacteria such as *Bifidobacterium* longum, and host-derived cholesterol is directly incorporated by pathogenic intracellular bacteria, including *Ehrlichia chaffeensis* and *Anaplasma phagocytophilum* [47], [48], [49]. These examples indicate that acquisition of exogenous sterols by bacteria is biologically plausible. However, our experiments do not demonstrate that supplemented cholesterol, or the detected cholesterol sulfate, is directly incorporated into the *Buchnera* membrane; indirect effects through host physiology therefore remain possible. Nevertheless, the combined lipidomic and supplementation data support the possibility that host-mediated sterol availability influences *Buchnera* abundance.

### Unresolved mechanisms of host-to-symbiont lipid transfer

Despite the multiple lines of evidence for host-mediated lipid provisioning, the molecular mechanism by which lipids or their precursors reach *Buchnera* remains unresolved. Each *Buchnera* cell is enclosed by a host-derived symbiosomal membrane, creating an additional interface through which host metabolites must pass before reaching the bacterial cell envelope. Moreover, the highly reduced *Buchnera* genome lacks recognizable homologs of most characterized phospholipid transporters and phospholipases, providing little indication of how exogenous lipids are imported or remodeled [20]. Our data therefore do not distinguish whether *Buchnera* acquires intact phospholipids, lipid precursors that are subsequently remodeled, or a combination of both. Studies of intracellular bacteria have established several possible routes for host lipid acquisition, including vesicular and non-vesicular lipid transfer, providing useful mechanistic frameworks for the aphid–*Buchnera* system [49], [50], [51], [52], [53]. In aphids, bacteriocyte-enriched Rab GTPase expression, specialized transporters at the symbiosomal membrane, and the close association of the endoplasmic reticulum with *Buchnera*-containing regions are consistent with an active host–symbiont interface, although none has yet been directly linked to lipid transfer [50], [54], [55]. Thus, how host-derived lipids cross the symbiotic interface and reach the *Buchnera* membrane remains an important unresolved question.

### Host-supported membrane biogenesis in reduced endosymbionts

The implications of our findings likely extend beyond the aphid–*Buchnera* symbiosis. Extensive loss of fatty acid and phospholipid biosynthetic capacity has evolved repeatedly among obligate intracellular symbionts with highly reduced genomes, including *Portiera*, *Moranella*, *Mikella*, and *Carsonella* [20], [21], [28], [56], [57], [58], [59]. Remarkably, several of these symbionts show a pattern similar to that of *Buchnera*, in which most phospholipid biosynthetic genes have been lost while *cls* is retained, suggesting that dependence on host lipid metabolism can coexist with conservation of a minimal bacterial membrane function. These recurrent genomic patterns suggest that phospholipid auxotrophy and host-supported membrane biogenesis are not peculiarities of the aphid–*Buchnera* association, but may represent a common consequence of extreme genome reduction and long-term intracellular life. Cross-partner lipid utilization has also been suggested in other symbiotic systems; in the giant clam–Symbiodiniaceae association, algal betaine lipids are distributed throughout host tissues, consistent with their uptake and use by the host [60]. Our study provides experimental support for this broader model by linking the lipid composition of a highly reduced symbiont to host metabolic specialization and functional dependence on host lipid metabolism. More generally, these findings identify membrane biogenesis as an important, yet previously underappreciated, dimension of metabolic integration in obligate intracellular symbioses and suggest that host-mediated lipid provisioning may be a general mechanism contributing to the long-term stability of such associations.

## Materials and Method

### Aphid rearing

This study used the parthenogenetic clone of the pea aphid *Acyrthosiphon pisum* strain ApL (also known as Sap05Ms2), originally obtained from Sapporo, Hokkaido, Japan, and maintained at the National Institute for Basic Biology (Aichi, Japan) [61]. Viviparous aphids were maintained on broad bean plants (*Vicia faba* L.) at 16 °C with a photoperiod of 16/8 h light/dark. For experimental assays, four viviparous adult female aphids were randomly selected and placed in culture cups containing young broad bean plants. The adult females were allowed to reproduce for 48 h before removal. Offspring aphid nymphs were then reared for 48 h, after which second-instar (N2) nymphs were used for siRNA and cholesterol supplementation experiments.

### Sample preparation of aphid tissues and *Buchnera* cells for lipidome analysis

For whole-body samples, young adult pea aphids (10 d old) were placed in 1.5 ml centrifuge tubes, weighed, and immediately frozen in liquid nitrogen. For appendage samples, young adult aphids (10 d old) were frozen in liquid nitrogen for 15 min, vortexed to detach appendages, and the remaining core bodies were removed with forceps; collected appendages were weighed. For bacteriocyte samples, aphids (6–10 d old) were dissected in 500 µl extraction buffer (KCl 25 mM, MgCl_2_ 10mM, Tris-HCl 35 mM, sucrose 0.25 M) was prepared (KCl and MgCl_2_ from Sigma-Aldrich®, Burlington, MA, USA; Tris-HCl from Invitrogen™, Thermo Fisher Scientific, Waltham, MA, USA, sucrose from Fujifilm Wako Pure Chemical Corp., Osaka, JP) [62]. A 20 µl aliquot was mixed with 20 µl DAPI solution in 4% paraformaldehyde, fixed for 5 min, and 5 µl was mounted on glass slides with an 18 × 18 mm coverslip. Bacteriocytes were counted using an Olympus BX61 microscope (Olympus, Tokyo, Japan) (20 × objective) in three replicate counts per sample.

The *Buchnera* isolation method is modified from Houk & Mclean (1974) and Poliakov et al. (2001) [62], [63]. For *Buchnera* isolation, bacteriocytes from 6–10 d old aphids were homogenized in a Biomasher SP disposable homogenizer (Funakoshi Co., Ltd., Tokyo, JP) using the Nippi-sterilized Biomasher II. Homogenates were centrifuged at 600 × *g* for 10 min at 4 °C, and the pellet resuspended in 0.25 M sucrose. A Percoll gradient (45%, 27%, 18%, 9%) was prepared in 2 ml tubes, and the cell suspension was layered onto the 9% Percoll and centrifuged at 200 × g for 30 min at 4 °C. From the total gradient volume (1,800 µl), the upper 1,200 µl was removed, leaving the lower 600 µl. A total of 150 µl of suspension was then collected from the fraction corresponding to 600–450 µl from the bottom of the tube (see SI Appendix, Fig. S1A). This fraction was selected based on the optimal ratio of *Buchnera dnaK* to *A. pisum* mitochondrial cox1 gene copy numbers (see SI Appendix, S1B-C). The collected fraction was diluted five-fold with extraction buffer and centrifuged at 700 × *g* for 10 min at 4 °C. The pellet was resuspended in extraction buffer, filtered through a 5 µm membrane (Isopore™ 5 μm PC membrane TMTP01300) (Merck KGaA, Darmstadt, Germany), and centrifuged at 8,000 × g for 5 min at 4 °C. *Buchnera* cells were counted using a Thoma deep 0.1 mm hemocytometer (Erma Inc., Tokyo, Japan).

Isolate *Buchnera* cells were assessed using transmission electron microscopy (TEM) to verify cellular integrity and confirm removal of the host-derived symbiosomal membrane (see SI Appendix, Fig. S2). TEM images were obtained for *Buchnera* samples prior to and following Percoll density gradient centrifugation.

### Lipid extraction

Tissue samples for lipid extraction, including whole body, appendages, bacteriocytes, and *Buchnera* cells, were prepared as described above (see Supplementary Dataset S1). Lipid metabolites were extracted using a single-phase chloroform/methanol/water method (1:2:0.2, v/v/v) with internal standards for quantification.

For whole body and appendage samples, tissue was mixed with 100 µl methanol (MeOH), sonicated for 2 min, and dried completely. The dried pellet was resuspended in 150 µl MeOH and spiked with 5 µl of EquiSPLASH™ LIPIDOMIX® Quantitative Mass Spec Internal Standard (Avanti Polar Lipids), 0.32 µl each of 10 mM FA 16:0-d3 and FA 18:0-d3, and 1 µl of cardiolipin (CL) 18:2 d5 in chloroform. After 4 min of sonication, samples were incubated overnight at -30 °C, followed by addition of 20 µl water, vortexing, 10 min at room temperature, and centrifugation (2,000 × g, 10 min, 20 °C). A total of 200 µl of supernatant was transferred to liquid chromotagraphy (LC) vials. Injection volumes were 1 µl for positive and 3 µl for negative ion mode LC-MS/MS analysis.

For *Buchnera* cells, the same protocol was used: resuspension in 80 µl MeOH, internal standards comprising 2.5 µl EquiSPLASH™, 0.16 µl each of 10 mM FA 16:0-d3 and FA 18:0-d3, and 0.5 µl CL 18:2 d5. Ten µl water was added prior to centrifugation, and 150 µl of the supernatant was collected. Injection volumes were 3 µl for both positive and negative ion modes.

For bacteriocytes, the extraction followed the same solvent system, with 100 µl MeOH for initial sonication and resuspension. Internal standards comprised 2.5 µl EquiSPLASH™, 0.16 µl each of 10 mM FA 16:0-d3 and FA 18:0-d3, and 0.5 µl CL 18:2 d5. After overnight incubation at - 30 °C, samples were dried, resuspended in 200 µl 70% MeOH, and purified using C18 Spin Columns. Lipids were eluted with 160 µl isopropanol and transferred to LC vials. Injection volumes were 1 µl for positive and 3 µl for negative ion mode. All extractions were performed in triplicate per tissue type.

### Untargeted lipid profiling by liquid chromatography tandem mass spectrometry (LC-MS/MS)

Lipid extracts were analyzed using AQUITY UPLC I-class and H-class systems (Waters Corporation, MA, USA) coupled to a TripleTOF 6600 mass spectrometer (SCIEX, MA, USA). High-resolution data were acquired for both positive and negative ion modes. Lipid species were identified and quantified using MS-DIAL 4 software, applying standard parameters for peak detection, alignment, and annotation [64]. Lipid species were annotated according to the LIPID MAPS shorthand nomenclature using MS-DIAL output.

Oxidation or hydroxylation states were annotated according to MS-DIAL conventions, in which semicolon suffixes “;O”, “;O₂”, and “;O₃” indicate one, two, or three additional oxygen atoms on the acyl chain (e.g., hydroxyl or oxo groups). For consistency, these notations were reformatted as “(O)”, “(O2)”, and “(O3)” throughout this study. For example, the MS-DIAL output “FAHFA 20:0/24:0;O” is represented as “FAHFA 20:0/24:0(O)”. In figures and tables, the reformatted notation using parentheses is shown, while the original MS-DIAL output format (e.g., “ST 27:1;O;S”) is retained in the main text for consistency with raw data.

### Lipidomic data analysis

Lipidomic data were analyzed using the R package “lipidr” (v2.10.0) [65]. Lipid names were curated to ensure consistent parsing and class assignment, followed by lipid annotation using LIPID MAPS shorthand nomenclature. Signal intensities were normalized using probabilistic quotient normalization (PQN) and log-transformed prior to downstream analyses.

Global variation in lipid composition among sample types was assessed using unsupervised principal component analysis (PCA). Statistical differences in overall lipid profiles were evaluated using permutational multivariate analysis of variance (PERMANOVA) based on Euclidean distance matrices, with default permutation setting, implemented in the R package vegan (v2.6-4) [66]. To visualize lipid composition at the class level, normalized intensities were aggregated across lipid species within each lipid class and visualized using hierarchical clustering heatmaps with row-wise scaling.

Differential lipid abundance analysis was performed using lipidr to compare biologically relevant sample types, including *Buchnera* versus aphid bacteriocytes, *Buchnera* versus whole body, bacteriocytes versus whole body, and appendages versus whole body. For each lipid species, log₂ fold changes and statistical significance were calculated, and multiple testing correction was applied using the Benjamini–Hochberg (BH) procedure. Lipid set enrichment analysis (LSEA) was conducted on differential results to identify significantly enriched lipid classes, total acyl chain lengths, and degrees of unsaturation. Enriched lipid sets were defined using a false discovery rate (FDR) threshold of 0.05.

To focus on membrane-associated lipid dynamics, a subset of glycerophospholipid-related classes (including phosphatidic acid, diacylglycerol, major glycerophospholipids, lysophospholipids, and cardiolipin species) was extracted from the normalized dataset for targeted analyses. Within this phospholipid subset, PCA and PERMANOVA were repeated to assess compositional differences between *Buchnera* and aphid bacteriocytes. Differential abundance analysis was then performed specifically for the *Buchnera*–bacteriocyte comparison, and the most strongly enriched phospholipid species were identified based on adjusted *p*-values and fold changes.

To examine coordinated variation among phospholipid species, a correlation-based network was constructed using Spearman rank correlations calculated across samples using R package Hmisc (v5.1-1)[67]. Pairwise lipid–lipid correlations were computed using normalized phospholipid relative abundances, and only strong, positive correlations (Spearman’s ρ ≥ 0.85) with nominal statistical significance (BH-adjusted p < 0.05) were retained. The resulting adjacency matrix was used to generate an undirected weighted network, where nodes represent phospholipid species and edge weights correspond to correlation strength.

Phospholipid species were grouped into correlation-defined modules by average-linkage hierarchical clustering using a correlation-based distance metric, 1−∣ & ∣, derived from the matrix of pairwise Spearman correlation coefficients. Modules were identified from the resulting dendrogram using package dynamicTreeCut (v1.63-1) (deepSplit = 2, minimum cluster size = 5 lipid species)[68]. Module assignments were used to visualize coordinated patterns of phospholipid variation and to contextualize species-level enrichment patterns.

To relate network organization to differential lipid relative abundance, the 30 phospholipid species most strongly enriched in *Buchnera* and the 30 most strongly enriched in bacteriocytes (BH-adjusted p < 0.05 and ∣log_2_ FC∣ > 1) were mapped onto the correlation network. Network coordinates were generated using the Kamada-Kawai layout algorithm. Density-based spatial clustering of applications with noise (DBSCAN) was applied to the network coordinates of *Buchnera*- and bacteriocyte-enriched phospholipid species (eps = 0.6, minPts = 3) (dbscan v1.2.3) [69]. The resulting clusters were outlined using convex hulls to visualize the spatial distribution of differentially enriched lipids within the network.

Network visualization was performed using the R packages ggraph (v2.1.0), igraph (v1.5.1), and ggplot2 (v4.0.0), with density-based clustering used for visual grouping of enriched lipid species using ggplot2 [70], [71], [72].

### RNA extraction and library preparation

For transcriptomic profiling, total RNA was extracted from both bacteriocyte and whole-body samples of *A. pisum*. Approximately 10 d old aphids were dissected on 1x Phosphate buffered saline (PBS) (Gibco 10 x PBS pH 7.4, Thermo Fisher Scientific, MA, USA). For whole body samples, entire aphids of the same age were collected without dissection. Tissues were homogenized and lysed in the lysis buffer supplied with the NucleoSpin RNA XS kit (Takara Bio Inc., Shiga, Japan), and RNA extraction was performed according to the manufacturer’s instructions. RNA integrity was assessed using an Agilent Bioanalyzer 2100 (Agilent Technologies, CA, USA).

cDNA synthesis was carried out with the TaKaRa SMART-Seq HT kit (Takara Bio Inc., Shiga, Japan) following the manufacturer’s protocol. Amplified cDNA was purified using AMPure XP magnetic beads (Beckman Coulter Life Sciences d, IN, USA). Library tagmentation and indexing were performed with the Nextera XT DNA Library Preparation Kit (Illumina Inc., CA, USA) using cDNA. Library concentrations were determined by qPCR with the KAPA Library Quantification Kit (Roche, Basel, Switzerland) on an Applied Biosystems™ 7500 Real-Time PCR System (Thermo Fisher Scientific, Waltham, MA, USA) and normalized to 2 nM. Indexed libraries were pooled (5 µl per library) and sequenced on an Illumina NextSeq 550 platform (High Output mode, paired-end 150 bp reads).

### Transcriptomic data processing and analysis

Raw paired-end RNA-seq reads were processed to remove adapter sequences using Trimmomatic (v0.39) [73]. Read quality was assessed with FastQC (v0.11.9), and summary reports were aggregated using MultiQC (v1.22.3) [74], [75]. Reference genomes for *A. pisum* (GenBank accession number: GCF_005508785.2) and *B. aphidicola* strain APS (GenBank accession number: NC_002528.1) were used with the bbsplit.sh script from BBMap (v39.06) to map reads simultaneously to both references [6], [42]. rRNA sequences were removed using RiboDetector (v0.3.1) [76]. The cleaned *A. pisum* reads were quantified in mapping-based mode with Salmon (v1.10.2) [77]. RNA-seq quality control metrics, including read counts before and after trimming, percentage of reads passing filters, rRNA content, mapping rates to the pea aphid genome, and key FastQC statistics (GC content, duplication rate, and read length), are provided in Supplementary Dataset S1.

Transcript count data from Salmon were imported into R using tximport (v1.24.0) [78]. Differential expression analysis was performed with the quasi-likelihood F-test (QLF) implemented in edgeR (v3.38.4) [79]. Genes were considered differentially expressed (DEGs) if they satisfied |log₂FC| > 1 and false discovery rate (FDR) < 0.05. Raw *p*-values are also adjusted by default using the BH method to control the FDR.

Gene Ontology (GO) annotation of DEGs was performed using the org.Apisum.eg.db package (v0.1) released by AphidBase [80]. GO enrichment analysis was conducted with goseq (v1.56.0), and hierarchical relationships between parent and child GO terms were resolved using GofuncR (v1.24.0) [81], [82]. KEGG IDs for significant DEGs were retrieved via org.Apisum.eg.db, and pathway mapping was performed using Pathview (v1.36.1) [19], [83]. The acetyl-CoA transporter 1 inparalogs search was conducted using eggNOG database [27].

### Integration of transcriptomic and lipidomic data

Significant DEGs from transcriptomic analysis and enriched *Buchnera* lipid species identified from the lipidome differential analysis were jointly mapped onto a single KEGG pathway (KEGG database: api00056 for *A. pisum* glycerophospholipid metabolism) (refer to Fig. S11, SI Appendix for original integrated map). Transcriptomic integration was performed at the enzyme level: DEGs were mapped to KEGG enzyme entries using their ENTREZ gene identifiers, and where multiple gene models or paralogs corresponded to the same enzyme, the gene exhibiting the largest absolute log₂ fold change was retained to represent that enzymatic step (Supplementary Dataset S1). Enriched lipid species were aggregated by lipid class, and both mean and maximum log₂ fold change values were calculated for each class (Supplementary Dataset S1). Lipid classes were then manually annotated with corresponding KEGG compound identifiers associated with the *A. pisum* glycerophospholipid metabolism pathway (api00564). The R package Pathview (v1.36.1) was then used to visualize both gene- and lipid-level changes within the pathway of interest [83]. A schematic overview of the RNA-seq preprocessing, downstream analyses, and transcriptome–lipidome integration pipeline is provided in Fig. S10 (SI Appendix) [84].

*Buchnera* glycerophospholipid metabolism (KEGG database: buc00564) and fatty acid biosynthesis pathway (KEGG database: buc00061) (refer to Fig. S7A-B, SI Appendix) was reconstructed based on KEGG pathway [19].

### siRNA design, synthesis and administration

Two aphid genes were targeted: glycerol-3-phosphate dehydrogenase (*LOC100160658/GPDH;* SiRNA designation: ApB744) and acetyl-CoA transporter 1 (*LOC100159262/AT-1*; siRNA designation: ApW584). Two independent siRNAs were designed from the target transcript sequences and synthesized by Nippon Gene Co., Ltd. (Toyama, Japan) (see Supplementary Dataset S1). A universal negative control siRNA (Nippon Gene) with no homology to known eukaryotic sequences was used as reference.

siRNAs were diluted to final working concentrations of 12 µM or 20 µM in RNAse free water, and was delivered into second-instar aphid nymphs by glass capillary microinjection. Details on microinjection were detailed in Tan & Shigenobu (2024). Based on the previously determined injection volume under these conditions (0.2805 ± 0.0462 µL), the estimated siRNA doses were approximately 3.4 and 5.6 pmol per aphid for the 12 and 20 µM treatments, respectively. Injected aphids were maintained on broad bean seedlings at 16 °C (16:8 h light:dark) for 24–50 h. Each data point represents one biological replicate consisting of three pooled aphid nymphs.

### Gene expression analysis

RT-qPCR followed the protocol of Tan & Shigenobu (2025) [35]. Briefly, total RNA was extracted from pools of three aphids and gene expression was assessed using the Takara One Step TB Green PrimeScript™ RT-PCR kit (Takara Bio Inc., Shiga, JP) and the Roche LightCycler® 96 real-time PCR instrument (Roche Applied Science, Penzberg, DE). Both actin and elongation factor 1α (*elfA*) served as reference genes [50]. Target gene expression was assessed with primers specific to LOC100160658 and LOC100159262 using relative quantification method. Each biological replicate consisting of three pooled aphid nymphs. Log-transformation of gene expression values was performed prior to statistical analyses.

### Quantification of *Buchnera* titer following siRNA treatment

*Buchnera* abundance in siRNA-treated aphids was quantified by qPCR targeting *dnaK* (primers BuchDnaK_F1018 and BuchDnaK_R1142) and normalized to aphid *rpL7* (primers rpL7_F and rpL7_R), following the protocol as described in our previous publication (refer to Supplementary Dataset S1) [35], [50]. Each data point represents one biological replicate consisting of three pooled aphid nymphs. Three injected aphid nymphs were homogenized with a Nippi-sterilized Biomasher II (Funakoshi Co., Ltd., Tokyo, JP) in a tube containing 100 μl Buffer A (10 mM Tris–HCl, 1 mM EDTA, 25 mM NaCl in UltraPure distilled water) with 0.4 mg/ml Proteinase K (QIAGEN N.V., Venlo, NL). Homogenized samples were incubated at 37 °C for 1.5 h followed by 98 °C for 2 min. Relative *Buchnera* titers were log-transformed prior to statistical analyses.

### Cholesterol supplementation

A cholesterol stock solution (10 mg/mL) was prepared by dissolving 11 mg cholesterol (Wako, Osaka, Japan) in 1.0 mL methanol (99.8%; Fujifilm Wako Pure Chemical Corp., Osaka, Japan) and 0.1 mL chloroform (99.5%; Wako) at 37 °C for 10 min. The mixture were vortex vigorously for 1 min thrice during the incubation. A working cholesterol solution (1 µg/µL) was prepared by mixing 100 µL of the stock solution with 890 µL of Ptx (1× PBS, supplemented with 0.2% Triton X-100; Sigma-Aldrich, Burlington, MA, USA) and an additional 10 µL chloroform, yielding a final volume of 1.0 mL (1 µg/µL cholesterol, 9.1 % methanol, 1.9 % chloroform and 89.0 % Ptx, v/v).

Aphids were injected with the cholesterol working solution using glass capillary microinjection as described in Tan & Shigenobu (2024). Two controls were included: (i) a vehicle control (NegControl) containing the full diluent used for cholesterol delivery (Ptx, methanol and chloroform at concentrations equivalent to those in the cholesterol working solution), and (ii) a Ptx control. The estimated cholesterol dose was approximately 0.28 µg per aphid. Following treatment, aphids were maintained on broad bean seedlings at 16 °C under a 16:8 h light:dark cycle and collected for *Buchnera* titer quantification at 24 h or 48 h post-treatment. Each biological replicate comprised of a single treated aphid nymphs.

### Statistical analysis

All analyses were performed in R (v4.2.3) using relevant packages, including devtools (v2.4.5), DescTools (v0.99.50), and dplyr (v1.1.4) [85], [86], [87], [88]. Data visualization was conducted in R with ggpubr (v0.6.0) and smplot2 (v0.1.0) for raincloud plots [89], [90]. Data were first evaluated for normality and homoscedasticity using the Shapiro–Wilk and Levene’s tests, respectively. Pairwise comparisons of gene expression were assessed using Welch’s one-tailed *t*-test. *Buchnera* titers across multiple treatments were analyzed by one-way ANOVA with Dunnett’s or LSD post hoc tests, as appropriate. All statistical analyses were conducted with a significance threshold of α = 0.05. Asterisks indicate statistical significance: *p* < 0.05 (*), *p* < 0.01 (**); NS, not significant. Statistical details, including test type, mean ± SD, *p*-values, and sample size (*n*), are reported in figure legend.

## Author Contributions

K.X.Y.T. and S.S. conceived and designed the study. K.X.Y.T. performed the lipidomic and transcriptomic analyses, conducted RNA interference and cholesterol supplementation experiments, carried out statistical analyses, and wrote the manuscript. D.M. provided suggestions for lipidome analyses and contributed to manuscript editing. M.A. and A.H. performed lipid extraction, LC–MS/MS experiment and lipid annotation. T.N. and N.M. prepared RNA-seq libraries for transcriptome analysis. M.K. assisted in transmission electron microscopy (TEM) sample preparation. K.X.Y.T., D.M., and S.S. revised the manuscript. S.S. provided overall project supervision, secured funding, and contributed to manuscript editing. All authors discussed the results and approved the final version of the manuscript.

## Competing Interest Statement

The authors declare no competing interests. Classification: Biological Sciences, Evolution

### Data, Materials, and Software Availability

The raw reads for RNA sequencing are available at the National Center for Biotechnology Information Short Read Archive, under BioProject: PRJDB42692

All custom scripts used for statistical analyses have been deposited at

GitHub: https://github.com/ktxy2004/Buchnera_Apisum_lipidome_transcriptome

Figshare: https://doi.org/10.6084/m9.figshare.c.8784204

## Acknowledgments

We would like to express our earnest gratitude toward all members of the Laboratory of Evolutionary Genomics for their advice and support. We would like to thank M. Suzuki and S. Shaqiri for their assistance in RNAseq sample preparation. We thank the Optics and Imaging Facility, NIBB Trans-Scale Biology Center, for their technical support. We appreciate the support provided by the Section of Electron Microscopy in National Institute for Physiological Sciences. We thank Dr. H. Salem and members of Mutualisms Research Group, MPI for Biology for their support and encouragement. This work was funded by the Japan Society for the Promotion of Science (JSPS) KAKENHI Grants-in-Aid for Scientific Research (A) (No. JP20H00478) to S. S., MEXT Scholarship and Grant-in-Aid for Research Activity Start-up (No. 25K23703) to K. T., and EMBO fellowship (ALTF 772-2023) to D.M. This work was also supported by the JST-ERATO “ARITA Lipidome Atlas Project” under grant number JPMJER2101 (M.A.), AMED Moonshot Research and Development Program JP22zf0127007 (M.A.), AMED NeDDTrim (21ae0121036 to M.A.).

## Supporting Information

Appendix 01

Dataset S01

## Supporting Information for

### Identification of the Percoll layer with an optimized ratio of *Buchnera* to mitochondria

To identify the Percoll density gradient fraction with the highest *Buchnera* purity and minimal host mitochondrial contamination, we compared the ratio of *Buchnera* to *Acyrthosiphon pisum* mitochondria in samples aspirated from different gradient layers (Fig. S1B-C). DNA was extracted from each fraction using the method described in method. Quantitative PCR (qPCR) was performed targeting the *Buchnera dnaK* gene with primers BuchDnaK_F1018 and BuchDnaK_R1142, and the *A. pisum* mitochondrial *cox1* gene with primers ApMito_F895 and ApMito_R1055 (see Supplementary Dataset S1). Standard curves were constructed for each target to quantify gene copy numbers.

The ratio of log-transformed *dnaK* to *cox1* copy number was calculated for each fraction. Ratios were compared among Percoll layers by one-way ANOVA at a significance threshold of α = 0.05. All analyses were performed in R (v4.2.3) using the packages devtools (v2.4.5), DescTools (v0.99.50), and dplyr (v1.1.4) [1], [2], [3], [4]. Boxplots were generated with ggplot2 (v4.0.0) [5]. Schematic illustrations of the workflow were prepared using draw.io (v15.5.2) [6].

**Fig. S1.**
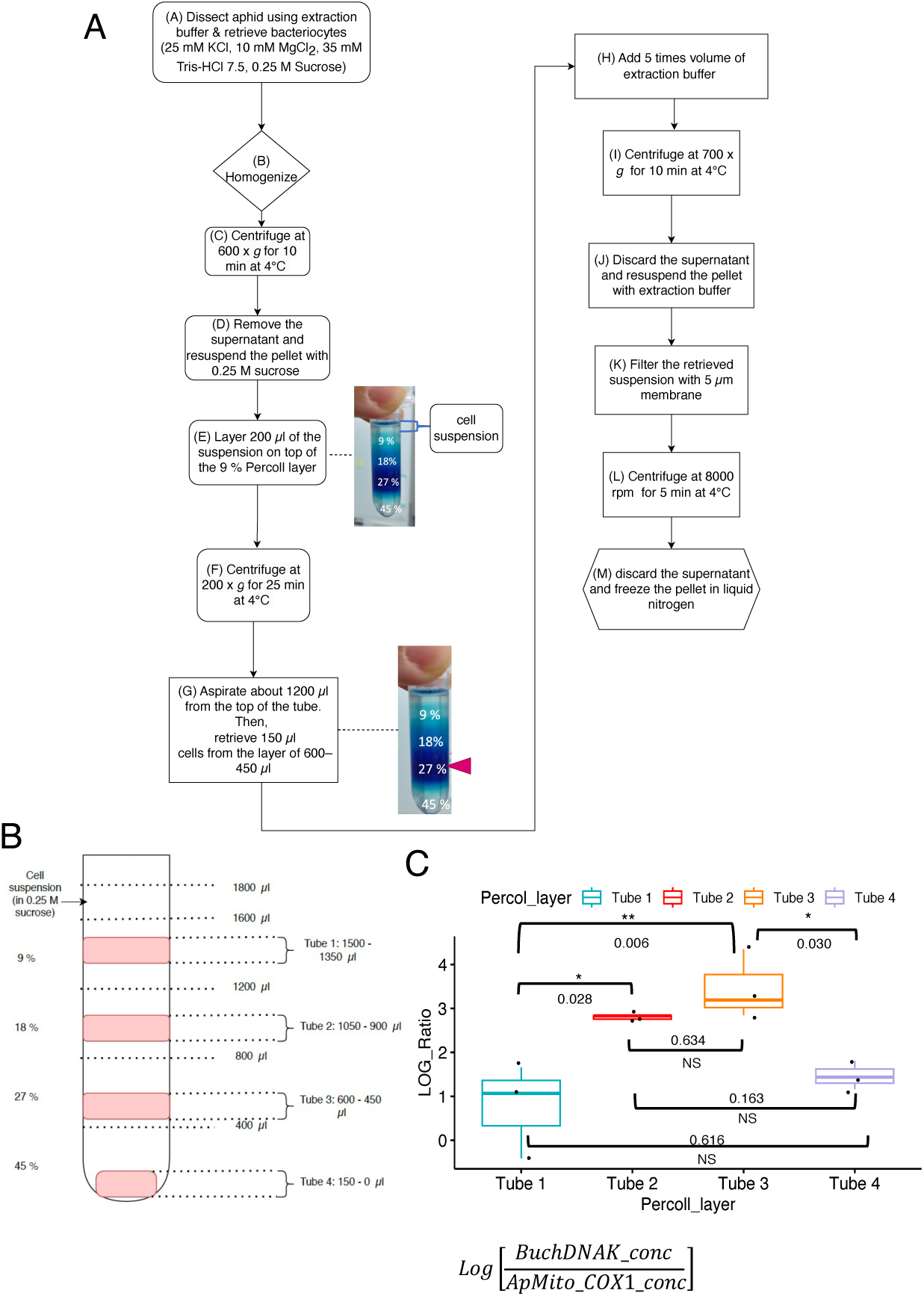
Workflow and optimization of *Buchnera* isolation using *Percoll* density gradient centrifugation. **(A)** Schematic overview depicting the isolation of *Buchnera* cells from dissected bacteriocytes via Percoll density gradient centrifugation. **(B)** Optimization of the Percoll gradient for effective *Buchnera* purification. Liquids aspirated from distinct gradient fractions (labeled Tube 1–4) were assessed by qPCR targeting *Buchnera dnaK* and *A. pisum* mitochondrial *cox1* genes to determine bacterial purity and mitochondrial contamination. **(C)** Log-transformed ratios of *Buchnera dnaK* to *A. pisum* mitochondria *cox1* copy numbers were analyzed using ANOVA (*p* = 0.005 < 0.05, Tukey’s HSD post hoc test), where each black dot represents an independent replicate from a single gradient fraction. This optimization confirmed minimal mitochondrial contamination, ensuring high-quality *Buchnera* preparations suitable for downstream lipid analysis.

### Transmission Electron Microscopy (TEM) for ultrastructural verification of *Buchnera* cells

#### prior to and after Percoll density gradient centrifugation

Electromircographs of *Buchnera* cells were presented in Figure S2. Aphid nymphs (6–10 d old) were dissected to extract bacteriocytes, which contain the obligate symbiont *Buchnera aphidicola*. *Buchnera* cells were isolated from bacteriocytes using Percoll density gradient centrifugation as outlined in Fig. S1. Samples were prepared from two steps: (i) prior to Percoll centrifugation (Fig. S1B) and (ii) following Percoll centrifugation (Fig. S1G). A double-fixation protocol was applied [7]. Pre-fixation was carried out in 4% paraformaldehyde and 1% glutaraldehyde in extraction buffer (25 mM KCl, 10 mM MgCl₂, 35 mM Tris-HCl, 0.25 M sucrose; KCl and MgCl₂ Sigma-Aldrich®, Burlington, MA, USA; Tris-HCl from Invitrogen™, Thermo Fisher Scientific, Waltham, MA, USA; sucrose from Fujifilm Wako Pure Chemical Corp., Osaka, Japan) for 2 h at room temperature, followed by overnight fixation at 4 °C. Samples were washed once with extraction buffer before post-fixation in a fresh solution of 2% osmium tetroxide in 0.1 M PBS for 2 h at room temperature.

After fixation, samples were dehydrated in graded ethanol, followed by propylene oxide and infiltrated with propylene oxide-Epon (Epon 812 resin; TAAB Laboratories, Aldermaston, UK) solution (propylene oxide-Epon resin, 1:1 [v/v]) overnight. Ultrathin sections were prepared using a Leica UCT Ultracut ultramicrotome with a glass knife and mounted on nickel grids. The sections were observed using a JEOL JEM-1010 transmission electron microscope (JEOL, Tokyo, Japan) equipped with a 2k × 2k CCD camera.

### Super-resolution imaging of aphid bacteriocytes and *Buchnera* cells following siRNA and cholesterol supplementation treatments

Super-resolution laser scanning confocal microscopy (Zeiss LSM 980 with Airyscan 2 in Airyscan SR mode) was used to visualize morphological effects at 46 h post-treatment. Aphid bacteriocytes were dissected and fixed as described in Tan & Shigenobu (2024), then stained with DAPI (4,6-diamidino-2-phenylindole; Dojindo Laboratories, Kumamoto, Japan) to visualize DNA [8]. Images were acquired as single optical planes using a Plan-Apochromat 63×/1.40 oil immersion lens, with fluorescence and DIC channels processed in ZEN Blue (v3.4.91) in default 2D super-resolution mode. Channel settings were λ_ex = 473 nm (0.2% laser power, detector gain = 750 V; detection = 380–735 nm) for DAPI, and λ_ex = 488 nm (0.2% laser power, detector gain = 350 V; detection = 300–900 nm) for DIC. Linear unmixing (Automatic Component Extraction) was applied to remove autofluorescence.

**Fig S2.**
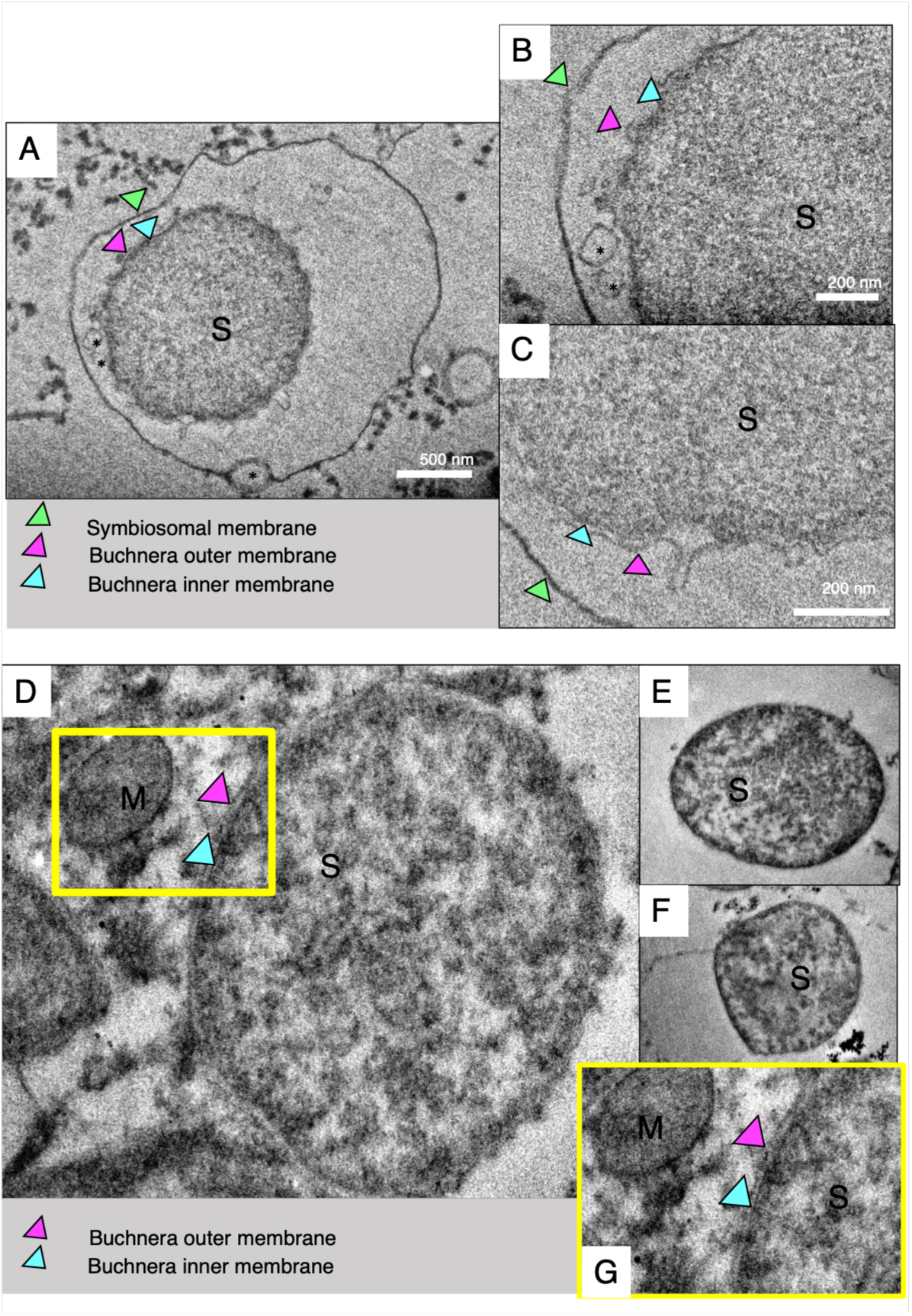
Validation of *Buchnera* cell isolation by Transmission Electron Microscopy (TEM). TEM was utilized to confirm the physical integrity and localization of *Buchnera* cells before and after Percoll gradient centrifugation. (**A-C**) Before Percoll gradient treatment (step shown in Fig S1B), isolated *Buchnera* cells retained intact host-derived symbiosomal membranes (green arrowhead). The clearly distinguishable bacterial outer (magenta arrowhead) and inner (cyan arrowhead) membranes are visible, as are vesicular structures (*). (**D-G**) Following Percoll gradient centrifugation (step shown in Fig S1G), the symbiosomal membrane was effectively removed. The outer (magenta arrowhead) and inner (cyan arrowhead) bacterial membranes remain structurally intact. (* represents vesicles; S: *Buchnera* cell; M: mitochondrion)

**Fig S3.**
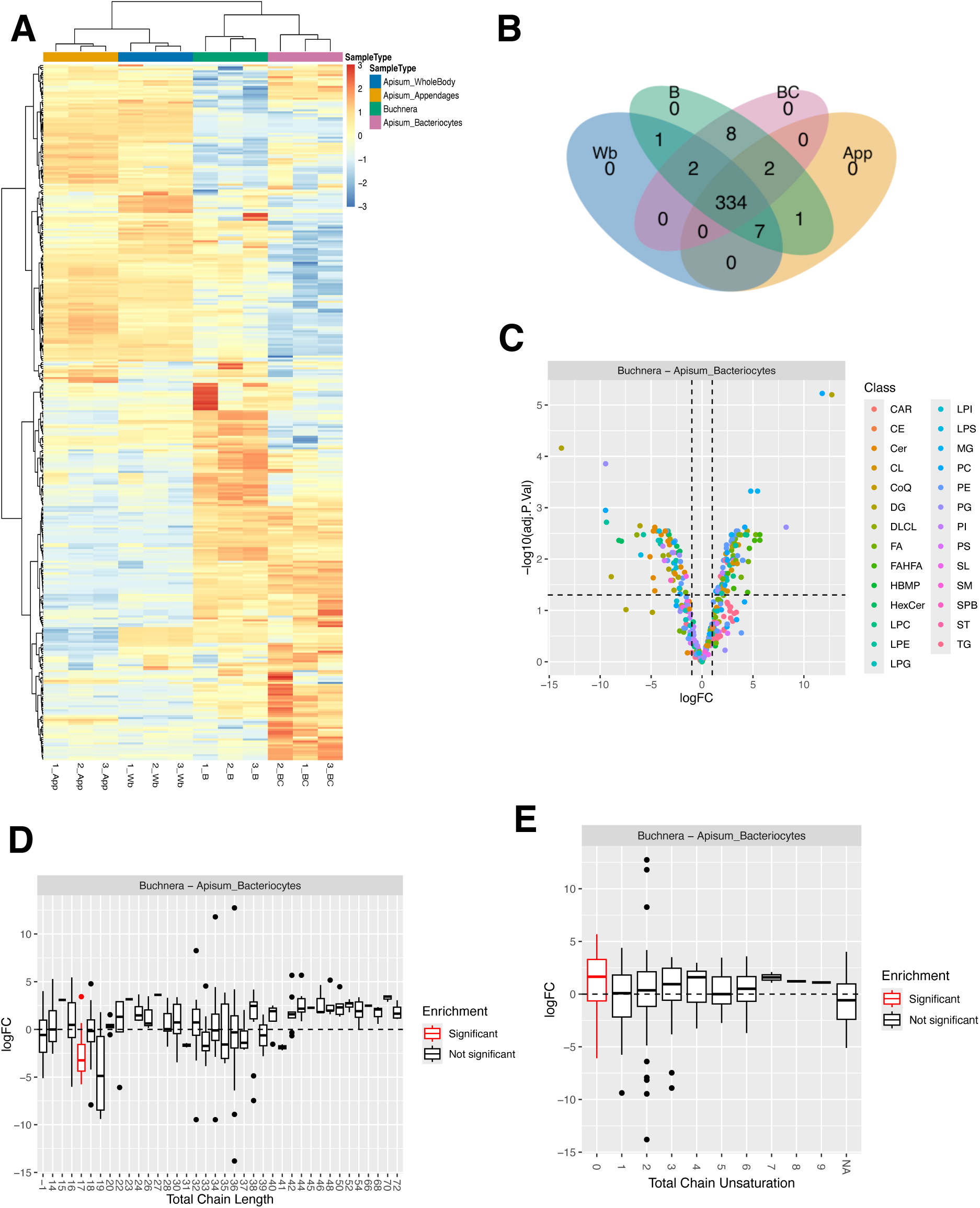
Differential lipidomic features of *Buchnera* compared to *A. pisum* bacteriocytes. **(A)** Heatmap showing clustering of all detected lipid species across aphid whole body, appendages, bacteriocytes and isolated *Buchnera* samples. Rows represent lipid species and columns represent individual biological replicates, colour scale denotes normalized, log-transformed and called by row. Colour scale denotes relative abundance (blue = low; red = high). Clustering was performed using Euclidean distance and Ward’s D^2^ method. Distinct clustering was observed in different tissue and cell types. (**B)** Venn diagram showing lipids features in whole body (Wb), appendages (App), bacteriocytes (BC), and isolated *Buchnera* (B). **(C)** Volcano plot showing differentially abundance of lipid between Buchnera and host bacteriocytes. Each dot represents one lipid species. Vertical dashed lines mark logFC = ±1, and the horizontal dashed line indicates adjusted *p* = 0.05 (−log₁₀ adj. *p* = 1.3). (**D)** Lipid-set enrichment analysis (LSEA) based on total acyl-chain length. Boxplots show log₂ fold-change distributions for each chain-length group; significantly enriched chain lengths are outlined in red. **(E)** LSEA based on total chain unsaturation. Boxplots represent distributions of log₂ fold-change by degree of unsaturation; significant enrichment is shown in red (FDR < 0.05). Lipid-class abbreviations: CAR, acylcarnitine; CE, cholesteryl ester; Cer, ceramide; CL, cardiolipin; DG, diacylglycerol; DLCL, dilysocardiolipin; FA, fatty acid; FAHFA, fatty acid ester of hydroxyl fatty acid; HBMP, hemibismonoacylglycerophosphate; HexCer, hexosylceramide; LPC, lysophosphatidylcholine; LPE, lysophosphatidylethanolamine; LPG, lysophosphatidylglycerol; LPI, lysophosphatidylinositol; LPS, lysophosphatidylserine; MG, monoacylglycerol; PC, phosphatidylcholine; PE, phosphatidylethanolamine; PG, phosphatidylglycerol; PI, phosphatidylinositol; PS, phosphatidylserine; SM, sphingomyelin; SPB, sphingoid base; TG, triacylglycerol, ST, steroid conjugates, SL, sulfonolipid.

**Fig. S4.**
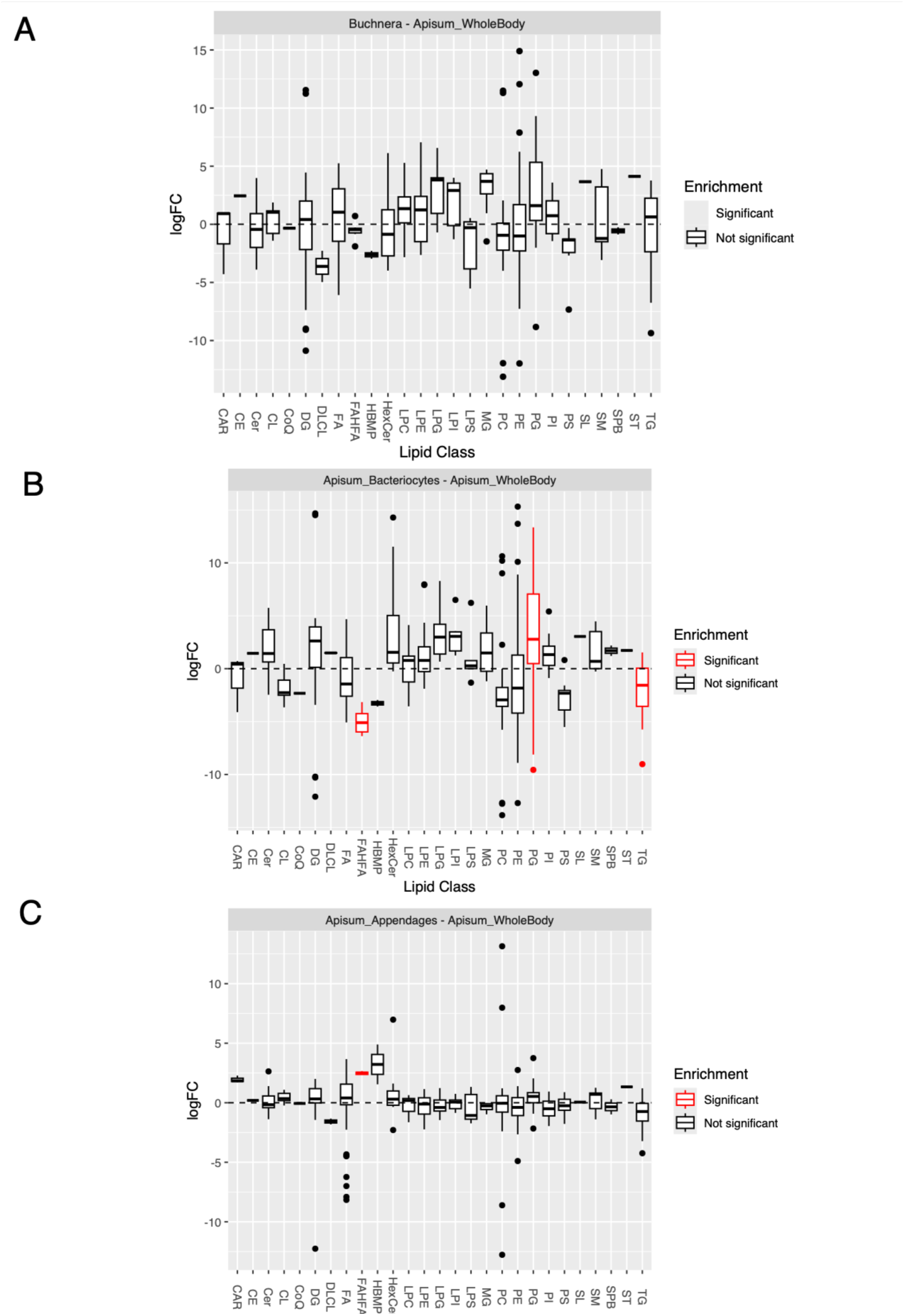
LSEA showing lipid class enrichment or depletion in *Buchnera* and aphid tissues compared to wholebody extracts of *A. pisum*. Lipid Set Enrichment Analysis (LSEA) showing classes significantly enriched or depleted compared with the aphid whole-body lipidome (FDR < 0.05). **(A)** No lipid class was significantly enriched in the Buchnera lipidome relative to the whole body. **(B)** In A. pisum bacteriocytes, FAHFA and TG were significantly depleted, whereas PG was significantly enriched. **(C)** In aphid appendages, only FAHFA was significantly enriched. Boxplots display the distribution of log₂ fold-change values for individual lipid species within each class. Red-coloured boxplot show either significant enrichment or depletion based on the log₂ FC.

**Fig. S5.**
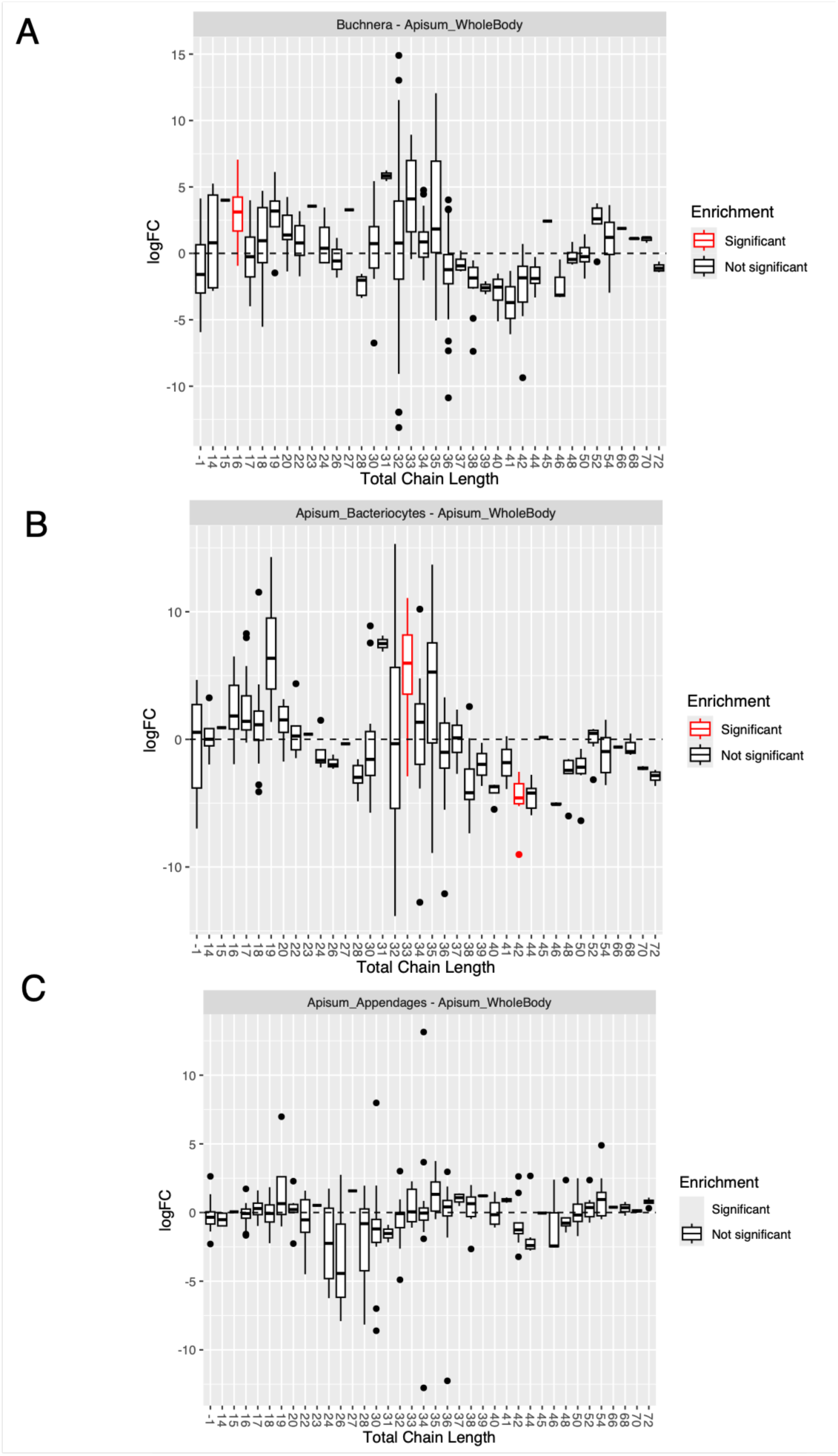
LSEA of total lipid acyl chain length in *Buchnera* and aphid tissues compared to whole-body lipid profiles of *A. pisum*. Total carbon numbers in lipid acyl chains and their corresponding log_2_ FC are shown. Significantly enriched chain lengths are marked in red (FDR < 0.05). (**A**) In the *Buchnera* lipidome, chain lengths of 16-carbon was significantly enriched. (**B**) In bacteriocytes, chain length of 33 showed significant enrichment while chain length of 42 was significantly depleted. (**C**) No specific total chain length was enriched in the lipidome of aphid appendages. Values at −1 represent lipids without parsed chain information. Red-coloured boxplot show either significant enrichment or depletion based on the log₂ FC.

**Fig. S6.**
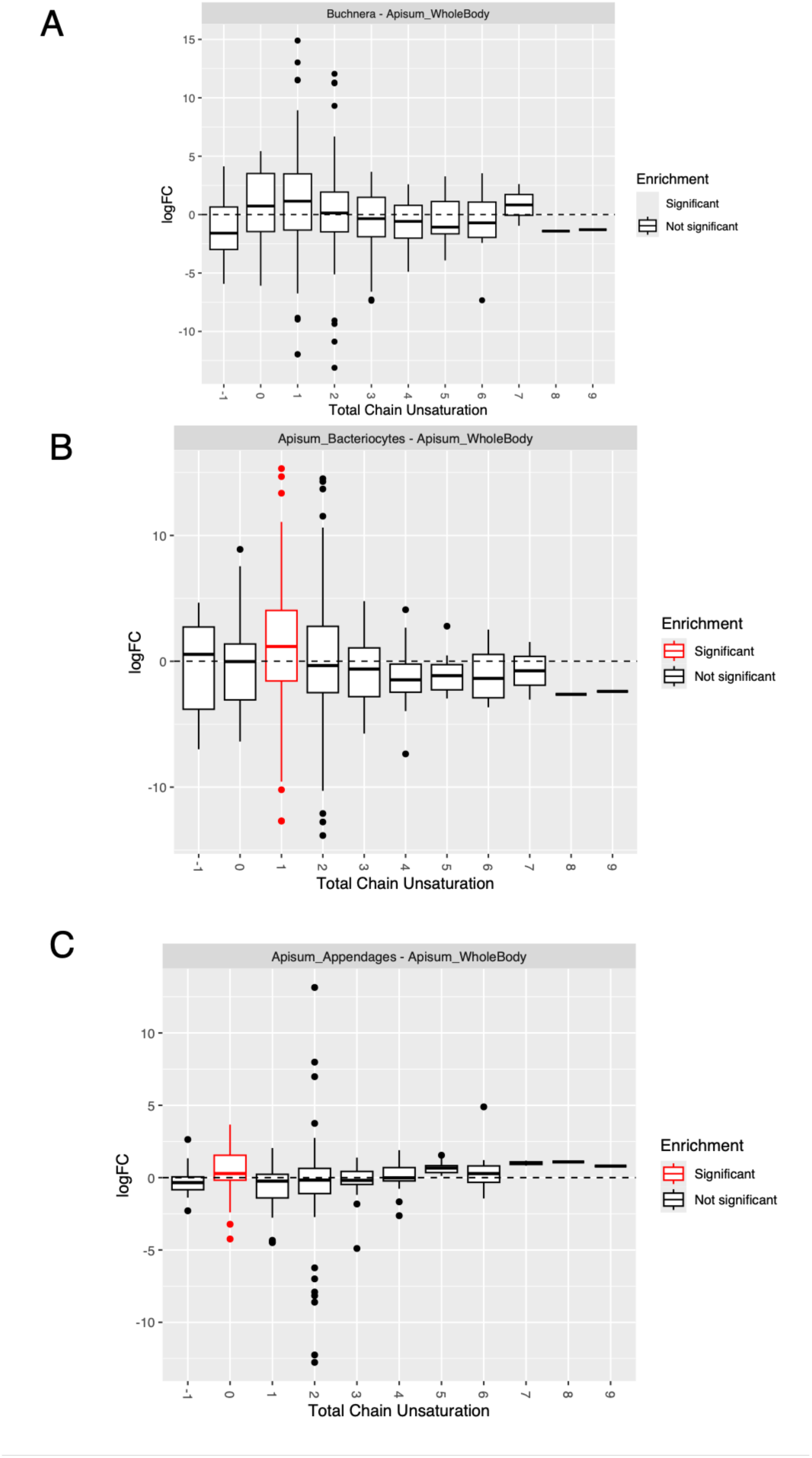
LSEA of total chain unsaturation in *Buchnera* and aphid tissues compared to *A. pisum* whole body. Total chain unsaturation values and their corresponding log_2_ FC are shown. Significantly enriched levels of unsaturation are marked in red (FDR < 0.05). (**A**) No significantly enriched total chain unsaturation was detected in the *Buchnera* lipidome. (**B**) Lipid chains with a single unsaturation were significantly enriched in bacteriocytes. (**C**) Saturated lipid chains (zero unsaturation) were significantly enriched in the lipidome of aphid appendages. Values at −1 represent lipids without parsed chain information.

**Fig. S7.**
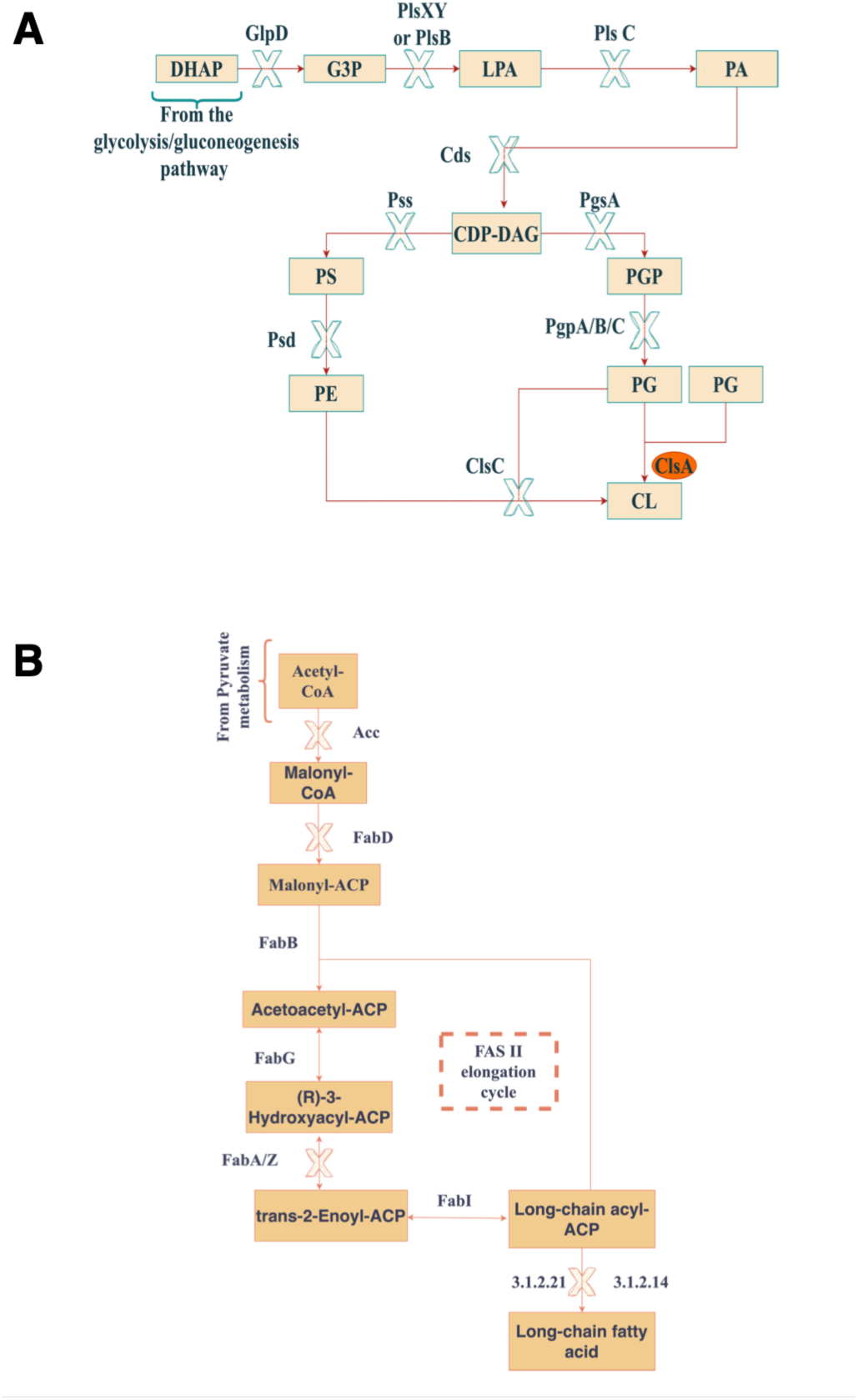
Loss of phospholipid and fatty-acid biosynthetic capacity in *Buchnera*. **(A)** Simplified schematic of the phospholipid biosynthesis pathway. Enzymes and intermediates absent from the *Buchnera* genome are indicated with “X” marks. *Buchnera* retains only a single phospholipid-related gene, *clsA*, encoding cardiolipin synthase (orange), which catalyzes the final step converting phosphatidylglycerol (PG) to cardiolipin (CL). All other upstream enzymes required to produce phosphatidic acid (PA), cytidine diphosphate-diacylglycerol (CDP-DAG), phosphatidylserine (PS), phosphatidylethanolamine (PE), and phosphatidylglycerol (PG) are missing, indicating a loss of autonomous phospholipid synthesis. Adapted from Shigenobu et al. (2000), Sohlenkamp & Geiger (2016), and KEGG pathway [9], [10], [11]. **(B)** Fatty acid synthesis (FAS II) pathway showing the conversion of acetyl-CoA to long-chain fatty acids. Enzymes crossed in blue are absent from the *Buchnera* genome. The extensive loss of key FAS II enzymes such as FabD and FabA/Z, demonstrates that *Buchnera* cannot produce fatty-acid precursors required for phospholipid assembly and must therefore rely on host-supplied lipids to maintain membrane biogenesis. Adapted from KEGG pathway [11].

**Fig. S8.**
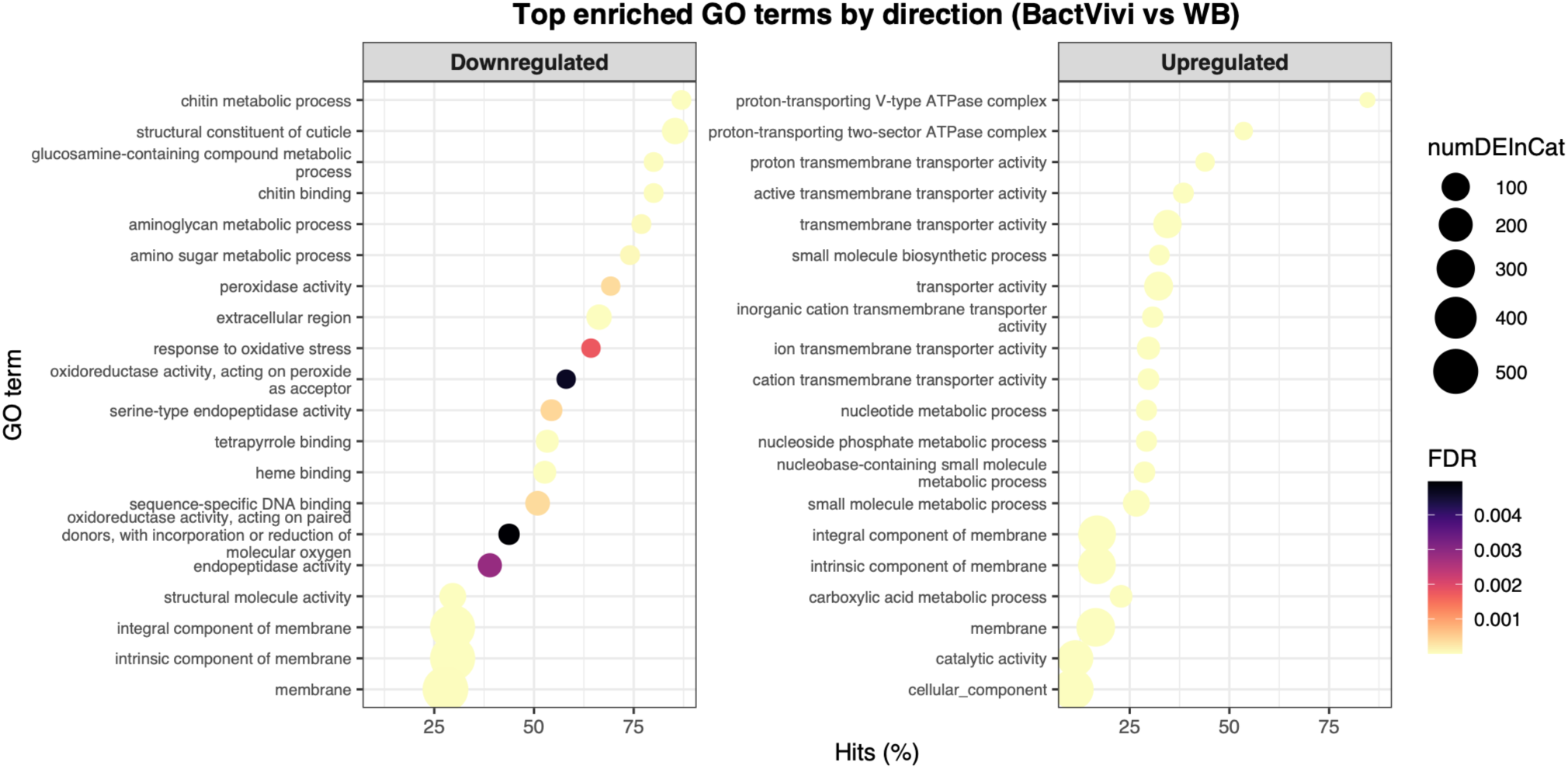
GO enrichment analyses of up- and downregulated genes in viviparous bacteriocytes compared with whole body aphids. GO analysis was performed separately on significantly up- and downregulated genes (FDR < 0.05, |log₂FC| > 1) to identify overrepresented Gene Ontology (GO) terms while correcting for gene length bias. Bubble plots show the top 20 enriched biological processes for each direction. Dot size represents the number of differentially expressed genes per GO term, and color intensity corresponds to the adjusted FDR. Downregulated genes were mainly associated with cuticle, oxidative stress response, and extracellular region functions, whereas upregulated genes were enriched for membrane-associated and transporter activities, including transmembrane transport and metabolic processes. These results reinforce the transcriptional specialization of bacteriocytes in mediating molecular exchange and metabolic regulation for symbiosis.

**Fig. S9:**
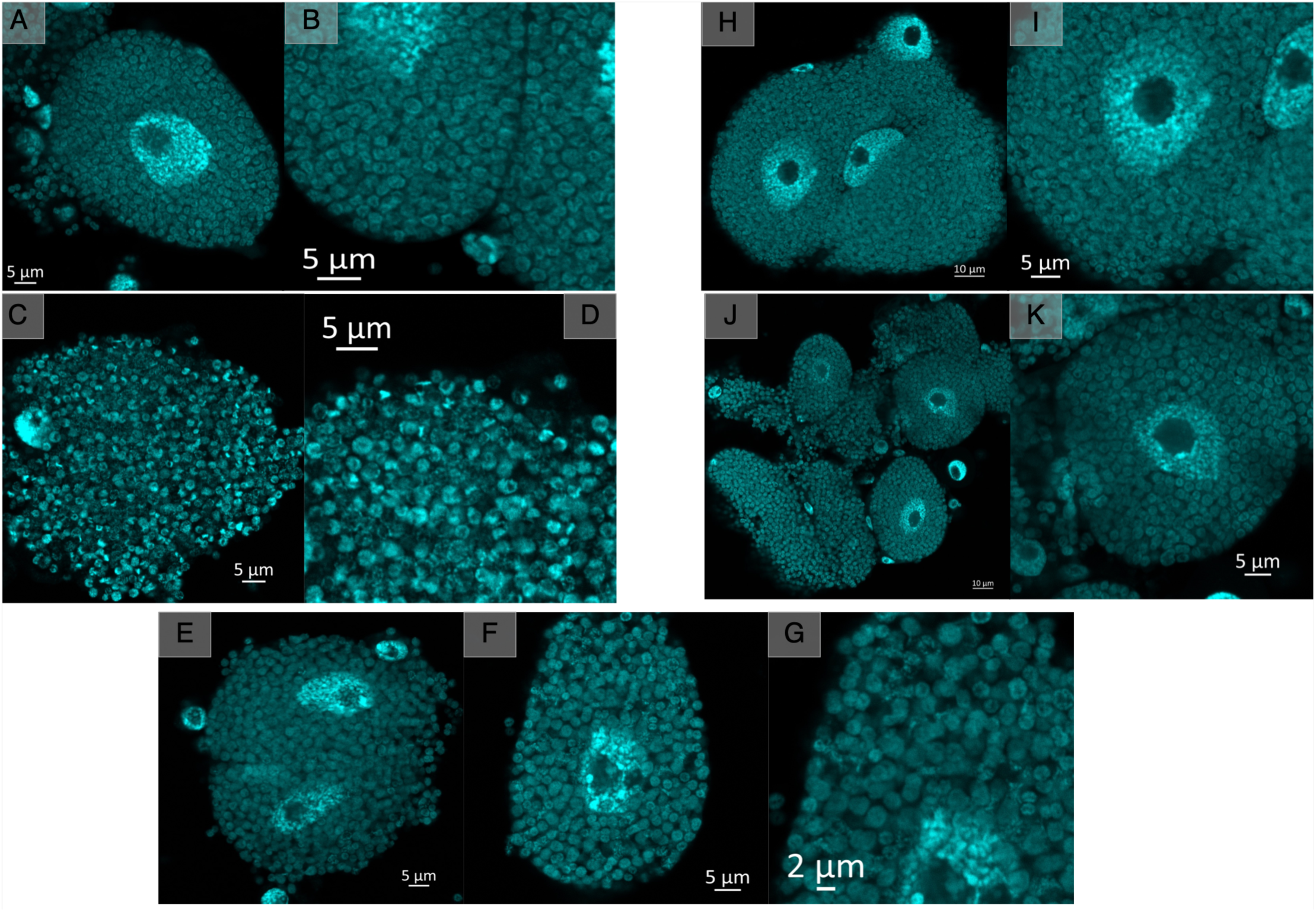
Super-resolution imaging of aphid bacteriocytes and *Buchnera* cells reveals effects of siRNA and cholesterol supplementation treatments at 46 h post-treatment. **A-B**: siRNA universal control treatment. **C-D:** ApW584 treatment. **E-G:** ApB744 treatment. H-I: Negative control (NegControl) for cholesterol supplementation treatment. **J-K:** Cholesterol supplementation treatment. DAPI staining was used to visualize DNA materials (cyan). (Scale bars: A-F, I & K = 5 µm; G = 2 µm; H & J = 10 µm)

**Fig. S10.**
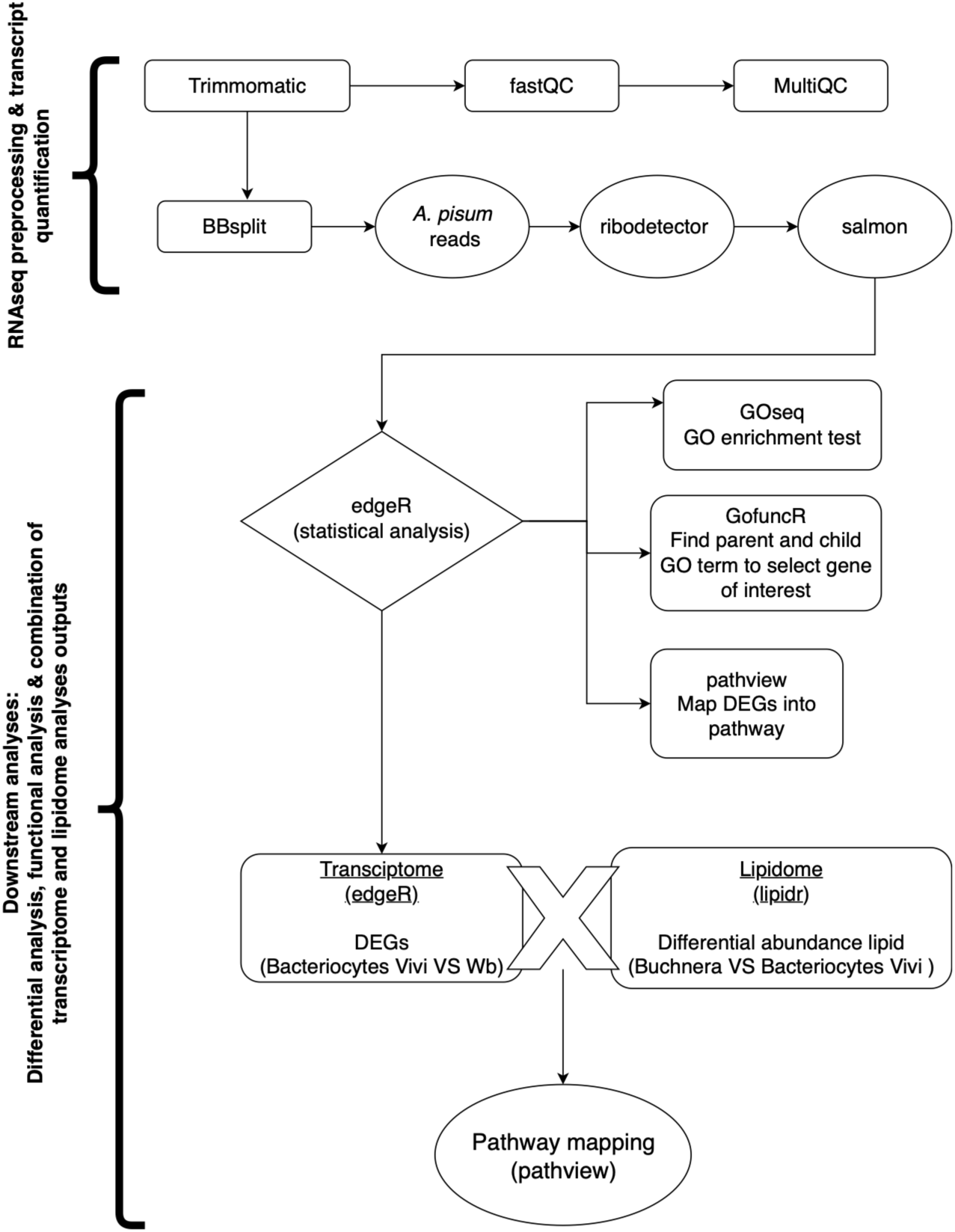
Workflow of *A. pisum* RNA-seq preprocessing, downstream analyses, and integration with lipidome data. Preprocessing began with adaptor trimming and quality control, followed by segregation of reads by reference genomes. *A. pisum* reads were subjected to rRNA removal, and the cleaned data were quantified using the mapping-based mode. Transcript count data were analyzed and differentially expressed genes (DEGs) were identified for subsequent downstream analyses. Integration of transcriptome and lipidome data was performed via KEGG pathway mapping, in which significant DEGs and enriched lipid species from the *Buchnera* lipidome were jointly mapped onto a single KEGG pathway to provide a comprehensive overview.

**Fig. S11.**
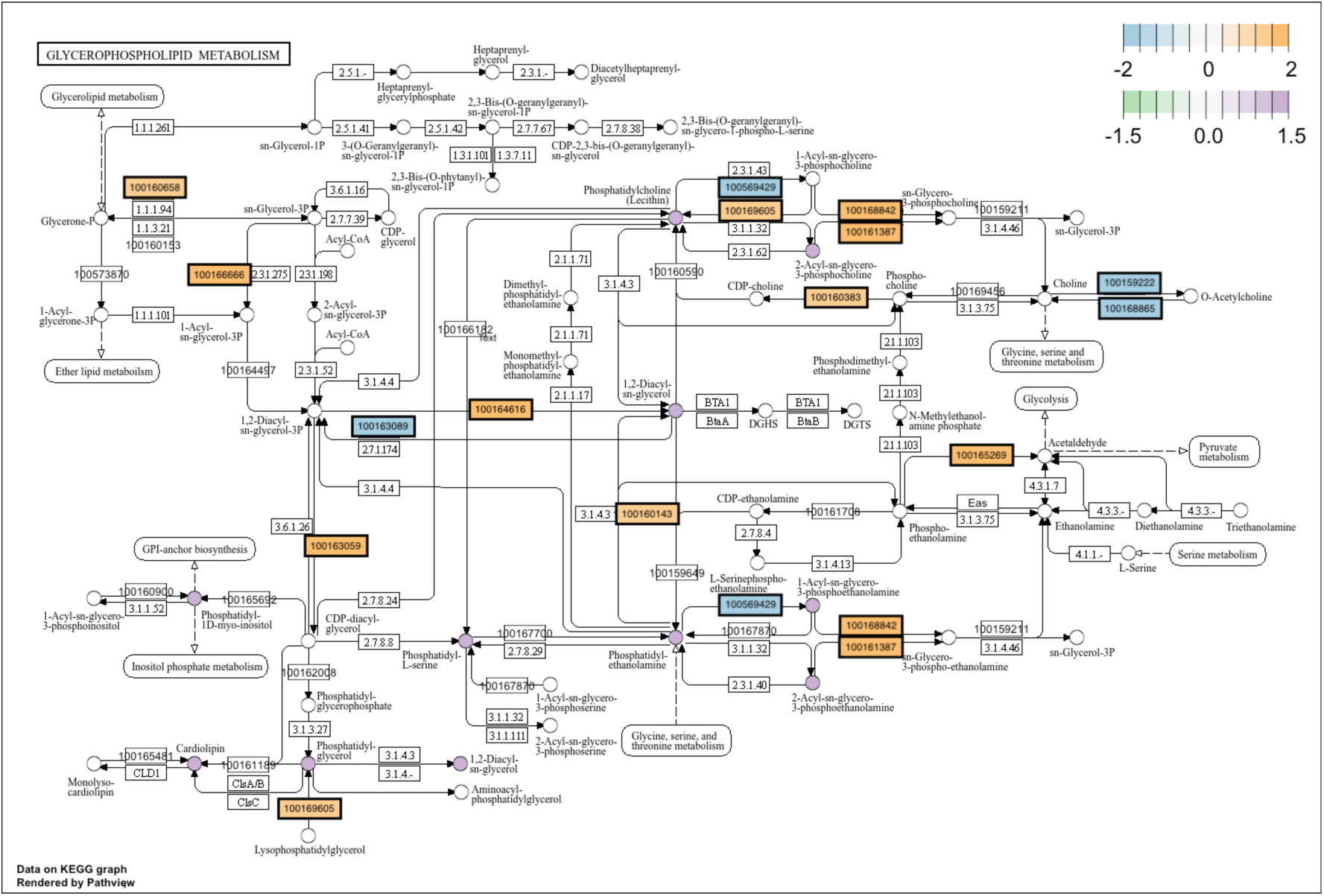
DEGs in viviparous bacteriocytes compared to whole body *A. pisum* and enriched lipid species in *Buchnera* are integrated and visualized in KEGG pathway, glycerophospholipid metabolism (api00564). DEGs identified in viviparous bacteriocytes versus whole body are mapped into KEGG glycerophospholipid metabolism pathway using R package, “Pathview” [11], [12]. When multiple transcripts or paralogs mapped to the same enzyme, the gene with the largest logFC was retained. Upregulated and downregulated genes in bacteriocytes are indicated in yellow and blue, respectively. Lipid species significantly enriched in isolated *Buchnera* (adjusted *p* <0.05, logFC >1) were grouped by lipid class and manually mapped to the corresponding KEGG compounds in the pathway. Lipid classes containing significantly enriched species are indicated in purple.

## Dataset S1

Multi-sheet Excel file containing lipidomic and transcriptomic data used for enrichment analysis and pathway integration.

The original MS Excel file, **DatasetS1.xlsx**, is available for download from our Figshare repository:

https://doi.org/10.6084/m9.figshare.34706211

For this PDF version, each worksheet in the Excel file has been converted to PDF and is included on the following pages.

**Sheet 1: Enriched_LipidSpecies_Buchnera–** Lipid species significantly enriched in *Buchnera* relative to bacteriocytes.
**Sheet 2: Enriched_LipidSpeciesBacterio–** Lipid species significantly enriched in bacteriocytes relative to *Buchnera*.
**Sheet 3: dePhosphoBchnera_Bacterioc** Results of differential abundance analysisof phosphorylated lipid species between *Buchnera* and aphid bacteriocytes.
**Sheet 4: Table_GO_lipid_DEGs –** This sheet lists selected lipid metabolism-related Gene Ontology (GO) terms identified by GOfuncR.
**Sheet 5: AcetylCoA_transporters –** This sheet presents the differential gene expression output generated by the “edgeR” package, comparing viviparous aphid bacteriocytes to whole body samples. It includes only the Acetyl-CoA Transporter 1 genes found in the transcriptome.
**Sheet 6: Lipid_class_summary_Buchnera –** This table summarizes lipid species that were significantly enriched in isolated *Buchnera* fractions (adjusted p < 0.05, logFC > 1). Lipid species were aggregated by class to compute mean and maximum logFC values and then manually matched to KEGG compound IDs corresponding to the *A. pisum* glycerophospholipid metabolism pathway (api00564).
**Sheet 7: api00564_gene_matches_max –** This table lists *A. pisum* genes that were significantly differentially expressed in viviparous bacteriocytes (adjusted p < 0.05) and successfully mapped to KEGG enzyme entries using their ENTREZ IDs.
**Sheet 8: LC_MSMS_sample_detail –** Tissue samples prepared for LC-MS/MS analyses. The table included the number of replicates, sample description and sample amount used in the experiment.
**Sheet 9: sequence_QC–** Summary of RNA-seq quality metrics for each sample.
**Sheet 10: siRNA_list–** Details of siRNAs assays
**Sheet 11: PrimerList–** List of primers used for quantitative PCR analyses

### 1 Enriched_LipidSpecies_Buchnera

This sheet lists lipid species that were significantly enriched in *Buchnera* compared to host bacteriocytes samples, based on differential abundance analysis using the “lipidr” R package. Each row corresponds to a lipid molecule with its corresponding class, fatty acid chain composition, and statistical output (adjusted *p* value <0.05, logFC > 1).

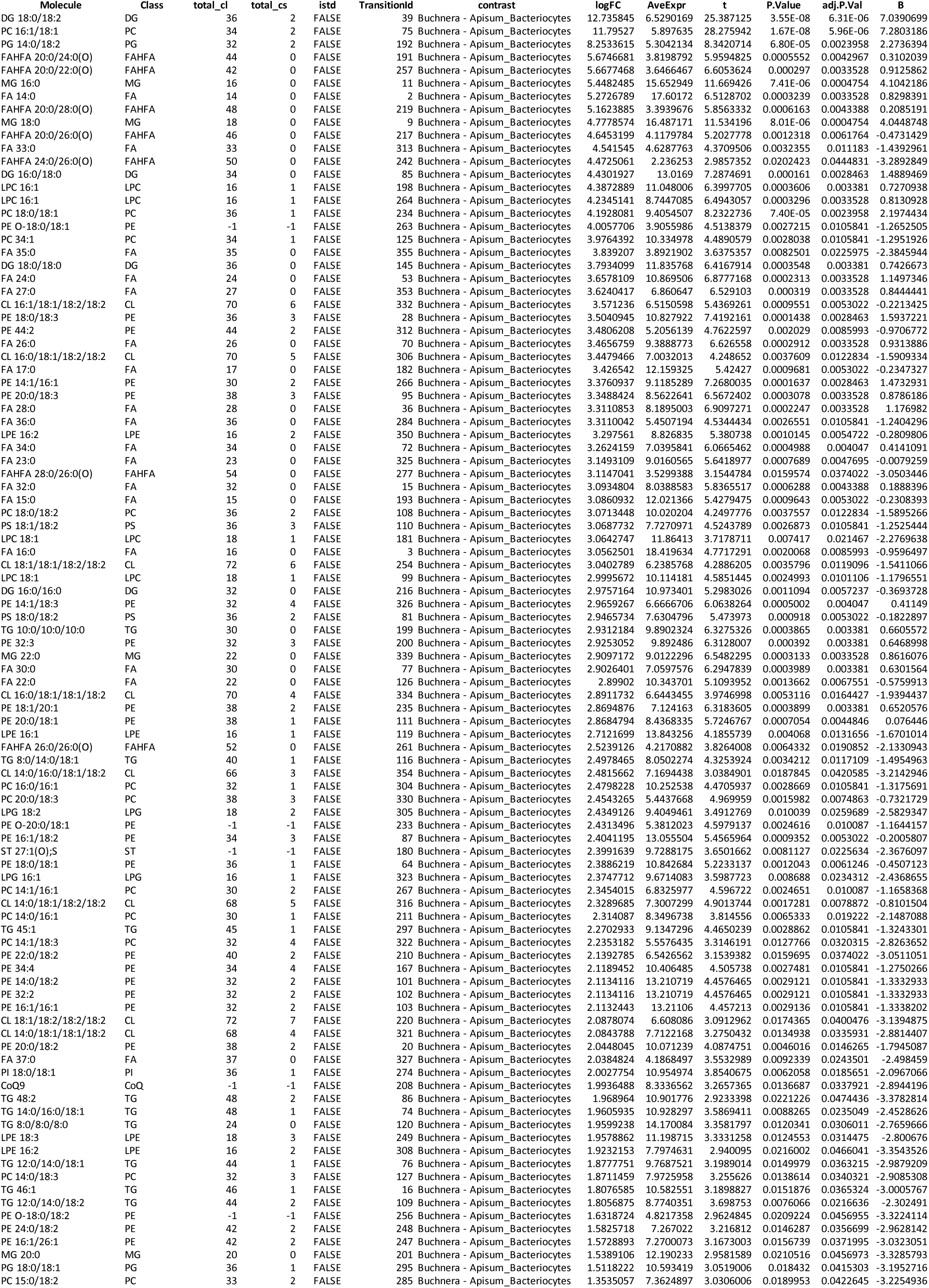

### 2 Enriched_LipidSpeciesBacterio

This sheet lists lipid species that were significantly enriched in host bacteriocytes compared to *Buchnera* samples, based on differential abundance analysis using the “lipidr” R package. Each row corresponds to a lipid molecule with its corresponding class, fatty acid chain composition, and statistical output (adjusted *p* value <0.05, logFC < -1).

| Molecule | Class | total_cl | total_cs | istd | TransitionId | contrast | logFC | AveExpr | t | P.Value | adj.P.Val | B |
|---|---|---|---|---|---|---|---|---|---|---|---|---|
| DG 18:1/18:1 | DG | 36 | 2 | FALSE | 17 | Buchnera - Apisum_Bacteriocytes | -13.802984 | 8.2850731 | -16.950406 | 5.82E-07 | 6.91E-05 | 5.7803622 |
| PC 16:0/18:2 | PC | 34 | 2 | FALSE | 54 | Buchnera - Apisum_Bacteriocytes | -9.470229 | 5.4663727 | -9.9019235 | 2.22E-05 | 0.0011269 | 3.2367218 |
| PG 32:2 | PG | 32 | 2 | FALSE | 278 | Buchnera - Apisum_Bacteriocytes | -9.4630194 | 5.4627679 | -14.67391 | 1.57E-06 | 0.0001394 | 5.1881188 |
| LPE 19:1 | LPE | 19 | 1 | FALSE | 194 | Buchnera - Apisum_Bacteriocytes | -9.3809127 | 5.4244392 | -8.946347 | 4.32E-05 | 0.0019208 | 2.6721505 |
| DG 18:1/18:2 | DG | 36 | 3 | FALSE | 69 | Buchnera - Apisum_Bacteriocytes | -8.9130504 | 9.322972 | -3.6798137 | 0.0077997 | 0.0220372 | -2.3278332 |
| HexCer 19:2(O2)/16:1(O) | HexCer | 19 | 2 | FALSE | 25 | Buchnera - Apisum_Bacteriocytes | -8.1596109 | 10.74462 | -5.8337964 | 0.0006306 | 0.0043388 | 0.1860938 |
| HexCer 18:2(O2)/16:1(O) | HexCer | 18 | 2 | FALSE | 150 | Buchnera - Apisum_Bacteriocytes | -7.913832 | 7.5727836 | -5.7293833 | 0.000702 | 0.0044846 | 0.0812121 |
| DLCL 18:1/18:1 | DLCL | 36 | 2 | FALSE | 338 | Buchnera - Apisum_Bacteriocytes | -6.3995186 | 5.3428314 | -6.3160406 | 0.0003908 | 0.003381 | 0.649906 |
| DG 8:0/14:0 | DG | 22 | 0 | FALSE | 91 | Buchnera - Apisum_Bacteriocytes | -6.0913928 | 11.261797 | -8.5743299 | 5.69E-05 | 0.0022507 | 2.4309028 |
| LPS 16:0 | LPS | 16 | 0 | FALSE | 231 | Buchnera - Apisum_Bacteriocytes | -6.0100687 | 6.9525208 | -4.8436861 | 0.0018464 | 0.0083206 | -0.8763334 |
| LPE 17:1 | LPE | 17 | 1 | FALSE | 107 | Buchnera - Apisum_Bacteriocytes | -5.7480247 | 10.254603 | -6.5283438 | 0.0003192 | 0.0033528 | 0.8437619 |
| Cer 17:1(O2)/16:0 | Cer | -1 | -1 | FALSE | 349 | Buchnera - Apisum_Bacteriocytes | -5.1090557 | 6.5950554 | -4.7124775 | 0.0021505 | 0.0090067 | -1.0288931 |
| Cer 17:0(O2)/16:0 | Cer | -1 | -1 | FALSE | 343 | Buchnera - Apisum_Bacteriocytes | -4.7934767 | 6.2194738 | -3.6111903 | 0.008545 | 0.0232216 | -2.4200967 |
| Cer 17:1(O2)/22:0(2OH) | Cer | 17 | 1 | FALSE | 156 | Buchnera - Apisum_Bacteriocytes | -4.782132 | 8.5538453 | -7.4866502 | 0.0001358 | 0.0028463 | 1.6465697 |
| Cer 17:1(O2)/18:0(2OH) | Cer | 17 | 1 | FALSE | 139 | Buchnera - Apisum_Bacteriocytes | -4.6388681 | 10.435276 | -8.1178334 | 8.10E-05 | 0.0024032 | 2.116728 |
| Cer 17:1(O2)/16:0(2OH) | Cer | 17 | 1 | FALSE | 346 | Buchnera - Apisum_Bacteriocytes | -4.6378623 | 6.7763746 | -3.0533868 | 0.0183934 | 0.0415303 | -3.1931645 |
| Cer 17:1(O2)/20:0(2OH) | Cer | 17 | 1 | FALSE | 149 | Buchnera - Apisum_Bacteriocytes | -4.4587812 | 10.360365 | -7.6931491 | 0.0001142 | 0.0028463 | 1.8051845 |
| LPG 17:1 | LPG | 17 | 1 | FALSE | 246 | Buchnera - Apisum_Bacteriocytes | -4.2893745 | 8.8072546 | -5.875271 | 0.0006045 | 0.0043388 | 0.2273052 |
| LPS 18:1 | LPS | 18 | 1 | FALSE | 227 | Buchnera - Apisum_Bacteriocytes | -4.2037169 | 6.2220614 | -7.6460332 | 0.0001188 | 0.0028463 | 1.7694158 |
| HexCer 17:0(O2)/22:0(O) | HexCer | 17 | 0 | FALSE | 175 | Buchnera - Apisum_Bacteriocytes | -4.1564209 | 8.5085297 | -5.9046845 | 0.0005868 | 0.0043388 | 0.2563783 |
| LPS 18:2 | LPS | 18 | 2 | FALSE | 140 | Buchnera - Apisum_Bacteriocytes | -4.1040822 | 9.2448139 | -5.8289091 | 0.0006338 | 0.0043388 | 0.1812209 |
| LPE 16:0 | LPE | 16 | 0 | FALSE | 115 | Buchnera - Apisum_Bacteriocytes | -4.081711 | 10.707749 | -5.5579924 | 0.0008397 | 0.0050665 | -0.0945079 |
| Cer 17:1(O2)/18:0 | Cer | -1 | -1 | FALSE | 106 | Buchnera - Apisum_Bacteriocytes | -4.0063832 | 11.925202 | -6.5097402 | 0.0003249 | 0.0033528 | 0.8270189 |
| PI 15:0/18:2 | PI | 33 | 2 | FALSE | 302 | Buchnera - Apisum_Bacteriocytes | -3.8455614 | 7.1629601 | -5.3419292 | 0.001058 | 0.0056203 | -0.3224607 |
| DLCL 18:1/18:2 | DLCL | 36 | 3 | FALSE | 328 | Buchnera - Apisum_Bacteriocytes | -3.8171101 | 6.9763002 | -4.7714347 | 0.0020075 | 0.0085993 | -0.9599923 |
| PS 18:3/18:3 | PS | 36 | 6 | FALSE | 299 | Buchnera - Apisum_Bacteriocytes | -3.691644 | 2.5798048 | -4.4301367 | 0.0030107 | 0.0106759 | -1.3667951 |
| Cer 17:0(O2)/22:0(2OH) | Cer | 17 | 0 | FALSE | 196 | Buchnera - Apisum_Bacteriocytes | -3.6172777 | 8.6881878 | -7.2172122 | 0.0001711 | 0.0028463 | 1.4322356 |
| PE 16:0/19:1 | PE | 35 | 1 | FALSE | 154 | Buchnera - Apisum_Bacteriocytes | -3.5414943 | 11.40601 | -7.3209683 | 0.0001564 | 0.0028463 | 1.5157812 |
| PE 14:0/16:0 | PE | 30 | 0 | FALSE | 164 | Buchnera - Apisum_Bacteriocytes | -3.4565958 | 11.232777 | -7.1855736 | 0.0001759 | 0.0028463 | 1.4065041 |
| PE 18:0/17:1 | PE | 35 | 1 | FALSE | 237 | Buchnera - Apisum_Bacteriocytes | -3.3831158 | 9.6877785 | -4.4367538 | 0.0029866 | 0.0106759 | -1.3587253 |
| HexCer 17:1(O2)/22:0(O) | HexCer | 17 | 1 | FALSE | 61 | Buchnera - Apisum_Bacteriocytes | -3.3420965 | 10.383811 | -7.3751658 | 0.0001493 | 0.0028463 | 1.5589161 |
| Cer 17:0(O2)/18:0 | Cer | -1 | -1 | FALSE | 152 | Buchnera - Apisum_Bacteriocytes | -3.3407568 | 9.19814 | -5.7301708 | 0.0007014 | 0.0044846 | 0.0820092 |
| Cer 17:1(O2)/22:0 | Cer | -1 | -1 | FALSE | 41 | Buchnera - Apisum_Bacteriocytes | -3.302972 | 10.015783 | -6.8437711 | 0.0002385 | 0.0033528 | 1.120698 |
| DG 16:1/16:1 | DG | 32 | 2 | FALSE | 26 | Buchnera - Apisum_Bacteriocytes | -3.2654519 | 12.878524 | -4.0155725 | 0.0050413 | 0.015743 | -1.8866805 |
| DG 18:2/18:2 | DG | 36 | 4 | FALSE | 37 | Buchnera - Apisum_Bacteriocytes | -3.2525749 | 12.8899 | -4.8312072 | 0.0018732 | 0.0083357 | -0.8907214 |
| HexCer 17:1(O2)/20:0(O) | HexCer | 17 | 1 | FALSE | 52 | Buchnera - Apisum_Bacteriocytes | -3.2411719 | 11.868207 | -6.7006656 | 0.0002719 | 0.0033528 | 0.9966647 |
| Cer 17:1(O2)/20:0 | Cer | -1 | -1 | FALSE | 142 | Buchnera - Apisum_Bacteriocytes | -3.1897904 | 11.198204 | -5.5051059 | 0.000888 | 0.0052691 | -0.1496357 |
| DG 16:0/16:1 | DG | 32 | 1 | FALSE | 62 | Buchnera - Apisum_Bacteriocytes | -3.1323349 | 13.110417 | -6.0327271 | 0.0005159 | 0.0040815 | 0.3814666 |
| SPB 17:1(O2) | SPB | 17 | 1 | FALSE | 134 | Buchnera - Apisum_Bacteriocytes | -3.0676097 | 10.924632 | -3.4996849 | 0.0099256 | 0.0259689 | -2.5714517 |
| PG 16:0/19:1 | PG | 35 | 1 | FALSE | 223 | Buchnera - Apisum_Bacteriocytes | -2.9319028 | 10.19116 | -4.9816246 | 0.0015772 | 0.0074863 | -0.7189857 |
| DG 16:1/18:3 | DG | 34 | 4 | FALSE | 121 | Buchnera - Apisum_Bacteriocytes | -2.8716913 | 10.649098 | -3.4892507 | 0.0100666 | 0.0259689 | -2.5857034 |
| SM 39:1(O3) | SM | 39 | 1 | FALSE | 179 | Buchnera - Apisum_Bacteriocytes | -2.807252 | 7.1707864 | -4.9761984 | 0.0015869 | 0.0074863 | -0.7251169 |
| PC 18:1/18:1 | PC | 36 | 2 | FALSE | 190 | Buchnera - Apisum_Bacteriocytes | -2.7840437 | 11.026951 | -5.6317429 | 0.0007771 | 0.0047695 | -0.0183476 |
| LPC 18:2 | LPC | 18 | 2 | FALSE | 22 | Buchnera - Apisum_Bacteriocytes | -2.7762279 | 13.990498 | -4.2944924 | 0.0035537 | 0.0119096 | -1.5338085 |
| MG 18:1 | MG | 18 | 1 | FALSE | 55 | Buchnera - Apisum_Bacteriocytes | -2.7661167 | 12.999891 | -3.9436901 | 0.0055271 | 0.0169626 | -1.9796511 |
| DG 16:1/18:2 | DG | 34 | 3 | FALSE | 29 | Buchnera - Apisum_Bacteriocytes | -2.7533186 | 12.71178 | -3.0798551 | 0.0177196 | 0.0404369 | -3.1556727 |
| DG 18:2/18:3 | DG | 36 | 5 | FALSE | 122 | Buchnera - Apisum_Bacteriocytes | -2.7382891 | 11.297637 | -3.8676923 | 0.006097 | 0.0183943 | -2.0788305 |
| MG 16:1 | MG | 16 | 1 | FALSE | 130 | Buchnera - Apisum_Bacteriocytes | -2.6747348 | 11.496021 | -4.2931217 | 0.0035598 | 0.0119096 | -1.5355115 |
| Cer 17:0(O2)/20:0 | Cer | -1 | -1 | FALSE | 281 | Buchnera - Apisum_Bacteriocytes | -2.6554417 | 8.1567331 | -3.8003966 | 0.0066553 | 0.0194203 | -2.1674029 |
| PE 16:0/18:2 | PE | 34 | 2 | FALSE | 32 | Buchnera - Apisum_Bacteriocytes | -2.6540435 | 14.663276 | -5.7870247 | 0.0006615 | 0.0044435 | 0.1393132 |
| LPE 14:0 | LPE | 14 | 0 | FALSE | 255 | Buchnera - Apisum_Bacteriocytes | -2.553094 | 7.5873325 | -4.4251807 | 0.0030288 | 0.0106759 | -1.3728439 |
| LPI 18:1 | LPI | 18 | 1 | FALSE | 162 | Buchnera - Apisum_Bacteriocytes | -2.527803 | 10.622419 | -4.0783826 | 0.0046548 | 0.0146647 | -1.8061186 |
| HexCer 17:1(O2)/24:0(O) | HexCer | 17 | 1 | FALSE | 117 | Buchnera - Apisum_Bacteriocytes | -2.4765473 | 9.6096188 | -5.3284755 | 0.0010735 | 0.0056203 | -0.3368956 |
| HexCer 17:1(O2)/20:0 | HexCer | -1 | -1 | FALSE | 166 | Buchnera - Apisum_Bacteriocytes | -2.3966834 | 10.919519 | -4.9468644 | 0.0016407 | 0.0075855 | -0.7583449 |
| DG 18:3/18:3 | DG | 36 | 6 | FALSE | 243 | Buchnera - Apisum_Bacteriocytes | -2.3361233 | 9.1052645 | -3.6373736 | 0.0082519 | 0.0225975 | -2.3848126 |
| PE 33:2 | PE | 33 | 2 | FALSE | 241 | Buchnera - Apisum_Bacteriocytes | -2.3196184 | 10.403216 | -3.6541244 | 0.0080701 | 0.0225634 | -2.3622917 |
| LPI 18:2 | LPI | 18 | 2 | FALSE | 136 | Buchnera - Apisum_Bacteriocytes | -2.2252524 | 11.186112 | -3.1760517 | 0.0154835 | 0.0369942 | -3.0200028 |
| PG 14:0/16:0 | PG | 30 | 0 | FALSE | 288 | Buchnera - Apisum_Bacteriocytes | -2.2229533 | 9.6544538 | -4.8114718 | 0.0019164 | 0.0084227 | -0.9135281 |
| PE 16:0/17:1 | PE | 33 | 1 | FALSE | 19 | Buchnera - Apisum_Bacteriocytes | -2.2053836 | 15.724148 | -3.1388536 | 0.0163103 | 0.0379175 | -3.0723512 |
| HexCer 17:1(O2)/18:0(O) | HexCer | 17 | 1 | FALSE | 124 | Buchnera - Apisum_Bacteriocytes | -2.1813256 | 12.604814 | -5.0594305 | 0.0014447 | 0.0070455 | -0.6315954 |
| DG 16:0/18:1 | DG | 34 | 1 | FALSE | 59 | Buchnera - Apisum_Bacteriocytes | -2.1777484 | 13.064192 | -3.1348289 | 0.0164025 | 0.0379175 | -3.0780238 |
| PC 18:2/18:3 | PC | 36 | 5 | FALSE | 31 | Buchnera - Apisum_Bacteriocytes | -2.1509488 | 10.110322 | -2.9150887 | 0.0223847 | 0.0477184 | -3.3900731 |
| PE 18:1/19:1 | PE | 37 | 2 | FALSE | 265 | Buchnera - Apisum_Bacteriocytes | -2.1224984 | 8.5455059 | -4.3730522 | 0.0032272 | 0.011183 | -1.4367119 |
| Cer 16:1(O2)/18:0 | Cer | -1 | -1 | FALSE | 301 | Buchnera - Apisum_Bacteriocytes | -2.0851715 | 8.9204186 | -4.1131689 | 0.0044548 | 0.0142875 | -1.7617737 |
| PE 18:2/18:2 | PE | 36 | 4 | FALSE | 10 | Buchnera - Apisum_Bacteriocytes | -2.049535 | 13.59822 | -3.3778246 | 0.0117146 | 0.0300029 | -2.738806 |
| PE 14:0/17:1 | PE | 31 | 1 | FALSE | 184 | Buchnera - Apisum_Bacteriocytes | -1.8884797 | 11.2346 | -3.5851833 | 0.0088473 | 0.0235049 | -2.455241 |
| Cer 17:1(O2)/24:0(2OH) | Cer | 17 | 1 | FALSE | 165 | Buchnera - Apisum_Bacteriocytes | -1.7144695 | 9.4845678 | -3.8843533 | 0.0059668 | 0.0181554 | -2.0570102 |
| FA 18:3 | FA | 18 | 3 | FALSE | 45 | Buchnera - Apisum_Bacteriocytes | -1.6133725 | 14.330884 | -2.9876378 | 0.0201877 | 0.0444831 | -3.2865761 |
| SM 41:1(O2) | SM | 41 | 1 | FALSE | 296 | Buchnera - Apisum_Bacteriocytes | -1.5664743 | 6.8350722 | -3.7026689 | 0.0075673 | 0.0216636 | -2.297258 |

### 3 dePhosphoBchnera_Bacterioc

Results of differential abundance analysis of phosphorylated lipid species between *Buchnera* and aphid bacteriocytes. Each row represents an individual phospholipid molecule with corresponding class annotation, quantification metrics, and statistical outputs (log fold change, average expression, t-value, p-value, and adjusted p-value)

| Molecule | Class | total_cl | total_cs | istd | TransitionId | contrast | logFC | AveExpr | t | P.Value | adj.P.Val | B |
|---|---|---|---|---|---|---|---|---|---|---|---|---|
| PE 14:0/18:3 | PE | 32 | 3 | FALSE | 1 | Buchnera - Apisum_Bacteriocytes | -1.5275556 | 10.950118 | -2.566472 | 0.0322915 | 0.0671812 | -3.8451929 |
| PE 36:2 | PE | 36 | 2 | FALSE | 5 | Buchnera - Apisum_Bacteriocytes | 1.1232207 | 14.052161 | 2.4661267 | 0.0378589 | 0.0747667 | -3.9993398 |
| PE 18:0/18:2 | PE | 36 | 2 | FALSE | 6 | Buchnera - Apisum_Bacteriocytes | 1.1230612 | 14.052116 | 2.4658874 | 0.0378732 | 0.0747667 | -3.9997066 |
| PC 18:2/18:2 | PC | 36 | 4 | FALSE | 7 | Buchnera - Apisum_Bacteriocytes | -2.5269457 | 12.466098 | -2.6156499 | 0.0298731 | 0.0628725 | -3.7694539 |
| PE 18:2/18:2 | PE | 36 | 4 | FALSE | 10 | Buchnera - Apisum_Bacteriocytes | -2.049535 | 13.59822 | -3.3990216 | 0.0088503 | 0.0246448 | -2.5681852 |
| PE 18:1/18:2 | PE | 36 | 3 | FALSE | 12 | Buchnera - Apisum_Bacteriocytes | -0.3401873 | 12.880227 | -0.6817117 | 0.5139541 | 0.5875042 | -6.2306268 |
| PC 18:1/18:2 | PC | 36 | 3 | FALSE | 14 | Buchnera - Apisum_Bacteriocytes | -0.0034421 | 10.972507 | -0.0034945 | 0.9972942 | 0.9972942 | -6.4778887 |
| PI 18:1/18:2 | PI | 36 | 3 | FALSE | 18 | Buchnera - Apisum_Bacteriocytes | -0.4509456 | 12.735636 | -0.8638808 | 0.411874 | 0.4904552 | -6.0872561 |
| PE 16:0/17:1 | PE | 33 | 1 | FALSE | 19 | Buchnera - Apisum_Bacteriocytes | -2.2053836 | 15.724148 | -3.2179616 | 0.0116627 | 0.0293187 | -2.8428093 |
| PE 20:0/18:2 | PE | 38 | 2 | FALSE | 20 | Buchnera - Apisum_Bacteriocytes | 2.0448045 | 10.071239 | 3.9804136 | 0.0037536 | 0.0119192 | -1.7110665 |
| LPC 18:2 | LPC | 18 | 2 | FALSE | 22 | Buchnera - Apisum_Bacteriocytes | -2.7762279 | 13.990498 | -4.3589423 | 0.0022029 | 0.0083069 | -1.1780757 |
| LPE 18:2 | LPE | 18 | 2 | FALSE | 23 | Buchnera - Apisum_Bacteriocytes | -0.7754677 | 12.883754 | -1.1017234 | 0.3014417 | 0.3842321 | -5.8589995 |
| PC 14:0/18:2 | PC | 32 | 2 | FALSE | 24 | Buchnera - Apisum_Bacteriocytes | 0.7312582 | 0.3656291 | 1.3523805 | 0.2118744 | 0.2883403 | -5.5755789 |
| PI 18:2/18:2 | PI | 36 | 4 | FALSE | 27 | Buchnera - Apisum_Bacteriocytes | -1.2193484 | 12.082216 | -2.0223821 | 0.0764213 | 0.1356588 | -4.6674231 |
| PE 18:0/18:3 | PE | 36 | 3 | FALSE | 28 | Buchnera - Apisum_Bacteriocytes | 3.5040945 | 10.827922 | 7.139128 | 8.09E-05 | 0.0016313 | 2.0533573 |
| PE 16:0/16:1 | PE | 32 | 1 | FALSE | 30 | Buchnera - Apisum_Bacteriocytes | -0.4135732 | 15.106904 | -0.6323377 | 0.5441549 | 0.6155753 | -6.2643313 |
| PC 18:2/18:3 | PC | 36 | 5 | FALSE | 31 | Buchnera - Apisum_Bacteriocytes | -2.1509488 | 10.110322 | -3.0042077 | 0.0162279 | 0.038146 | -3.1702417 |
| PE 16:0/18:2 | PE | 34 | 2 | FALSE | 32 | Buchnera - Apisum_Bacteriocytes | -2.6540435 | 14.663276 | -5.5322994 | 0.000482 | 0.0036146 | 0.3305281 |
| PE 16:1/18:1 | PE | 34 | 2 | FALSE | 34 | Buchnera - Apisum_Bacteriocytes | -0.8540858 | 14.303287 | -1.0701293 | 0.3146325 | 0.3982411 | -5.891773 |
| PE 18:1/18:1 | PE | 36 | 2 | FALSE | 35 | Buchnera - Apisum_Bacteriocytes | 0.3609006 | 12.261115 | 0.7776471 | 0.4583255 | 0.5386812 | -6.1587566 |
| PE 16:0/18:1 | PE | 34 | 1 | FALSE | 40 | Buchnera - Apisum_Bacteriocytes | 0.2168057 | 15.260345 | 0.345315 | 0.7384294 | 0.7753588 | -6.4131259 |
| LPE 18:0 | LPE | 18 | 0 | FALSE | 42 | Buchnera - Apisum_Bacteriocytes | -0.7415901 | 10.479252 | -1.1103935 | 0.2978993 | 0.38267 | -5.8498823 |
| PE 18:2/18:3 | PE | 36 | 5 | FALSE | 43 | Buchnera - Apisum_Bacteriocytes | -0.9312476 | 11.428901 | -1.7108499 | 0.1240355 | 0.2009234 | -5.1110095 |
| PI 18:1/18:1 | PI | 36 | 2 | FALSE | 49 | Buchnera - Apisum_Bacteriocytes | 0.3592295 | 12.391506 | 0.7176202 | 0.4926533 | 0.5679634 | -6.2046981 |
| PG 16:0/17:1 | PG | 33 | 1 | FALSE | 51 | Buchnera - Apisum_Bacteriocytes | -2.1449352 | 13.930432 | -2.3567294 | 0.045035 | 0.0867163 | -4.16651 |
| PC 16:0/18:2 | PC | 34 | 2 | FALSE | 54 | Buchnera - Apisum_Bacteriocytes | -9.470229 | 5.4663727 | -10.422206 | 4.66E-06 | 0.0002809 | 4.6051646 |
| PC 16:1/18:2 | PC | 34 | 3 | FALSE | 57 | Buchnera - Apisum_Bacteriocytes | 1.8992592 | 10.720705 | 2.4060673 | 0.0416438 | 0.0810486 | -4.0912539 |
| PG 18:2/18:2 | PG | 36 | 4 | FALSE | 58 | Buchnera - Apisum_Bacteriocytes | -0.472794 | 10.097051 | -0.8197121 | 0.4352409 | 0.5148928 | -6.1246731 |
| PS 18:2/18:2 | PS | 36 | 4 | FALSE | 63 | Buchnera - Apisum_Bacteriocytes | 1.6668382 | 8.9692284 | 1.573761 | 0.152754 | 0.2363117 | -5.2956077 |
| PE 18:0/18:1 | PE | 36 | 1 | FALSE | 64 | Buchnera - Apisum_Bacteriocytes | 2.3886219 | 10.842684 | 4.9901238 | 0.0009492 | 0.0053687 | -0.3391104 |
| LPE 18:2 | LPE | 18 | 2 | FALSE | 65 | Buchnera - Apisum_Bacteriocytes | 1.2000064 | 14.472355 | 2.0154343 | 0.0772605 | 0.1357685 | -4.6776087 |
| PI 34:2 | PI | 34 | 2 | FALSE | 73 | Buchnera - Apisum_Bacteriocytes | -0.3832003 | 13.524763 | -0.5820324 | 0.5759788 | 0.6408288 | -6.2963025 |
| PC 16:1/18:1 | PC | 34 | 2 | FALSE | 75 | Buchnera - Apisum_Bacteriocytes | 11.79527 | 5.897635 | 26.411982 | 2.57E-09 | 4.66E-07 | 9.1839379 |
| PE 17:1/18:1 | PE | 35 | 2 | FALSE | 79 | Buchnera - Apisum_Bacteriocytes | -1.6376159 | 12.873475 | -1.715365 | 0.1231802 | 0.2009234 | -5.1048036 |
| PS 18:0/18:2 | PS | 36 | 2 | FALSE | 81 | Buchnera - Apisum_Bacteriocytes | 2.9465734 | 7.6304796 | 5.4041108 | 0.0005637 | 0.0036439 | 0.1764315 |
| PE 32:1 | PE | 32 | 1 | FALSE | 82 | Buchnera - Apisum_Bacteriocytes | 0.7312582 | 0.3656291 | 1.3523805 | 0.2118744 | 0.2883403 | -5.5755789 |
| LPC 18:2 | LPC | 18 | 2 | FALSE | 84 | Buchnera - Apisum_Bacteriocytes | 0.9505499 | 13.150114 | 1.1098951 | 0.298102 | 0.38267 | -5.8504078 |
| PE 16:1/18:2 | PE | 34 | 3 | FALSE | 87 | Buchnera - Apisum_Bacteriocytes | 2.4041195 | 13.055504 | 5.1675868 | 0.0007569 | 0.0045814 | -0.11475 |
| LPE 18:1 | LPE | 18 | 1 | FALSE | 88 | Buchnera - Apisum_Bacteriocytes | -0.7211572 | 11.296337 | -1.0296434 | 0.332189 | 0.4140043 | -5.9327164 |
| PG 16:0/18:1 | PG | 34 | 1 | FALSE | 89 | Buchnera - Apisum_Bacteriocytes | -0.8140246 | 13.578047 | -1.3377395 | 0.2164064 | 0.2923101 | -5.59319 |
| PC 14:0/18:1 | PC | 32 | 1 | FALSE | 90 | Buchnera - Apisum_Bacteriocytes | 0.7312582 | 0.3656291 | 1.3523805 | 0.2118744 | 0.2883403 | -5.5755789 |
| PE 20:0/18:3 | PE | 38 | 3 | FALSE | 95 | Buchnera - Apisum_Bacteriocytes | 3.3488424 | 8.5622641 | 6.4191778 | 0.0001732 | 0.00209 | 1.3274787 |
| PG 16:0/18:2 | PG | 34 | 2 | FALSE | 96 | Buchnera - Apisum_Bacteriocytes | -1.1731123 | 12.888201 | -1.5199823 | 0.1655779 | 0.2436554 | -5.3658305 |
| PG 34:2 | PG | 34 | 2 | FALSE | 97 | Buchnera - Apisum_Bacteriocytes | -1.1732497 | 12.888278 | -1.5202828 | 0.1655037 | 0.2436554 | -5.3654419 |
| PG 16:1/18:1 | PG | 34 | 2 | FALSE | 98 | Buchnera - Apisum_Bacteriocytes | -1.1732497 | 12.888278 | -1.5202828 | 0.1655037 | 0.2436554 | -5.3654419 |
| LPC 18:1 | LPC | 18 | 1 | FALSE | 99 | Buchnera - Apisum_Bacteriocytes | 2.9995672 | 10.114181 | 4.6610236 | 0.0014616 | 0.007107 | -0.7686039 |
| PE 14:0/18:2 | PE | 32 | 2 | FALSE | 101 | Buchnera - Apisum_Bacteriocytes | 2.1134116 | 13.210719 | 4.2929156 | 0.0024139 | 0.0084069 | -1.2694827 |
| PE 32:2 | PE | 32 | 2 | FALSE | 102 | Buchnera - Apisum_Bacteriocytes | 2.1134116 | 13.210719 | 4.2929156 | 0.0024139 | 0.0084069 | -1.2694827 |
| PE 16:1/16:1 | PE | 32 | 2 | FALSE | 103 | Buchnera - Apisum_Bacteriocytes | 2.1132443 | 13.21106 | 4.2925149 | 0.0024152 | 0.0084069 | -1.2700394 |
| PG 18:1/18:2 | PG | 36 | 3 | FALSE | 104 | Buchnera - Apisum_Bacteriocytes | -0.5877338 | 9.85992 | -1.0237402 | 0.3348107 | 0.4140043 | -5.9385855 |
| PC 34:4 | PC | 34 | 4 | FALSE | 105 | Buchnera - Apisum_Bacteriocytes | 1.2892354 | 8.9041414 | 1.8312359 | 0.1030208 | 0.1759128 | -4.9430914 |
| LPE 17:1 | LPE | 17 | 1 | FALSE | 107 | Buchnera - Apisum_Bacteriocytes | -5.7480247 | 10.254603 | -6.8325278 | 0.0001111 | 0.0016757 | 1.7528816 |
| PC 18:0/18:2 | PC | 36 | 2 | FALSE | 108 | Buchnera - Apisum_Bacteriocytes | 3.0713448 | 10.020204 | 4.3702555 | 0.0021688 | 0.0083069 | -1.1624816 |
| PS 18:1/18:2 | PS | 36 | 3 | FALSE | 110 | Buchnera - Apisum_Bacteriocytes | 3.0687732 | 7.7270971 | 4.6196514 | 0.0015449 | 0.007107 | -0.8238279 |
| PE 20:0/18:1 | PE | 38 | 1 | FALSE | 111 | Buchnera - Apisum_Bacteriocytes | 2.8684794 | 8.4368335 | 5.5765224 | 0.0004569 | 0.0036146 | 0.3830883 |
| PG 16:0/16:1 | PG | 32 | 1 | FALSE | 112 | Buchnera - Apisum_Bacteriocytes | -0.3277015 | 13.192545 | -0.4499349 | 0.6642541 | 0.71142 | -6.3684869 |
| LPE 16:0 | LPE | 16 | 0 | FALSE | 115 | Buchnera - Apisum_Bacteriocytes | -4.081711 | 10.707749 | -5.7251333 | 0.0003825 | 0.003297 | 0.5574775 |
| LPE 16:1 | LPE | 16 | 1 | FALSE | 119 | Buchnera - Apisum_Bacteriocytes | 2.7121699 | 13.843256 | 4.2496719 | 0.0025638 | 0.0087555 | -1.3297132 |
| PC 34:1 | PC | 34 | 1 | FALSE | 125 | Buchnera - Apisum_Bacteriocytes | 3.9764392 | 10.334978 | 4.7003096 | 0.0013871 | 0.0069739 | -0.7164182 |
| PC 14:0/18:3 | PC | 32 | 3 | FALSE | 127 | Buchnera - Apisum_Bacteriocytes | 1.8711459 | 7.9725958 | 3.2493892 | 0.0111143 | 0.0287383 | -2.7949363 |
| LPE 20:0 | LPE | 20 | 0 | FALSE | 129 | Buchnera - Apisum_Bacteriocytes | 0.3780709 | 9.0542029 | 0.6781547 | 0.5160948 | 0.5875042 | -6.2331308 |
| LPE 18:1 | LPE | 18 | 1 | FALSE | 133 | Buchnera - Apisum_Bacteriocytes | 0.9217769 | 12.214744 | 1.4714317 | 0.177972 | 0.2597817 | -5.4280612 |
| LPI 18:2 | LPI | 18 | 2 | FALSE | 136 | Buchnera - Apisum_Bacteriocytes | -2.2252524 | 11.186112 | -3.2550748 | 0.011018 | 0.0287383 | -2.7862841 |
| PS 18:2/18:3 | PS | 36 | 5 | FALSE | 138 | Buchnera - Apisum_Bacteriocytes | -0.4114571 | 7.7293782 | -0.6049888 | 0.561327 | 0.6310571 | -6.2820129 |
| LPS 18:2 | LPS | 18 | 2 | FALSE | 140 | Buchnera - Apisum_Bacteriocytes | -4.1040822 | 9.2448139 | -5.9772153 | 0.0002848 | 0.0027135 | 0.8454972 |
| PC 18:3/18:3 | PC | 36 | 6 | FALSE | 141 | Buchnera - Apisum_Bacteriocytes | 0.2476924 | 8.359163 | 0.3975929 | 0.7009407 | 0.7462956 | -6.3922324 |
| PS 16:0/18:2 | PS | 34 | 2 | FALSE | 146 | Buchnera - Apisum_Bacteriocytes | -1.1280903 | 9.7075502 | -1.0605674 | 0.3187125 | 0.4006039 | -5.9015509 |
| PG 17:1/18:1 | PG | 35 | 2 | FALSE | 147 | Buchnera - Apisum_Bacteriocytes | 2.2568401 | 9.6987361 | 0.7414316 | 0.4788429 | 0.5591649 | -6.186859 |
| PG 18:2/18:3 | PG | 36 | 5 | FALSE | 153 | Buchnera - Apisum_Bacteriocytes | -0.2855322 | 8.540621 | -0.4940236 | 0.6340726 | 0.6913683 | -6.3463247 |
| PE 16:0/19:1 | PE | 35 | 1 | FALSE | 154 | Buchnera - Apisum_Bacteriocytes | -3.5414943 | 11.40601 | -7.0806384 | 8.59E-05 | 0.0016313 | 1.9969968 |
| PE 18:3/18:3 | PE | 36 | 6 | FALSE | 157 | Buchnera - Apisum_Bacteriocytes | 0.425664 | 8.6202057 | 0.7226812 | 0.4896969 | 0.5679634 | -6.2009493 |
| LPI 16:0 | LPI | 16 | 0 | FALSE | 158 | Buchnera - Apisum_Bacteriocytes | -2.4942643 | 9.5281543 | -2.8043476 | 0.0221832 | 0.0489654 | -3.4782981 |
| PE 16:0/16:0 | PE | 32 | 0 | FALSE | 159 | Buchnera - Apisum_Bacteriocytes | -2.0819587 | 10.705307 | -2.4936803 | 0.0362401 | 0.0743574 | -3.9570768 |
| LPI 18:1 | LPI | 18 | 1 | FALSE | 162 | Buchnera - Apisum_Bacteriocytes | -2.527803 | 10.622419 | -4.116329 | 0.0030924 | 0.0101769 | -1.5172237 |
| PE 14:0/16:0 | PE | 30 | 0 | FALSE | 164 | Buchnera - Apisum_Bacteriocytes | -3.4565958 | 11.232777 | -6.9415215 | 9.91E-05 | 0.0016313 | 1.8611332 |
| PE 34:4 | PE | 34 | 4 | FALSE | 167 | Buchnera - Apisum_Bacteriocytes | 2.1189452 | 10.406485 | 4.3315855 | 0.0022878 | 0.0084069 | -1.2158664 |
| PI 16:0/18:1 | PI | 34 | 1 | FALSE | 168 | Buchnera - Apisum_Bacteriocytes | 1.2003364 | 12.417389 | 2.2315033 | 0.054928 | 0.1014487 | -4.3562374 |
| PE 19:0/18:2 | PE | 37 | 2 | FALSE | 170 | Buchnera - Apisum_Bacteriocytes | -0.8995588 | 8.0241023 | -1.6083681 | 0.1449824 | 0.2262226 | -5.2497381 |
| PI 18:2/18:3 | PI | 36 | 5 | FALSE | 171 | Buchnera - Apisum_Bacteriocytes | -1.3760792 | 9.6689688 | -2.4637317 | 0.038003 | 0.0747667 | -4.0030108 |
| PE 17:0/18:2 | PE | 35 | 2 | FALSE | 177 | Buchnera - Apisum_Bacteriocytes | 3.8555465 | 1.9277733 | 1.5484684 | 0.1586697 | 0.241338 | -5.3287983 |
| LPC 18:1 | LPC | 18 | 1 | FALSE | 181 | Buchnera - Apisum_Bacteriocytes | 3.0642747 | 11.86413 | 3.8709509 | 0.0043959 | 0.0137182 | -1.8691055 |
| PE 14:0/17:1 | PE | 31 | 1 | FALSE | 184 | Buchnera - Apisum_Bacteriocytes | -1.8884797 | 11.2346 | -3.5257081 | 0.0073136 | 0.0206837 | -2.377937 |
| LPE 16:1 | LPE | 16 | 1 | FALSE | 186 | Buchnera - Apisum_Bacteriocytes | 1.8206425 | 12.179821 | 2.7268502 | 0.0250614 | 0.0540014 | -3.597924 |
| LPC 14:0 | LPC | 14 | 0 | FALSE | 189 | Buchnera - Apisum_Bacteriocytes | -0.8619189 | 7.4103273 | -1.7856348 | 0.110555 | 0.1852819 | -5.0072777 |
| PC 18:1/18:1 | PC | 36 | 2 | FALSE | 190 | Buchnera - Apisum_Bacteriocytes | -2.7840437 | 11.026951 | -5.4711975 | 0.0005192 | 0.0036146 | 0.2574006 |
| PG 14:0/18:2 | PG | 32 | 2 | FALSE | 192 | Buchnera - Apisum_Bacteriocytes | 8.2533615 | 5.3042134 | 8.7986796 | 1.71E-05 | 0.0006196 | 3.4807478 |
| LPE 19:1 | LPE | 19 | 1 | FALSE | 194 | Buchnera - Apisum_Bacteriocytes | -9.3809127 | 5.4244392 | -9.4669707 | 9.79E-06 | 0.0004429 | 3.9723908 |
| PI 16:1/18:2 | PI | 34 | 3 | FALSE | 197 | Buchnera - Apisum_Bacteriocytes | -0.1062988 | 11.114968 | -0.1902907 | 0.8536444 | 0.8729358 | -6.4581268 |
| LPC 16:1 | LPC | 16 | 1 | FALSE | 198 | Buchnera - Apisum_Bacteriocytes | 4.3872889 | 11.048006 | 6.5427151 | 0.0001514 | 0.0019569 | 1.4571275 |
| PE 32:3 | PE | 32 | 3 | FALSE | 200 | Buchnera - Apisum_Bacteriocytes | 2.9253052 | 9.892486 | 6.0490404 | 0.0002623 | 0.0027058 | 0.925795 |
| PS 18:0/18:1 | PS | 36 | 1 | FALSE | 202 | Buchnera - Apisum_Bacteriocytes | 0.8260002 | 7.9169042 | 0.9379115 | 0.3747249 | 0.4582785 | -6.0209208 |
| LPE 18:3 | LPE | 18 | 3 | FALSE | 203 | Buchnera - Apisum_Bacteriocytes | 0.945518 | 9.9664535 | 1.3629267 | 0.2086606 | 0.2883403 | -5.5628188 |
| PG 14:0/18:1 | PG | 32 | 1 | FALSE | 205 | Buchnera - Apisum_Bacteriocytes | 0.7312582 | 0.3656291 | 1.3523805 | 0.2118744 | 0.2883403 | -5.5755789 |
| PG 18:1/18:1 | PG | 36 | 2 | FALSE | 206 | Buchnera - Apisum_Bacteriocytes | -0.0505625 | 10.58329 | -0.0876399 | 0.9322365 | 0.9479484 | -6.4736948 |
| PE 22:0/18:2 | PE | 40 | 2 | FALSE | 210 | Buchnera - Apisum_Bacteriocytes | 2.1392785 | 6.5426562 | 3.2203598 | 0.0116199 | 0.0293187 | -2.8391534 |
| PC 14:0/16:1 | PC | 30 | 1 | FALSE | 211 | Buchnera - Apisum_Bacteriocytes | 2.314087 | 8.3496738 | 3.8383886 | 0.0046089 | 0.0141391 | -1.9164386 |
| PS 16:1/18:2 | PS | 34 | 3 | FALSE | 218 | Buchnera - Apisum_Bacteriocytes | 1.0269283 | 7.7703387 | 1.7093113 | 0.1243283 | 0.2009234 | -5.1131225 |
| CL 18:1/18:2/18:2/18:2 | CL | 72 | 7 | FALSE | 220 | Buchnera - Apisum_Bacteriocytes | 2.0878074 | 6.608086 | 3.1547864 | 0.0128526 | 0.0314367 | -2.9392726 |
| PG 16:1/18:2 | PG | 34 | 3 | FALSE | 221 | Buchnera - Apisum_Bacteriocytes | -0.2346223 | 10.196366 | -0.3817735 | 0.7121995 | 0.7538486 | -6.3988544 |
| PG 16:0/19:1 | PG | 35 | 1 | FALSE | 223 | Buchnera - Apisum_Bacteriocytes | -2.9319028 | 10.19116 | -4.9903627 | 0.0009489 | 0.0053687 | -0.3388049 |
| PC 20:0/18:2 | PC | 38 | 2 | FALSE | 224 | Buchnera - Apisum_Bacteriocytes | 4.1692631 | 6.0200042 | 2.8236058 | 0.0215221 | 0.0480926 | -3.4485748 |
| LPI 18:1 | LPI | 18 | 1 | FALSE | 225 | Buchnera - Apisum_Bacteriocytes | 0.3345681 | 10.848414 | 0.5802895 | 0.5771 | 0.6408288 | -6.2973667 |
| LPG 18:1 | LPG | 18 | 1 | FALSE | 226 | Buchnera - Apisum_Bacteriocytes | -1.627331 | 9.8160086 | -2.3423384 | 0.0460751 | 0.0877853 | -4.1884125 |
| LPS 18:1 | LPS | 18 | 1 | FALSE | 227 | Buchnera - Apisum_Bacteriocytes | -4.2037169 | 6.2220614 | -7.5761043 | 5.24E-05 | 0.0013551 | 2.4605933 |
| PS 18:0/16:1 | PS | 34 | 1 | FALSE | 229 | Buchnera - Apisum_Bacteriocytes | 0.2575517 | 8.1232134 | 0.4566113 | 0.65964 | 0.7106835 | -6.365257 |
| LPI 18:2 | LPI | 18 | 2 | FALSE | 230 | Buchnera - Apisum_Bacteriocytes | 0.580246 | 11.068552 | 1.0205466 | 0.3362356 | 0.4140043 | -5.9417497 |
| LPS 16:0 | LPS | 16 | 0 | FALSE | 231 | Buchnera - Apisum_Bacteriocytes | -6.0100687 | 6.9525208 | -5.1650462 | 0.0007594 | 0.0045814 | -0.1179263 |
| PI 32:1 | PI | 32 | 1 | FALSE | 232 | Buchnera - Apisum_Bacteriocytes | 0.2484079 | 10.994341 | 0.4707262 | 0.6499358 | 0.7044214 | -6.3582798 |
| PC 18:0/18:1 | PC | 36 | 1 | FALSE | 234 | Buchnera - Apisum_Bacteriocytes | 4.1928081 | 9.4054507 | 8.0447946 | 3.36E-05 | 0.0010131 | 2.8715073 |
| PE 18:1/20:1 | PE | 38 | 2 | FALSE | 235 | Buchnera - Apisum_Bacteriocytes | 2.8694876 | 7.124163 | 6.0267141 | 0.0002691 | 0.0027058 | 0.900918 |
| PC 16:1/16:1 | PC | 32 | 2 | FALSE | 236 | Buchnera - Apisum_Bacteriocytes | 0.6781076 | 10.96685 | 1.1853038 | 0.268658 | 0.3523703 | -5.7689629 |
| PE 18:0/17:1 | PE | 35 | 1 | FALSE | 237 | Buchnera - Apisum_Bacteriocytes | -3.3831158 | 9.6877785 | -4.5873098 | 0.0016135 | 0.0071231 | -0.8671881 |
| PI 14:0/18:2 | PI | 32 | 2 | FALSE | 239 | Buchnera - Apisum_Bacteriocytes | -0.8301162 | 10.356273 | -1.6377291 | 0.1386724 | 0.2182584 | -5.210423 |
| PE 30:1 | PE | 30 | 1 | FALSE | 240 | Buchnera - Apisum_Bacteriocytes | 0.6731136 | 10.917169 | 1.2054078 | 0.2612202 | 0.3464626 | -5.7466101 |
| PE 33:2 | PE | 33 | 2 | FALSE | 241 | Buchnera - Apisum_Bacteriocytes | -2.3196184 | 10.403216 | -3.7001218 | 0.0056437 | 0.0167461 | -2.1189997 |
| PG 17:1/19:1 | PG | 36 | 2 | FALSE | 244 | Buchnera - Apisum_Bacteriocytes | 0.0941322 | 11.549085 | 0.2034795 | 0.843651 | 0.8676184 | -6.4552987 |
| LPG 18:2 | LPG | 18 | 2 | FALSE | 245 | Buchnera - Apisum_Bacteriocytes | -0.3530218 | 9.6503835 | -0.5089882 | 0.623987 | 0.6844948 | -6.3383588 |
| LPG 17:1 | LPG | 17 | 1 | FALSE | 246 | Buchnera - Apisum_Bacteriocytes | -4.2893745 | 8.8072546 | -6.0482652 | 0.0002625 | 0.0027058 | 0.9249325 |
| PE 16:1/26:1 | PE | 42 | 2 | FALSE | 247 | Buchnera - Apisum_Bacteriocytes | 1.5728893 | 7.2700073 | 3.0798289 | 0.0144305 | 0.0343674 | -3.0540832 |
| PE 24:0/18:2 | PE | 42 | 2 | FALSE | 248 | Buchnera - Apisum_Bacteriocytes | 1.5825718 | 7.267022 | 3.1220525 | 0.0135183 | 0.0326241 | -2.9893659 |
| LPE 18:3 | LPE | 18 | 3 | FALSE | 249 | Buchnera - Apisum_Bacteriocytes | 1.9578862 | 11.198715 | 3.3379887 | 0.0097089 | 0.0262286 | -2.6604249 |
| LPC 16:0 | LPC | 16 | 0 | FALSE | 252 | Buchnera - Apisum_Bacteriocytes | -0.7431598 | 8.6129546 | -0.8826346 | 0.4022241 | 0.4821362 | -6.070875 |
| CL 18:1/18:2/18:2/18:3 | CL | 72 | 8 | FALSE | 253 | Buchnera - Apisum_Bacteriocytes | 1.2133681 | 5.6635772 | 1.5594286 | 0.1560814 | 0.2394129 | -5.3144508 |
| CL 18:1/18:1/18:2/18:2 | CL | 72 | 6 | FALSE | 254 | Buchnera - Apisum_Bacteriocytes | 3.0402789 | 6.2385768 | 4.4010522 | 0.0020788 | 0.0081797 | -1.1201327 |
| LPE 14:0 | LPE | 14 | 0 | FALSE | 255 | Buchnera - Apisum_Bacteriocytes | -2.553094 | 7.5873325 | -4.4193636 | 0.0020272 | 0.0081715 | -1.0950228 |
| PS 14:0/18:2 | PS | 32 | 2 | FALSE | 258 | Buchnera - Apisum_Bacteriocytes | 1.4642231 | 7.6581016 | 1.8860164 | 0.0946099 | 0.1630895 | -4.8651242 |
| LPC 16:1 | LPC | 16 | 1 | FALSE | 264 | Buchnera - Apisum_Bacteriocytes | 4.2345141 | 8.7447085 | 6.5990166 | 0.0001424 | 0.0019569 | 1.5154974 |
| PE 18:1/19:1 | PE | 37 | 2 | FALSE | 265 | Buchnera - Apisum_Bacteriocytes | -2.1224984 | 8.5455059 | -4.2324312 | 0.0026263 | 0.008803 | -1.3538062 |
| PE 14:1/16:1 | PE | 30 | 2 | FALSE | 266 | Buchnera - Apisum_Bacteriocytes | 3.3760937 | 9.1185289 | 6.9680949 | 9.64E-05 | 0.0016313 | 1.8872837 |
| PC 14:1/16:1 | PC | 30 | 2 | FALSE | 267 | Buchnera - Apisum_Bacteriocytes | 2.3454015 | 6.8325977 | 4.4935974 | 0.0018317 | 0.0078939 | -0.9937677 |
| LPS 18:0 | LPS | 18 | 0 | FALSE | 268 | Buchnera - Apisum_Bacteriocytes | -0.2203358 | 7.7776429 | -0.3416476 | 0.7410888 | 0.7753588 | -6.4144846 |
| LPC 18:3 | LPC | 18 | 3 | FALSE | 271 | Buchnera - Apisum_Bacteriocytes | 1.2811501 | 9.6519182 | 2.2818107 | 0.0507181 | 0.094639 | -4.2802644 |
| LPI 18:0 | LPI | 18 | 0 | FALSE | 272 | Buchnera - Apisum_Bacteriocytes | -1.181158 | 8.2967249 | -1.3942045 | 0.1993749 | 0.2841485 | -5.5246135 |
| PS 34:4 | PS | 34 | 4 | FALSE | 273 | Buchnera - Apisum_Bacteriocytes | 2.5075877 | 5.5922827 | 1.5232823 | 0.1647641 | 0.2436554 | -5.3615598 |
| PI 18:0/18:1 | PI | 36 | 1 | FALSE | 274 | Buchnera - Apisum_Bacteriocytes | 2.0027754 | 10.954974 | 3.7806681 | 0.0050139 | 0.0151254 | -2.0006937 |
| PG 32:2 | PG | 32 | 2 | FALSE | 278 | Buchnera - Apisum_Bacteriocytes | -9.4630194 | 5.4627679 | -14.889434 | 2.75E-07 | 2.49E-05 | 6.7479758 |
| LPG 16:0 | LPG | 16 | 0 | FALSE | 279 | Buchnera - Apisum_Bacteriocytes | -1.1736465 | 8.7449506 | -1.764766 | 0.1141725 | 0.1895892 | -5.0364216 |
| PC 16:2/18:3 | PC | 34 | 5 | FALSE | 280 | Buchnera - Apisum_Bacteriocytes | 1.1962017 | 5.6863586 | 1.6456545 | 0.1370127 | 0.2175377 | -5.1997498 |
| LPC 18:3 | LPC | 18 | 3 | FALSE | 282 | Buchnera - Apisum_Bacteriocytes | 0.0129424 | 8.7032546 | 0.0225438 | 0.9825457 | 0.9880043 | -6.4776173 |
| PC 15:0/18:2 | PC | 33 | 2 | FALSE | 285 | Buchnera - Apisum_Bacteriocytes | 1.3535057 | 7.3624897 | 2.8794238 | 0.0197181 | 0.0446123 | -3.3624552 |
| PE 17:1/18:2 | PE | 35 | 3 | FALSE | 286 | Buchnera - Apisum_Bacteriocytes | -1.5472804 | 8.4645674 | -2.7854187 | 0.0228534 | 0.0498368 | -3.5075164 |
| LPC 18:0 | LPC | 18 | 0 | FALSE | 287 | Buchnera - Apisum_Bacteriocytes | 0.9945399 | 7.5276148 | 1.4184875 | 0.1924149 | 0.2764055 | -5.4945882 |
| PG 14:0/16:0 | PG | 30 | 0 | FALSE | 288 | Buchnera - Apisum_Bacteriocytes | -2.2229533 | 9.6544538 | -4.607354 | 0.0015706 | 0.007107 | -0.8402954 |
| LPG 18:1 | LPG | 18 | 1 | FALSE | 290 | Buchnera - Apisum_Bacteriocytes | 1.4927254 | 10.116562 | 2.295444 | 0.0496334 | 0.0935796 | -4.2596158 |
| PG 14:0/17:1 | PG | 31 | 1 | FALSE | 294 | Buchnera - Apisum_Bacteriocytes | -1.4416866 | 9.5083878 | -2.4880949 | 0.0365625 | 0.0743574 | -3.9656483 |
| PG 18:0/18:1 | PG | 36 | 1 | FALSE | 295 | Buchnera - Apisum_Bacteriocytes | 1.5118222 | 10.593419 | 2.9661351 | 0.0172193 | 0.0394517 | -3.2288274 |
| PS 18:3/18:3 | PS | 36 | 6 | FALSE | 299 | Buchnera - Apisum_Bacteriocytes | -3.691644 | 2.5798048 | -4.6167323 | 0.0015509 | 0.007107 | -0.8277347 |
| PC 14:0/16:2 | PC | 30 | 2 | FALSE | 300 | Buchnera - Apisum_Bacteriocytes | 0.7339828 | 0.3669914 | 1.3549631 | 0.2110835 | 0.2883403 | -5.57246 |
| PI 15:0/18:2 | PI | 33 | 2 | FALSE | 302 | Buchnera - Apisum_Bacteriocytes | -3.8455614 | 7.1629601 | -5.4910824 | 0.0005068 | 0.0036146 | 0.2812636 |
| PG 16:0/16:0 | PG | 32 | 0 | FALSE | 303 | Buchnera - Apisum_Bacteriocytes | -1.0376418 | 8.6408221 | -1.2026247 | 0.2622397 | 0.3464626 | -5.7497201 |
| PC 16:0/16:1 | PC | 32 | 1 | FALSE | 304 | Buchnera - Apisum_Bacteriocytes | 2.4798228 | 10.252538 | 4.4363501 | 0.0019806 | 0.0081715 | -1.0717766 |
| LPG 18:2 | LPG | 18 | 2 | FALSE | 305 | Buchnera - Apisum_Bacteriocytes | 2.4349126 | 9.4049461 | 3.576298 | 0.0067811 | 0.0194822 | -2.3024582 |
| CL 16:0/18:1/18:2/18:2 | CL | 70 | 5 | FALSE | 306 | Buchnera - Apisum_Bacteriocytes | 3.4479466 | 7.0032013 | 4.4177897 | 0.0020316 | 0.0081715 | -1.097179 |
| LPE 16:2 | LPE | 16 | 2 | FALSE | 308 | Buchnera - Apisum_Bacteriocytes | 1.9232153 | 7.7974631 | 2.9887164 | 0.0166239 | 0.038576 | -3.1940721 |
| PG 34:4 | PG | 34 | 4 | FALSE | 309 | Buchnera - Apisum_Bacteriocytes | -0.1527153 | 8.0539265 | -0.2551762 | 0.8047845 | 0.8323771 | -6.4424097 |
| PG 16:1/17:1 | PG | 33 | 2 | FALSE | 311 | Buchnera - Apisum_Bacteriocytes | -0.8488493 | 9.3936033 | -1.46 | 0.1810071 | 0.2620983 | -5.4425462 |
| PE 44:2 | PE | 44 | 2 | FALSE | 312 | Buchnera - Apisum_Bacteriocytes | 3.4806208 | 5.2056139 | 4.9030422 | 0.0010624 | 0.0058271 | -0.4510597 |
| CL 18:1/18:2/18:3/18:3 | CL | 72 | 9 | FALSE | 314 | Buchnera - Apisum_Bacteriocytes | 1.1017947 | 4.9069196 | 1.6477006 | 0.1365872 | 0.2175377 | -5.1969902 |
| CL 14:0/18:1/18:2/18:2 | CL | 68 | 5 | FALSE | 316 | Buchnera - Apisum_Bacteriocytes | 2.3289685 | 7.3007299 | 4.7225119 | 0.0013468 | 0.0069646 | -0.6870352 |
| LPG 19:1 | LPG | 19 | 1 | FALSE | 317 | Buchnera - Apisum_Bacteriocytes | -1.598307 | 7.9517719 | -2.0221547 | 0.0764486 | 0.1356588 | -4.6677567 |
| CL 14:0/18:1/18:1/18:2 | CL | 68 | 4 | FALSE | 321 | Buchnera - Apisum_Bacteriocytes | 2.0843788 | 7.7122168 | 3.3174115 | 0.0100178 | 0.026665 | -2.6916029 |
| PC 14:1/18:3 | PC | 32 | 4 | FALSE | 322 | Buchnera - Apisum_Bacteriocytes | 2.2353182 | 5.5576435 | 3.382096 | 0.0090801 | 0.0249016 | -2.5937288 |
| LPG 16:1 | LPG | 16 | 1 | FALSE | 323 | Buchnera - Apisum_Bacteriocytes | 2.3747712 | 9.6714083 | 3.6623034 | 0.0059679 | 0.0174225 | -2.1748327 |
| PE 14:1/18:3 | PE | 32 | 4 | FALSE | 326 | Buchnera - Apisum_Bacteriocytes | 2.9659267 | 6.6666706 | 5.8782076 | 0.0003195 | 0.0028915 | 0.7335324 |
| PI 18:3/18:3 | PI | 36 | 6 | FALSE | 329 | Buchnera - Apisum_Bacteriocytes | -1.1177372 | 7.2517992 | -2.1051495 | 0.0670788 | 0.122639 | -4.5452659 |
| PC 20:0/18:3 | PC | 38 | 3 | FALSE | 330 | Buchnera - Apisum_Bacteriocytes | 2.4543265 | 5.4437668 | 4.8272592 | 0.0011729 | 0.0062442 | -0.5494785 |
| PC 16:1/26:1 | PC | 42 | 2 | FALSE | 331 | Buchnera - Apisum_Bacteriocytes | 3.3912075 | 3.9264631 | 2.0970358 | 0.0679432 | 0.1229772 | -4.5573045 |
| CL 16:1/18:1/18:2/18:2 | CL | 70 | 6 | FALSE | 332 | Buchnera - Apisum_Bacteriocytes | 3.571236 | 6.5150598 | 5.5297209 | 0.0004835 | 0.0036146 | 0.327454 |
| CL 16:0/18:1/18:1/18:2 | CL | 70 | 4 | FALSE | 334 | Buchnera - Apisum_Bacteriocytes | 2.8911732 | 6.6443455 | 4.0901633 | 0.0032093 | 0.0103729 | -1.5543311 |
| LPS 16:1 | LPS | 16 | 1 | FALSE | 335 | Buchnera - Apisum_Bacteriocytes | -0.302501 | 7.3442776 | -0.531077 | 0.6092521 | 0.6724063 | -6.3261939 |
| CL 14:0/18:1/18:2/18:3 | CL | 68 | 6 | FALSE | 336 | Buchnera - Apisum_Bacteriocytes | 0.5827974 | 7.178936 | 0.8900346 | 0.398461 | 0.4808096 | -6.0643314 |
| PI 34:4 | PI | 34 | 4 | FALSE | 337 | Buchnera - Apisum_Bacteriocytes | -0.7707822 | 8.0325698 | -1.1554787 | 0.2800094 | 0.3646165 | -5.8016345 |
| PE 16:0/17:2 | PE | 33 | 2 | FALSE | 340 | Buchnera - Apisum_Bacteriocytes | -1.3446858 | 7.6828322 | -1.2651031 | 0.2401285 | 0.32195 | -5.6787203 |
| PE 26:0/16:1 | PE | 42 | 1 | FALSE | 341 | Buchnera - Apisum_Bacteriocytes | 1.8152258 | 4.8281329 | 2.6166547 | 0.0298257 | 0.0628725 | -3.7679054 |
| PC 16:1/28:1 | PC | 44 | 2 | FALSE | 345 | Buchnera - Apisum_Bacteriocytes | 2.4493394 | 2.6586384 | 1.8016377 | 0.1078539 | 0.1824445 | -4.9848299 |
| PG 18:3/18:3 | PG | 36 | 6 | FALSE | 348 | Buchnera - Apisum_Bacteriocytes | -0.5433207 | 6.4072804 | -0.9002089 | 0.3933284 | 0.4778016 | -6.0552614 |
| LPE 16:2 | LPE | 16 | 2 | FALSE | 350 | Buchnera - Apisum_Bacteriocytes | 3.297561 | 8.826835 | 5.4222167 | 0.0005513 | 0.0036439 | 0.1983543 |
| PE 38:4 | PE | 38 | 4 | FALSE | 351 | Buchnera - Apisum_Bacteriocytes | 2.4668556 | 1.2334278 | 1.9043838 | 0.0919387 | 0.1600087 | -4.8387844 |
| LPG 16:1 | LPG | 16 | 1 | FALSE | 352 | Buchnera - Apisum_Bacteriocytes | -0.0520314 | 9.2071277 | -0.0798818 | 0.9382201 | 0.9487031 | -6.4744053 |
| CL 14:0/16:0/18:1/18:2 | CL | 66 | 3 | FALSE | 354 | Buchnera - Apisum_Bacteriocytes | 2.4815662 | 7.1694438 | 3.1611659 | 0.0127268 | 0.0314367 | -2.9295185 |
| PG 14:0/18:3 | PG | 32 | 3 | FALSE | 356 | Buchnera - Apisum_Bacteriocytes | 0.1748465 | 6.8530352 | 0.2756408 | 0.7895492 | 0.8213127 | -6.4365172 |

### 4 Table_GO_lipid_DEGs

This sheet lists selected lipid metabolism-related Gene Ontology (GO) terms identified by GOfuncR, a GO enrichment analysis tool based on gene-regulatory networks and structured ontology relationships. The GO terms were selected based on their relevance to lipid metabolic processes, including parent-child relationships within the GO hierarchy. [GO:0006629 = lipid metabolic process; GO:0006644 = phospholipid metabolic process]

| gene_id | logFC | unshrunk.logFC | logCPM | PValue | adj_PValue | GOALL |
|---|---|---|---|---|---|---|
| LOC100166788 | -9.997709423 | -144269489.1 | 3.53114826 | 4.37E-10 | 2.70E-08 | GO:0006629 |
| LOC100160356 | -9.878346463 | -10.03283814 | 6.594717427 | 4.91E-11 | 6.42E-09 | GO:0006629 |
| LOC100159797 | -9.825231917 | -10.24545163 | 5.594706557 | 9.21E-10 | 4.52E-08 | GO:0006629 |
| LOC100168085 | -9.605880319 | -144269488.7 | 3.213450371 | 4.17E-08 | 8.66E-07 | GO:0006629 |
| LOC100160158 | -8.552300169 | -144269487.6 | 2.645688863 | 8.33E-07 | 1.04E-05 | GO:0006629 |
| LOC100169017 | -8.487646358 | -8.90660006 | 4.100245987 | 5.84E-11 | 6.95E-09 | GO:0006629 |
| LOC100159110 | -8.34893805 | -144269487.4 | 2.19813291 | 3.34E-06 | 3.39E-05 | GO:0006629 |
| LOC100161452 | -8.11991796 | -144269487.2 | 2.284328419 | 2.13E-05 | 1.69E-04 | GO:0006629 |
| LOC100575819 | -7.842709662 | -144269486.9 | 1.732399721 | 5.62E-05 | 3.97E-04 | GO:0006629 |
| LOC100164245 | -7.585282639 | -7.999909989 | 3.541184633 | 1.50E-06 | 1.71E-05 | GO:0006629 |
| LOC100159050 | -7.578629328 | -7.992994218 | 3.651091073 | 1.29E-11 | 2.63E-09 | GO:0006629 |
| LOC100162974 | -7.329251741 | -7.746441729 | 3.087555349 | 1.22E-08 | 3.14E-07 | GO:0006629 |
| LOC100574502 | -7.255351587 | -7.476989063 | 5.596716171 | 1.43E-10 | 1.26E-08 | GO:0006629 |
| LOC100573835 | -7.154272949 | -7.569838502 | 2.987138593 | 8.15E-07 | 1.02E-05 | GO:0006629 |
| LOC100163281 | -7.114912794 | -144269486.2 | 1.599842363 | 1.75E-04 | 0.001069606 | GO:0006629 |
| LOC100166286 | -7.102061775 | -7.217265752 | 4.758071228 | 1.18E-08 | 3.08E-07 | GO:0006629 |
| LOC100169110 | -6.767285246 | -6.813375059 | 5.242963692 | 8.21E-12 | 1.91E-09 | GO:0006629 |
| LOC100167975 | -6.737866242 | -7.151989444 | 2.475464544 | 6.98E-06 | 6.42E-05 | GO:0006629 |
| LOC100169208 | -6.558178586 | -6.624758685 | 4.696198658 | 2.26E-08 | 5.22E-07 | GO:0006629 |
| LOC100160396 | -6.321767025 | -6.734896642 | 2.309590454 | 5.03E-06 | 4.82E-05 | GO:0006629 |
| LOC100163959 | -5.677144138 | -5.743550282 | 3.722907924 | 4.99E-07 | 6.76E-06 | GO:0006629 |
| LOC100574710 | 5.660605183 | 5.663758504 | 8.024079402 | 3.29E-11 | 4.90E-09 | GO:0006629 |
| LOC100165504 | -5.631696479 | -5.688703002 | 3.889380562 | 5.88E-08 | 1.13E-06 | GO:0006629 |
| LOC100571105 | -5.522470115 | -144269484.6 | 0.223833263 | 0.002947733 | 0.012696554 | GO:0006629 |
| LOC100168713 | -5.245574488 | -5.337048716 | 5.430107811 | 1.73E-06 | 1.92E-05 | GO:0006629 |
| LOC100574626 | -4.977080084 | -144269484 | 1.020278879 | 0.008656302 | 0.032691381 | GO:0006629 |
| LOC100159330 | -4.696545221 | -5.103275227 | 1.412890166 | 0.002124044 | 0.009581391 | GO:0006629 |
| LOC100165323 | -4.616444481 | -4.831974677 | 1.863194312 | 1.51E-04 | 9.42E-04 | GO:0006629 |
| LOC100160501 | -4.334760932 | -4.424708023 | 3.466968274 | 2.27E-04 | 0.001345974 | GO:0006629 |
| LOC100164029 | 4.272288152 | 4.274972154 | 7.539887451 | 5.58E-08 | 1.08E-06 | GO:0006629 |
| LOC100569429 | -4.242638365 | -4.332006172 | 2.955895432 | 3.71E-04 | 0.00207249 | GO:0006629 |
| LOC100575760 | -3.922791694 | -4.130244084 | 1.298246164 | 4.29E-04 | 0.002349977 | GO:0006629 |
| LOC100166187 | -3.79537767 | -144269482.8 | 2.659585481 | 0.008122712 | 0.031032924 | GO:0006629 |
| LOC100162817 | 3.580336601 | 3.587227777 | 4.739802717 | 1.01E-07 | 1.73E-06 | GO:0006629 |
| LOC100169226 | -3.560473869 | -3.591412919 | 2.971588347 | 2.01E-05 | 1.61E-04 | GO:0006629 |
| LOC100571472 | -3.55817473 | -3.664915452 | 2.210913118 | 0.001119864 | 0.005472899 | GO:0006629 |
| LOC100169263 | -3.439844117 | -3.459232444 | 3.784901279 | 0.00157375 | 0.007354498 | GO:0006629 |
| LOC100161104 | -3.408200262 | -3.414128282 | 5.022638565 | 2.44E-08 | 5.59E-07 | GO:0006629 |
| LOC100166546 | 3.336612097 | 3.337269803 | 8.401811507 | 4.66E-08 | 9.41E-07 | GO:0006629 |
| LOC100161380 | -3.272822075 | -3.288832743 | 5.101168164 | 2.13E-07 | 3.30E-06 | GO:0006629 |
| LOC100165911 | -3.159984911 | -3.183751292 | 3.819682227 | 1.95E-04 | 0.00117369 | GO:0006629 |
| LOC100569605 | -3.157672203 | -3.2636538 | 1.294864466 | 0.007045733 | 0.027317405 | GO:0006629 |
| LOC100161792 | -3.115286043 | -3.117975064 | 6.988551952 | 1.00E-05 | 8.70E-05 | GO:0006629 |
| LOC100168230 | 3.04343673 | 3.06165573 | 3.120030291 | 2.68E-05 | 2.07E-04 | GO:0006629 |
| LOC100160488 | 3.004591873 | 3.004897225 | 9.096079622 | 4.27E-10 | 2.70E-08 | GO:0006629 |
| LOC100160442 | 2.861668969 | 2.862146264 | 8.028737497 | 8.11E-07 | 1.02E-05 | GO:0006629 |
| LOC100569970 | -2.805735364 | -3.000521015 | 1.775270332 | 0.028907864 | 0.092375403 | GO:0006629 |
| Ipps | -2.78423572 | -2.786803199 | 5.488334484 | 7.43E-07 | 9.46E-06 | GO:0006629 |
| LOC100159637 | -2.749218694 | -2.759021351 | 6.214067035 | 0.007017398 | 0.027224018 | GO:0006629 |
| LOC100164907 | 2.734956518 | 2.742199211 | 5.696080171 | 7.96E-06 | 7.20E-05 | GO:0006629 |
| LOC100160237 | -2.702624247 | -2.778210109 | 2.479799366 | 0.013078066 | 0.046667319 | GO:0006629 |
| LOC100162176 | -2.66416773 | -2.669046997 | 5.122098809 | 0.008682222 | 0.032769954 | GO:0006629 |
| LOC100165462 | 2.657015329 | 2.657319237 | 8.760226858 | 8.16E-07 | 1.02E-05 | GO:0006629 |
| LOC100165835 | -2.473740439 | -2.477105579 | 6.497823728 | 0.009401318 | 0.035111772 | GO:0006629 |
| Pla2g7 | -2.419385328 | -2.423583741 | 5.389668581 | 0.003075339 | 0.013175323 | GO:0006629 |
| LOC100571500 | 2.375258531 | 2.386045299 | 2.872291003 | 0.005364866 | 0.021550282 | GO:0006629 |
| LOC100164244 | -2.255508965 | -2.256618294 | 6.370165342 | 0.020972001 | 0.07023921 | GO:0006629 |
| LOC100163059 | 2.23957405 | 2.240133659 | 7.493431498 | 7.97E-08 | 1.42E-06 | GO:0006629 |
| LOC100168090 | -2.224091099 | -2.230116849 | 4.459538914 | 0.027459863 | 0.088520926 | GO:0006629 |
| LOC100164181 | -2.206597031 | -2.300394415 | 3.512678917 | 0.04305138 | 0.128983022 | GO:0006629 |
| LOC100166021 | 2.19435719 | 2.195500148 | 6.201416327 | 1.10E-06 | 1.33E-05 | GO:0006629 |
| LOC100165996 | -2.071425117 | -2.077565803 | 5.955595443 | 0.009073854 | 0.034097468 | GO:0006629 |
| LOC100572496 | -2.051693986 | -2.074285702 | 2.739493275 | 0.047554154 | 0.14059966 | GO:0006629 |
| LOC100165900 | 1.998081901 | 1.998400933 | 7.149068085 | 1.91E-04 | 0.001149764 | GO:0006629 |
| LOC103310029 | 1.994253432 | 2.098302185 | 1.478456062 | 0.215267367 | 0.472591624 | GO:0006629 |
| LOC100162149 | -1.993498833 | -1.996799129 | 5.033671047 | 0.061614699 | 0.174327834 | GO:0006629 |
| LOC100163267 | -1.964950269 | -1.974722345 | 3.051907796 | 0.008046465 | 0.03078752 | GO:0006629 |
| LOC100161202 | 1.89246941 | 1.894246357 | 6.560640956 | 6.84E-05 | 4.71E-04 | GO:0006629 |
| LOC100164394 | 1.882074355 | 1.882745286 | 6.636159412 | 2.84E-04 | 0.001636835 | GO:0006629 |
| LOC100168304 | 1.869668983 | 1.869985452 | 7.341693407 | 0.001915305 | 0.008730958 | GO:0006629 |
| LOC100163155 | 1.785314413 | 1.785843812 | 6.363741206 | 0.002824468 | 0.012252112 | GO:0006629 |
| LOC100166666 | 1.773196272 | 1.773809185 | 6.506544058 | 3.58E-04 | 0.002010688 | GO:0006629 |
| LOC100168690 | 1.690719176 | 1.691631747 | 6.915006496 | 0.001254815 | 0.006040269 | GO:0006629 |
| LOC100162020 | 1.65625269 | 1.657783926 | 6.789646578 | 0.002452237 | 0.01087915 | GO:0006629 |
| LOC100162356 | 1.652865315 | 1.653383677 | 6.838375451 | 0.003620021 | 0.015144286 | GO:0006629 |
| LOC100159145 | 1.644040016 | 1.64416955 | 8.595599416 | 0.002366906 | 0.010538405 | GO:0006629 |
| Hadh | 1.625767243 | 1.626007025 | 7.852497062 | 0.00382544 | 0.01592049 | GO:0006629 |
| LOC100166744 | 1.582959499 | 1.583532419 | 7.353931391 | 0.003423965 | 0.014432462 | GO:0006629 |
| LOC100165142 | -1.534642321 | -1.535683391 | 5.438773184 | 0.016287591 | 0.056545432 | GO:0006629 |
| LOC100162405 | 1.443904663 | 1.444600934 | 5.856988812 | 0.015102237 | 0.052874335 | GO:0006629 |
| LOC100165236 | 1.441621828 | 1.442243406 | 5.958890222 | 0.012990383 | 0.046406125 | GO:0006629 |
| LOC100167700 | 1.343335606 | 1.343543583 | 7.481374419 | 0.052155247 | 0.151993935 | GO:0006629 |
| LOC100168607 | -1.32616851 | -1.335552799 | 3.881782321 | 0.101126603 | 0.262793812 | GO:0006629 |
| LOC100158759 | 1.260957975 | 1.263405206 | 4.217028299 | 0.074169119 | 0.203120985 | GO:0006629 |
| LOC100159056 | -1.255570247 | -1.258175929 | 4.085036768 | 0.067239101 | 0.18751381 | GO:0006629 |
| LOC100163189 | 1.242776193 | 1.244095066 | 4.724914132 | 0.051198009 | 0.150011313 | GO:0006629 |
| LOC100167163 | -1.208870028 | -1.212151407 | 3.364415129 | 0.11895373 | 0.299994704 | GO:0006629 |
| LOC100160468 | 1.208371483 | 1.208512289 | 8.351729686 | 0.070194171 | 0.19416269 | GO:0006629 |
| LOC100163970 | -1.156731837 | -1.157337822 | 5.828294786 | 0.182581346 | 0.41875736 | GO:0006629 |
| LOC103309064 | 1.133985272 | 1.135101816 | 4.703982692 | 0.124400195 | 0.310431178 | GO:0006629 |
| LOC100165481 | -1.125713862 | -1.127079765 | 4.719032293 | 0.155933869 | 0.37036823 | GO:0006629 |
| LOC100166182 | 1.073805247 | 1.074316889 | 5.72322749 | 0.17511516 | 0.405846566 | GO:0006629 |
| LOC100166338 | -1.067051739 | -1.069307987 | 3.77586295 | 0.207627931 | 0.460301782 | GO:0006629 |
| LOC100161768 | 1.039357296 | 1.041970692 | 3.640709642 | 0.212584873 | 0.468615615 | GO:0006629 |
| LOC100166836 | 1.020470342 | 1.020687689 | 7.66201391 | 0.227050525 | 0.490628234 | GO:0006629 |
| LOC100169456 | 1.006216304 | 1.006542665 | 7.121545111 | 0.241887714 | 0.513850989 | GO:0006629 |
| LOC100165657 | 1.005067501 | 1.005159988 | 8.524879057 | 0.244853189 | 0.518260451 | GO:0006629 |
| LOC100164985 | -0.985702216 | -0.987502525 | 4.183919415 | 0.266443961 | 0.552461637 | GO:0006629 |
| LOC100163593 | 0.94923132 | 0.951598635 | 3.496390127 | 0.290585471 | 0.589357743 | GO:0006629 |
| LOC100169259 | 0.932718368 | 0.934985796 | 3.479669836 | 0.299974954 | 0.603529448 | GO:0006629 |
| LOC100162715 | -0.918701082 | -0.923924278 | 4.075285358 | 0.338600994 | 0.655303368 | GO:0006629 |
| LOC100162304 | 0.892632413 | 0.892891552 | 6.597204368 | 0.402795583 | 0.739988904 | GO:0006629 |
| LOC100167113 | -0.891371914 | -0.894683827 | 4.53572355 | 0.356301103 | 0.68017609 | GO:0006629 |
| LOC100164713 | 0.89082111 | 0.891619109 | 5.656489646 | 0.380772863 | 0.710946156 | GO:0006629 |
| LOC100168099 | -0.883295015 | -0.88556055 | 3.889536579 | 0.360437843 | 0.684451825 | GO:0006629 |
| LOC100162237 | -0.879604346 | -0.880030693 | 6.20287763 | 0.487906366 | 0.838074018 | GO:0006629 |
| LOC100164923 | 0.851333538 | 0.851606267 | 6.530411074 | 0.49922217 | 0.84953956 | GO:0006629 |
| Mvd | -0.848652698 | -0.849121977 | 5.813382692 | 0.465770714 | 0.816803877 | GO:0006629 |
| LOC100164453 | 0.769888572 | 0.770217557 | 6.518010343 | 0.605669009 | 0.95756233 | GO:0006629 |
| LOC100169381 | 0.768825531 | 0.769460955 | 5.102189215 | 0.587686563 | 0.940856524 | GO:0006629 |
| LOC100160824 | 0.766080658 | 0.767489521 | 4.02531992 | 0.490331633 | 0.84081148 | GO:0006629 |
| LOC103310430 | -0.75065475 | -0.753799624 | 3.552064276 | 0.531506736 | 0.885429968 | GO:0006629 |
| LOC100167764 | 0.715377857 | 0.715683427 | 6.805510463 | 0.52000074 | 0.87230414 | GO:0006629 |
| LOC100164505 | -0.699237097 | -0.699854413 | 5.115311149 | 0.629582388 | 0.979894173 | GO:0006629 |
| LOC100162472 | -0.696340018 | -0.697075872 | 6.247606179 | 0.488474597 | 0.838825179 | GO:0006629 |
| LOC100159919 | 0.656175239 | 0.660711164 | 2.549678674 | 0.504268896 | 0.855627865 | GO:0006629 |
| LOC100166849 | 0.632886672 | 0.633053229 | 6.398497581 | 0.642821131 | 0.992662083 | GO:0006629 |
| ACYPI27127 | 0.632820008 | 0.63336772 | 4.974596679 | 0.774475723 | 1 | GO:0006629 |
| LOC100167190 | 0.629146419 | 0.630688997 | 3.752442183 | 0.640952426 | 0.990970607 | GO:0006629 |
| LOC100168459 | 0.625640073 | 0.625896869 | 6.31320255 | 0.829095764 | 1 | GO:0006629 |
| Elovl6 | 0.624596652 | 0.62468992 | 7.682210791 | 0.712260912 | 1 | GO:0006629 |
| LOC100167007 | 0.597304694 | 0.597362274 | 8.365397836 | 0.615920593 | 0.968198424 | GO:0006629 |
| LOC100163305 | 0.589737317 | 0.589903648 | 6.880941934 | 0.743927879 | 1 | GO:0006629 |
| LOC100166007 | -0.574491201 | -0.574651728 | 6.798731802 | 0.903212912 | 1 | GO:0006629 |
| LOC100159250 | -0.570931241 | -0.572058445 | 3.977407357 | 0.715217066 | 1 | GO:0006629 |
| LOC100159658 | 0.549085993 | 0.549481143 | 5.02265222 | 0.792225769 | 1 | GO:0006629 |
| LOC100160697 | 0.547470033 | 0.548752613 | 3.668929152 | 0.680326949 | 1 | GO:0006629 |
| LOC100164628 | 0.542958645 | 0.543330867 | 6.019260244 | 0.854039229 | 1 | GO:0006629 |
| LOC100169073 | 0.521700072 | 0.522896183 | 3.839026163 | 0.788604732 | 1 | GO:0006629 |
| LOC100161189 | 0.521054964 | 0.521980102 | 3.960977192 | 0.808407149 | 1 | GO:0006629 |
| LOC107882160 | -0.518020105 | -0.518672671 | 6.950862522 | 0.626707656 | 0.976843681 | GO:0006629 |
| LOC100164095 | 0.512825551 | 0.512913665 | 7.325281677 | 0.913477513 | 1 | GO:0006629 |
| LOC100573721 | 0.498259776 | 0.499221498 | 3.847548425 | 0.999344419 | 1 | GO:0006629 |
| LOC100164748 | 0.473507779 | 0.473838628 | 6.022681728 | 0.92976426 | 1 | GO:0006629 |
| LOC100162008 | -0.404932303 | -0.406193598 | 3.368558337 | 0.794271951 | 1 | GO:0006629 |
| LOC100166289 | 0.384240476 | 0.384680632 | 4.949678388 | 0.957012175 | 1 | GO:0006629 |
| LOC100168103 | -0.382006604 | -0.382107242 | 6.643910579 | 0.958487907 | 1 | GO:0006629 |
| LOC103309533 | 0.38192658 | 0.383742128 | 2.911828946 | 0.805338225 | 1 | GO:0006629 |
| LOC100163413 | 0.380474975 | 0.381086386 | 4.224472144 | 0.915977423 | 1 | GO:0006629 |
| LOC100163582 | -0.36226551 | -0.377219071 | 1.073731868 | 0.730477972 | 1 | GO:0006629 |
| LOC100574505 | 0.356349119 | 0.35786205 | 2.999933949 | 0.856427893 | 1 | GO:0006629 |
| LOC100169386 | 0.353765115 | 0.353775032 | 10.15972707 | 0.91176682 | 1 | GO:0006629 |
| LOC100163451 | 0.338970828 | 0.339839502 | 3.363051416 | 0.784702687 | 1 | GO:0006629 |
| LOC100163247 | -0.32220765 | -0.322591027 | 7.631144421 | 0.804067475 | 1 | GO:0006629 |
| LOC100159927 | 0.307448883 | 0.307702989 | 5.435018899 | 0.978533546 | 1 | GO:0006629 |
| LOC100161670 | 0.30183188 | 0.301875989 | 7.517798341 | 0.871490268 | 1 | GO:0006629 |
| LOC100162131 | -0.298885375 | -0.298991926 | 6.890544477 | 0.954281088 | 1 | GO:0006629 |
| LOC100166047 | 0.261852049 | 0.262374505 | 3.796699039 | 0.936347987 | 1 | GO:0006629 |
| LOC100160780 | 0.253665839 | 0.254044556 | 4.620951782 | 0.907652554 | 1 | GO:0006629 |
| LOC100160544 | 0.227466109 | 0.227540236 | 6.472296254 | 0.989811503 | 1 | GO:0006629 |
| LOC100159649 | 0.201927174 | 0.202083448 | 5.262808044 | 0.945491884 | 1 | GO:0006629 |
| LOC100573870 | -0.197693988 | -0.204582484 | 0.628956744 | 0.846156098 | 1 | GO:0006629 |
| Atpcl | -0.196825566 | -0.196829736 | 10.19557491 | 0.906987355 | 1 | GO:0006629 |
| LOC100160016 | 0.1748362 | 0.174973675 | 5.017103973 | 0.994575094 | 1 | GO:0006629 |
| LOC100169408 | 0.170528089 | 0.170757139 | 4.508136252 | 0.980884064 | 1 | GO:0006629 |
| LOC100167250 | -0.170234975 | -0.170292465 | 6.163384639 | 0.961081219 | 1 | GO:0006629 |
| LOC100160467 | 0.15045663 | 0.150524827 | 5.835063076 | 0.971548444 | 1 | GO:0006629 |
| LOC100145824 | -0.128698439 | -0.128712204 | 8.456665576 | 0.993378805 | 1 | GO:0006629 |
| LOC100144797 | 0.127165475 | 0.127193035 | 6.862185779 | 0.986723563 | 1 | GO:0006629 |
| LOC100165692 | 0.095153113 | 0.095208685 | 6.068069835 | 0.978619562 | 1 | GO:0006629 |
| LOC100159894 | -0.07413011 | -0.07461767 | 2.858882831 | 0.964194941 | 1 | GO:0006629 |
| LOC100158821 | 0.060695839 | 0.060756352 | 4.714173861 | 0.989137288 | 1 | GO:0006629 |
| LOC100574913 | 0.033246695 | 0.033327035 | 3.38940266 | 0.989811147 | 1 | GO:0006629 |
| LOC100161239 | 0.02067268 | 0.020692287 | 4.802401231 | 0.996410112 | 1 | GO:0006629 |
| LOC100159110 | -8.34893805 | -144269487.4 | 2.19813291 | 3.34E-06 | 3.39E-05 | GO:0006644 |
| LOC100569429 | -4.242638365 | -4.332006172 | 2.955895432 | 3.71E-04 | 0.00207249 | GO:0006644 |
| LOC100166187 | -3.79537767 | -144269482.8 | 2.659585481 | 0.008122712 | 0.031032924 | GO:0006644 |
| LOC100569970 | -2.805735364 | -3.000521015 | 1.775270332 | 0.028907864 | 0.092375403 | GO:0006644 |
| Ipps | -2.78423572 | -2.786803199 | 5.488334484 | 7.43E-07 | 9.46E-06 | GO:0006644 |
| LOC100163059 | 2.23957405 | 2.240133659 | 7.493431498 | 7.97E-08 | 1.42E-06 | GO:0006644 |
| LOC100168304 | 1.869668983 | 1.869985452 | 7.341693407 | 0.001915305 | 0.008730958 | GO:0006644 |
| LOC100163155 | 1.785314413 | 1.785843812 | 6.363741206 | 0.002824468 | 0.012252112 | GO:0006644 |
| LOC100165142 | -1.534642321 | -1.535683391 | 5.438773184 | 0.016287591 | 0.056545432 | GO:0006644 |
| LOC100165236 | 1.441621828 | 1.442243406 | 5.958890222 | 0.012990383 | 0.046406125 | GO:0006644 |
| LOC100160658 | 1.434472088 | 1.43458224 | 10.08985913 | 0.011071483 | 0.040476459 | GO:0006644 |
| LOC100167700 | 1.343335606 | 1.343543583 | 7.481374419 | 0.052155247 | 0.151993935 | GO:0006644 |
| LOC100158759 | 1.260957975 | 1.263405206 | 4.217028299 | 0.074169119 | 0.203120985 | GO:0006644 |
| LOC100159056 | -1.255570247 | -1.258175929 | 4.085036768 | 0.067239101 | 0.18751381 | GO:0006644 |
| LOC100163189 | 1.242776193 | 1.244095066 | 4.724914132 | 0.051198009 | 0.150011313 | GO:0006644 |
| LOC100165481 | -1.125713862 | -1.127079765 | 4.719032293 | 0.155933869 | 0.37036823 | GO:0006644 |
| LOC100166182 | 1.073805247 | 1.074316889 | 5.72322749 | 0.17511516 | 0.405846566 | GO:0006644 |
| LOC100161768 | 1.039357296 | 1.041970692 | 3.640709642 | 0.212584873 | 0.468615615 | GO:0006644 |
| LOC100166836 | 1.020470342 | 1.020687689 | 7.66201391 | 0.227050525 | 0.490628234 | GO:0006644 |
| LOC100169456 | 1.006216304 | 1.006542665 | 7.121545111 | 0.241887714 | 0.513850989 | GO:0006644 |
| LOC100163593 | 0.94923132 | 0.951598635 | 3.496390127 | 0.290585471 | 0.589357743 | GO:0006644 |
| LOC100169259 | 0.932718368 | 0.934985796 | 3.479669836 | 0.299974954 | 0.603529448 | GO:0006644 |
| LOC100162304 | 0.892632413 | 0.892891552 | 6.597204368 | 0.402795583 | 0.739988904 | GO:0006644 |
| LOC100164713 | 0.89082111 | 0.891619109 | 5.656489646 | 0.380772863 | 0.710946156 | GO:0006644 |
| LOC100162237 | -0.879604346 | -0.880030693 | 6.20287763 | 0.487906366 | 0.838074018 | GO:0006644 |
| Mvd | -0.848652698 | -0.849121977 | 5.813382692 | 0.465770714 | 0.816803877 | GO:0006644 |
| LOC100164453 | 0.769888572 | 0.770217557 | 6.518010343 | 0.605669009 | 0.95756233 | GO:0006644 |
| LOC100169381 | 0.768825531 | 0.769460955 | 5.102189215 | 0.587686563 | 0.940856524 | GO:0006644 |
| LOC100160824 | 0.766080658 | 0.767489521 | 4.02531992 | 0.490331633 | 0.84081148 | GO:0006644 |
| LOC103310430 | -0.75065475 | -0.753799624 | 3.552064276 | 0.531506736 | 0.885429968 | GO:0006644 |
| LOC100164505 | -0.699237097 | -0.699854413 | 5.115311149 | 0.629582388 | 0.979894173 | GO:0006644 |
| LOC100159919 | 0.656175239 | 0.660711164 | 2.549678674 | 0.504268896 | 0.855627865 | GO:0006644 |
| LOC100167190 | 0.629146419 | 0.630688997 | 3.752442183 | 0.640952426 | 0.990970607 | GO:0006644 |
| LOC100168459 | 0.625640073 | 0.625896869 | 6.31320255 | 0.829095764 | 1 | GO:0006644 |
| LOC100163305 | 0.589737317 | 0.589903648 | 6.880941934 | 0.743927879 | 1 | GO:0006644 |
| LOC100159250 | -0.570931241 | -0.572058445 | 3.977407357 | 0.715217066 | 1 | GO:0006644 |
| LOC100159658 | 0.549085993 | 0.549481143 | 5.02265222 | 0.792225769 | 1 | GO:0006644 |
| LOC100164628 | 0.542958645 | 0.543330867 | 6.019260244 | 0.854039229 | 1 | GO:0006644 |
| LOC100169073 | 0.521700072 | 0.522896183 | 3.839026163 | 0.788604732 | 1 | GO:0006644 |
| LOC100161189 | 0.521054964 | 0.521980102 | 3.960977192 | 0.808407149 | 1 | GO:0006644 |
| LOC100162008 | -0.404932303 | -0.406193598 | 3.368558337 | 0.794271951 | 1 | GO:0006644 |
| LOC100163451 | 0.338970828 | 0.339839502 | 3.363051416 | 0.784702687 | 1 | GO:0006644 |
| LOC100166047 | 0.261852049 | 0.262374505 | 3.796699039 | 0.936347987 | 1 | GO:0006644 |
| LOC100160780 | 0.253665839 | 0.254044556 | 4.620951782 | 0.907652554 | 1 | GO:0006644 |
| LOC100160544 | 0.227466109 | 0.227540236 | 6.472296254 | 0.989811503 | 1 | GO:0006644 |
| LOC100159649 | 0.201927174 | 0.202083448 | 5.262808044 | 0.945491884 | 1 | GO:0006644 |
| LOC100160016 | 0.1748362 | 0.174973675 | 5.017103973 | 0.994575094 | 1 | GO:0006644 |
| LOC100169408 | 0.170528089 | 0.170757139 | 4.508136252 | 0.980884064 | 1 | GO:0006644 |
| LOC100160467 | 0.15045663 | 0.150524827 | 5.835063076 | 0.971548444 | 1 | GO:0006644 |
| LOC100165692 | 0.095153113 | 0.095208685 | 6.068069835 | 0.978619562 | 1 | GO:0006644 |
| LOC100159894 | -0.07413011 | -0.07461767 | 2.858882831 | 0.964194941 | 1 | GO:0006644 |
| LOC100158821 | 0.060695839 | 0.060756352 | 4.714173861 | 0.989137288 | 1 | GO:0006644 |
| LOC100574913 | 0.033246695 | 0.033327035 | 3.38940266 | 0.989811147 | 1 | GO:0006644 |
| LOC100161239 | 0.02067268 | 0.020692287 | 4.802401231 | 0.996410112 | 1 | GO:0006644 |

### 5 AcetylCoA_transporters

This sheet presents the differential gene expression output generated by the “edgeR” package, comparing viviparous aphid bacteriocytes to whole body samples. It includes only the Acetyl-CoA Transporter 1 genes found in the transcriptome, regardless of significance. Each row corresponds to a gene, with its expression statistics and test results.

| gene_id | logFC | PValue | adj_PValue | Regulation |
|---|---|---|---|---|
| LOC100159262 | 6.0242534 | 2.19E-10 | 1.70E-08 | Upregulated |
| LOC100167419 | 5.6551974 | 5.97E-09 | 1.85E-07 | Upregulated |
| LOC100163344 | 5.2098402 | 5.83E-12 | 1.78E-09 | Upregulated |
| LOC100169490 | 5.1015388 | 2.33E-08 | 5.36E-07 | Upregulated |
| LOC100163610 | 3.8962029 | 1.67E-06 | 1.86E-05 | Upregulated |
| LOC100159922 | 3.8940456 | 1.45E-08 | 3.67E-07 | Upregulated |
| LOC100162627 | 3.6527956 | 2.83E-09 | 1.05E-07 | Upregulated |
| LOC100165404 | 2.4976641 | 2.43E-05 | 0.0001894 | Upregulated |
| LOC100161783 | 0.3609725 | 0.8538321 | 1 | Not significant |
| LOC100160845 | 0.1847519 | 0.9733957 | 1 | Not significant |
| LOC100164131 | -0.5723514 | 0.5466684 | 0.900274 | Not significant |
| LOC100167438 | -0.7184734 | 0.4470611 | 0.7943022 | Not significant |
| LOC100169508 | -0.9239304 | 0.3593153 | 0.68303 | Not significant |
| LOC100161308 | -1.3591138 | 0.0631379 | 0.1778522 | Not significant |
| LOC100163065 | -4.3402657 | 7.88E-05 | 0.000534 | Downregulated |
| LOC100161039 | -4.6517991 | 2.93E-06 | 3.02E-05 | Downregulated |
| LOC100166703 | -7.5586845 | 3.97E-08 | 8.34E-07 | Downregulated |

### 6 Lipid_class_summary_Buchnera

This table summarizes lipid species that were significantly enriched in isolated Buchnera fractions (adjusted p < 0.05, logFC > 1). Lipid species were aggregated by class to compute mean and maximum logFC values and then manually matched to KEGG compound IDs corresponding to the *A. pisum* glycerophospholipid metabolism pathway (api00564). Columns include lipid class, mean logFC, maximum logFC, number of species per class, and mapped KEGG compound ID.

| Class | mean_logFC | max_logFC | n_species | mean_adjP | KEGG_ID |
|---|---|---|---|---|---|
| CL | 2.7416694 | 3.571236 | 8 | 0.0176389 | C05980 |
| DG | 5.9837911 | 12.735845 | 4 | 0.0023415 | C00641 |
| LPC | 3.6714112 | 4.3872889 | 4 | 0.0071274 | C04233 |
| LPE | 2.4727081 | 3.297561 | 4 | 0.0213484 | C04438 |
| LPG | 2.4048419 | 2.4349126 | 2 | 0.0204738 | C05973 |
| PC | 3.4626791 | 11.79527 | 11 | 0.0151531 | C00157 |
| PE | 2.5713877 | 4.0057706 | 21 | 0.0122476 | C00350 |
| PG | 4.8825918 | 8.2533615 | 2 | 0.0215882 | C00344 |
| PI | 2.0027754 | 2.0027754 | 1 | 0.0169744 | C01194 |
| PS | 3.0076733 | 3.0687732 | 2 | 0.0063503 | C02737 |

### 7 api00564_gene_matches_max

This table lists *A. pisum* genes that were significantly differentially expressed in viviparous bacteriocytes (adjusted p < 0.05) and successfully mapped to KEGG enzyme entries using their ENTREZ IDs. For genes with multiple transcripts or paralogs per enzyme, the entry with the largest absolute log₂ fold change was retained. Columns include ENTREZ ID, maximum logFC, and the number of gene models represented per enzyme. Data was manually verified against differential expression data.

| kegg.names | labels | all.mapped | type | x | y | width | height | mol.data | mol.col |
|---|---|---|---|---|---|---|---|---|---|
| 100166182 | 100166182 |  | gene | 521 | 368 | 46 | 17 | NA | #FFFFFF |
| 100573870 | 100573870 |  | gene | 112 | 301 | 46 | 17 | NA | #FFFFFF |
| 100169605 | 100169605 | 100169605 | gene | 813 | 234 | 46 | 17 | 1.5090256 | #FCCD91 |
| 100160590 | 100160590 |  | gene | 759 | 303 | 46 | 17 | NA | #FFFFFF |
| 100160383 | 100160383 | 100160383 | gene | 911 | 334 | 46 | 17 | 1.4667669 | #FCCD91 |
| 100169456 | 100169456 |  | gene | 1031 | 326 | 46 | 17 | NA | #FFFFFF |
| 100168865 | 100168865 | 100168865,100169250,103308679 | gene | 1143 | 345 | 46 | 17 | -6.9948703 | #A6CEE3 |
| 100159222 | 100159222 | 100159222 | gene | 1143 | 325 | 46 | 17 | -7.1713119 | #A6CEE3 |
| 100161708 | 100161708 |  | gene | 911 | 545 | 46 | 17 | NA | #FFFFFF |
| 100160369 | Eas |  | gene | 1031 | 532 | 46 | 17 | NA | #FFFFFF |
| 100569429 | 100569429 | 100569429 | gene | 813 | 210 | 46 | 17 | -4.2426384 | #A6CEE3 |
| 100160810 | 100163089 | 100160810,100163089 | gene | 426 | 482 | 46 | 17 | -6.1588864 | #A6CEE3 |
| 100164616 | 100164616 | 100164616,100165537 | gene | 562 | 461 | 46 | 17 | 2.7313623 | #FDBF6F |
| 100569429 | 100569429 | 100569429 | gene | 813 | 615 | 46 | 17 | -4.2426384 | #A6CEE3 |
| 100164497 | 100164497 |  | gene | 277 | 414 | 46 | 17 | NA | #FFFFFF |
| 100165692 | 100165692 |  | gene | 278 | 608 | 46 | 17 | NA | #FFFFFF |
| 100166666 | 100166666 | 100166666 | gene | 251 | 301 | 46 | 17 | 1.7731963 | #FDBF6F |
| 100160658 | 100160658 | 100160658 | gene | 170 | 214 | 46 | 17 | 1.4344721 | #FCCD91 |
| 100160153 | 100160153 |  | gene | 170 | 273 | 46 | 17 | NA | #FFFFFF |
| 100159211 | 100159211 |  | gene | 1031 | 233 | 46 | 17 | NA | #FFFFFF |
| 100159649 | 100159649 |  | gene | 759 | 590 | 46 | 17 | NA | #FFFFFF |
| 100168842 | 100168842 | 100168842 | gene | 911 | 641 | 46 | 17 | 1.6374889 | #FDBF6F |
| 100167870 | 100167870 |  | gene | 813 | 641 | 46 | 17 | NA | #FFFFFF |
| 100161387 | 100161387 | 100161387,100168842 | gene | 911 | 660 | 46 | 17 | 1.673151 | #FDBF6F |
| 100163059 | 100163059 | 100163059 | gene | 371 | 590 | 46 | 17 | 2.239574 | #FDBF6F |
| 100161387 | 100161387 | 100161387,100168842 | gene | 911 | 254 | 46 | 17 | 1.673151 | #FDBF6F |
| 100167700 | 100167700 |  | gene | 588 | 647 | 46 | 17 | NA | #FFFFFF |
| 100159211 | 100159211 |  | gene | 1031 | 639 | 46 | 17 | NA | #FFFFFF |
| 100168842 | 100168842 | 100168842 | gene | 911 | 235 | 46 | 17 | 1.6374889 | #FDBF6F |
| 100162008 | 100162008 |  | gene | 346 | 692 | 46 | 17 | NA | #FFFFFF |
| 100161189 | 100161189 |  | gene | 277 | 794 | 46 | 17 | NA | #FFFFFF |
| 100160900 | 100160900 |  | gene | 162 | 599 | 46 | 17 | NA | #FFFFFF |
| 100165481 | 100165481 |  | gene | 162 | 784 | 46 | 17 | NA | #FFFFFF |
| 100169605 | 100169605 | 100169605 | gene | 345 | 842 | 46 | 17 | 1.5090256 | #FCCD91 |
| 100167870 | 100167870 |  | gene | 565 | 614 | 46 | 17 | NA | #FFFFFF |
| 100165269 | 100165269 | 100165269 | gene | 1031 | 487 | 46 | 17 | 1.6321515 | #FDBF6F |
| 100160143 | 100160143 | 100160143 | gene | 725 | 564 | 46 | 17 | 1.5659713 | #FCCD91 |

### 8 LC_MSMS_sample_detail

Tissue samples prepared for LC-MS/MS analyses. The table included the number of replicates, sample description and sample amount used in the experiment.

| Sample name | Replicate | Description | Sample amount | Unit |
|---|---|---|---|---|
| Apisum_Appendages | 1 | Appendages of pea aphid (without the core body)_aphid_frozen tissue | 1.58 | mg |
|  | 2 |  | 2.24 |  |
|  | 3 |  | 4.43 |  |
| Apisum_Bacteriocytes | 1 | Specialised organ housing the symbiont (*B. aphidicola* )_pelleted | 733 | cells |
|  | 2 |  | 1067 |  |
|  | 3 |  | 2600 |  |
| Apisum_WholeBody | 1 | Whole body of pea aphid (*A. pisum* )_frozen whole aphid | 2.23 | mg |
|  | 2 |  | 2.33 |  |
|  | 3 |  | 4.69 |  |
| Buchnera | 1 | Isolated endosymbionts of pea aphid (*B. aphidicola* ) 500 aphids pelleted | 1.51 x 10^6 | cells |
|  | 2 | Isolated endosymbionts of pea aphid (*B. aphidicola* ) 400 aphids pelleted | 2.18 x 10^6 |  |
|  | 3 | Isolated endosymbionts of pea aphid (*B. aphidicola* ) 600 aphids pelleted | 9.34 x 10^6 |  |

### 9 sequence_QC

Summary of RNA-seq quality control and preprocessing metrics. For each sample, the table reports the number of input read pairs, surviving read pairs after Trimmomatic quality trimming, percentage of surviving read pairs, percentage of rRNA reads identified by RiboDetector, total unambiguous reads mapped to the Acyrthosiphon pisum genome using BBSplit, and FastQC quality metrics including GC content, sequence duplication, average read length, and sequence length range.

| Sample | Input read pairs | Read pairs after trimming | Reads retained (%) | Total Sequences | GC content (%) | Unique sequences (%) | Mean read length (bp) | Read length range (bp) | Duplicate sequences (%) | rRNA reads (%) | Total unambiguous reads mapped to A. pisum |
|---|---|---|---|---|---|---|---|---|---|---|---|
| WB_F1 | 10040642 | 8185407 | 81.5 | 32741628 | 40 | 50.9 | 102.89 | 18-151 | 49.1 | 4.1 | 13999376 |
| WB_F2 | 10169122 | 8052988 | 79.2 | 32211952 | 41 | 48.5 | 116.38 | 18-151 | 51.5 | 4.5 | 13859582 |
| WB_F3 | 14588688 | 7819215 | 53.6 | 31276860 | 40 | 51.0 | 66.64 | 18-76 | 49.0 | 3.5 | 13531252 |
| WB_F4 | 13758847 | 7115681 | 51.7 | 28462724 | 41 | 49.1 | 66.56 | 18-76 | 50.9 | 7.8 | 11675570 |
| WB_F5 | 14017458 | 7308028 | 52.1 | 29232112 | 41 | 49.2 | 67.01 | 18-76 | 50.8 | 7.5 | 12047851 |
| WBMale_F1 | 13788819 | 7279749 | 52.8 | 29118996 | 42 | 41.2 | 66.49 | 18-76 | 58.8 | 6.5 | 12270065 |
| WBMale_F2 | 14543257 | 7339633 | 50.5 | 29358532 | 41 | 42.5 | 66.86 | 18-76 | 57.5 | 7.3 | 11981212 |
| WBMale_F3 | 14761701 | 7739807 | 52.4 | 30959228 | 41 | 43.4 | 66.69 | 18-76 | 56.6 | 5.9 | 12924476 |
| BactVivi_F1 | 8149213 | 6657524 | 81.7 | 26630096 | 36 | 32.4 | 97.88 | 18-151 | 67.6 | 6.0 | 11763841 |
| BactVivi_F2 | 9349676 | 7309722 | 78.2 | 29238888 | 39 | 30.7 | 106.63 | 18-151 | 69.3 | 10.2 | 12667941 |
| BactVivi_F3 | 14347728 | 7887505 | 55.0 | 31550020 | 39 | 31.2 | 67.20 | 18-76 | 68.8 | 6.5 | 14072485 |
| BactVivi_F4 | 15736748 | 8579261 | 54.5 | 34317044 | 39 | 30.2 | 67.39 | 18-76 | 69.8 | 6.2 | 15098010 |
| BactVivi_F5 | 12859863 | 6806149 | 52.9 | 27224596 | 39 | 33.3 | 66.95 | 18-76 | 66.7 | 7.0 | 11820226 |
| BactMale_F1 | 10019382 | 8038126 | 80.2 | 32152504 | 37 | 45.5 | 123.22 | 18-151 | 54.5 | 2.4 | 14115973 |
| BactMale_F2 | 5834579 | 4675424 | 80.1 | 18701696 | 37 | 47.4 | 104.06 | 18-151 | 52.6 | 10.0 | 7775974 |
| BactMale_F3 | 15978426 | 9043874 | 56.6 | 36175496 | 39 | 38.5 | 67.27 | 18-76 | 61.5 | 5.2 | 15395182 |
| BactMale_F4 | 7574481 | 4117103 | 54.4 | 16468412 | 37 | 42.3 | 64.24 | 18-76 | 57.7 | 9.1 | 6063880 |
| BactMale_F5 | 15043091 | 8544314 | 56.8 | 34177256 | 40 | 35.2 | 67.00 | 18-76 | 64.8 | 5.7 | 14946326 |
| BactOvi_F1 | 7392649 | 6017712 | 81.4 | 24070848 | 35 | 37.1 | 92.55 | 18-151 | 62.9 | 5.3 | 10609191 |
| BactOvi_F2 | 9439585 | 7609912 | 80.6 | 30439648 | 36 | 30.9 | 109.63 | 18-151 | 69.1 | 6.7 | 13423029 |

### 10 siRNA_list

Details of siRNAs assays, including transcript accession numbers, target gene names and descriptions, siRNA sequence identifiers, strand orientation, length, nucleotide sequence, synthesis state, and production batch information.

| **siRNA** | **Transcript_accession** | **Target_gene** | **Gene_description** | **Sequence Name** | **Strand** | **Length (mer)** | **Sequence (5' to 3')** | **State** | **Lot No./Batch No.** |
|---|---|---|---|---|---|---|---|---|---|
| ApB744 | XM_008186273.3 | LOC100160658 | glycerol-3-phosphate dehydrogenase [NAD(+)], cytoplasmic | ApB744 siRNA1 S | sense | 21 | **UAGCUCUGAUUUUCACAAAUdTdT** | Annealed | 537178 |
|  |  |  |  | ApB744 siRNA1 AS | antisense | 21 | **auuugugaaaucagagcuadTdT** |  | 537179 |
|  |  |  |  | ApB744 siRNA2 S | sense | 21 | **CUCUCAUGCUCGACUUAAGUdTdT** | Annealed | 537180 |
|  |  |  |  | ApB744 siRNA2 AS | antisense | 21 | **acuaagucgagcaugagagdTdT** |  | 537181 |
| ApW584 | XM_001950868 | LOC100159262 | acetyl-coenzyme A transporter 1 | #ApW584 siRNA1 S | sense | 21 | **CUACAGCUAUGUUUCAUUUCUdTdT** | Annealed | 539585 |
|  |  |  |  | #ApW584 siRNA1 AS | antisense | 21 | **agaaugaacauagcuguagdTdT** |  | 539586 |
|  |  |  |  | #ApW584 siRNA4 S | sense | 21 | **CUCAGGACAUAGUAGUUGAdTdT** | Annealed | 539687 |
|  |  |  |  | #ApW584 siRNA4 AS | antisense | 21 | **ucaacucuacuauguccugagdTdT** |  | 539588 |
| Universal control |  |  |  | Universal control | sense<br>antisense | 21<br>21 | The universal negative control siRNA (Nippon Gene) used in this study is designed based on a validated sequence that exhibits no homology to any known eukaryotic gene. The exact sequence is proprietary and not publicly disclosed by the manufacturer. | Annealed | NEGS240920<br>NEGAS240920 |

### 11 PrimerList

List of primers used for quantitative PCR analyses, including primer sequences, primer length, expected amplicon size, corresponding target gene, associated siRNA target (if applicable), and amplification efficiency.

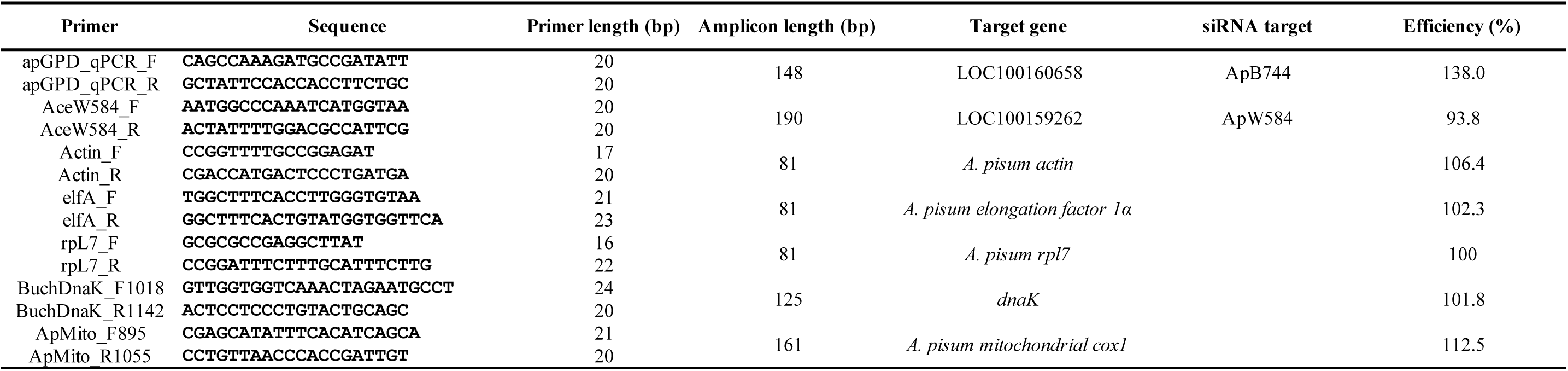

## References

1. Douglas, A. E. (1994). Symbiotic interactions. Oxford University Press.

2. Douglas, A. E. (1998). Nutritional interactions in insect-microbial symbioses: Aphids and their symbiotic bacteria *Buchnera*. Annual Review of Entomology, 43, 17–37. 10.1146/annurev.ento.43.1.17

3. Margulis L. (1971). Symbiosis and evolution. Scientific American, 225(2), 48–57. 10.1038/scientificamerican0871-48

4. Moran, N. A. (2007). Symbiosis as an adaptive process and source of phenotypic complexity. Proc. Natl. Acad. Sci. USA, 104(Suppl. 1), 8627–8633. 10.1073/pnas.0611659104

5. Shigenobu, S., & Wilson, A. C. (2011). Genomic revelations of a mutualism: the pea aphid and its obligate bacterial symbiont. Cellular and molecular life sciences, 68(8), 1297–1309. 10.1007/s00018-011-0645-2

6. Shigenobu, S., Watanabe, H., Hattori, M., Sakaki, Y., & Ishikawa, H. (2000). Genome sequence of the endocellular bacterial symbiont of aphids *Buchnera* sp. APS. Nature, 407(6800), 81–86. 10.1038/35024074

7. Koga, R., Meng, X.Y., Tsuchida, T., and Fukatsu, T. (2012). Cellular mechanism for selective vertical transmission of an obligate insect symbiont at the bacteriocyte-embryo interface. Proc. Natl. Acad. Sci. USA 109, E1230–E1237. 10.1073/pnas.1119212109

8. Koga, R., Tsuchida, T., Sakurai, M., & Fukatsu, T. (2007). Selective elimination of aphid endosymbionts: Effects of antibiotic dose and host genotype, and fitness consequences. FEMS Microbiology Ecology, 60(2), 229–239. 10.1111/j.1574-6941.2007.00284.x

9. Douglas, A.E. (1992). Requirement of pea aphids *(Acyrthosiphon pisum*) for their symbiotic bacteria. Entomol. Exp. Appl. 65, 195–198. 10.1111/j.1570-7458.1992.tb01643.x

10. Gil, R., Sabater-Muñoz, B., Latorre, A., Silva, F. J., & Moya, A. (2002). Extreme genome reduction in *Buchnera* spp.: toward the minimal genome needed for symbiotic life. Proc. Natl. Acad. Sci. USA, 99(7), 4454–4458. 10.1073/pnas.062067299

11. Baumann, P., Baumann, L., Lai, C. Y., Rouhbakhsh, D., Moran, N. A., & Clark, M. A. (1995). Genetics, physiology, and evolutionary relationships of the genus *Buchnera*: intracellular symbionts of aphids. Annual review of microbiology, 49, 55–94. 10.1146/annurev.mi.49.100195.000415

12. Moran, N. A. (1996). Accelerated evolution and Muller’s rachet in endosymbiotic bacteria. Proc. Natl. Acad. Sci. USA, 93(7), 2873–2878. 10.1073/pnas.93.7.2873

13. Mira, A., Ochman, H., & Moran, N. A. (2001). Deletional bias and the evolution of bacterial genomes. Trends in genetics, 17(10), 589–596. 10.1016/s0168-9525(01)02447-7

14. Blow, F., Bueno, E., Clark, N., Zhu, D. T., Chung, S. H., Güllert, S., Schmitz, R. A., & Douglas, A. E. (2020). B-vitamin nutrition in the pea aphid-*Buchnera* symbiosis. Journal of insect physiology, 126, 104092. 10.1016/j.jinsphys.2020.104092

15. McCutcheon, J. P., Garber, A. I., Spencer, N., & Warren, J. M. (2024). How do bacterial endosymbionts work with so few genes? PLoS Biology, 22(4), e3002577. 10.1371/journal.pbio.3002577

16. Cronan, J. E., & Vagelos, P. R. (1972). Metabolism and function of the membrane phospholipids of *Escherichia coli*. Biochimica et biophysica acta, 265(1), 25–60. 10.1016/0304-4157(72)90018-4

17. Cronan J. E. (2003). Bacterial membrane lipids: where do we stand?. Annual review of microbiology, 57, 203–224. 10.1146/annurev.micro.57.030502.090851

18. Sohlenkamp, C., & Geiger, O. (2016). Bacterial membrane lipids: diversity in structures and pathways. FEMS microbiology reviews, 40(1), 133–159. 10.1093/femsre/fuv008

19. Kanehisa, M., Goto, S., Kawashima, S., & Nakaya, A. (2002). The KEGG databases at GenomeNet. Nucleic acids research, 30(1), 42–46. 10.1093/nar/30.1.42

20. Shigenobu, S., Michellod, D., & Tan, K. X. Y. (2026). Lipid metabolism in the aphid-*Buchnera* symbiosis: An underexplored dimension of insect physiology. Current Opinion in Insect Science, 78, 101581. 10.1016/j.cois.2026.101581

21. McCutcheon, J. P., & Moran, N. A. (2011). Extreme genome reduction in symbiotic bacteria. Nature reviews. Microbiology, 10(1), 13–26. 10.1038/nrmicro2670

22. Houk, E. J., Griffiths, G. W., Hadjokas, N. E., & Beck, S. D. (1977). Peptidoglycan in the cell wall of the primary intracellular symbiote of the pea aphid. Science (New York, N.Y.), 198(4315), 401–403. 10.1126/science.198.4315.401

23. Munson, M. A., Baumann, P., & Kinsey, M. G. (1991). *Buchnera* gen. nov. and *Buchnera aphidicola* sp. nov., a taxon consisting of the mycetocyte-associated, primary endosymbionts of aphids. International Journal of Systematic and Evolutionary Microbiology, 41(4), 566–568. 10.1099/00207713-41-4-566

24. Tan, B.K., Bogdanov, M., Zhao, J., Dowhan, W., Raetz, C.R.H., and Guan, Z. (2012). Discovery of a cardiolipin synthase utilizing phosphatidylethanolamine and phosphatidylglycerol as substrates. Proc. Natl. Acad. Sci. USA 109, 16504–16509. 10.1073/pnas.1212797109.

25. Nishijima, S., Asami, Y., Uetake, N., Yamagoe, S., Ohta, A., & Shibuya, I. (1988). Disruption of the *Escherichia coli cls* gene responsible for cardiolipin synthesis. Journal of bacteriology, 170(2), 775–780. 10.1128/jb.170.2.775-780.1988

26. The International Aphid Genomics Consortium, “Genome sequence of the pea aphid *Acyrthosiphon pisum*,” PLoS Biol., vol. 8, no. 2, Art. no. e1000313, 2010. 10.1371/journal.pbio.1000313

27. Huerta-Cepas, J., et al. (2019). eggNOG 5.0: A hierarchical, functionally and phylogenetically annotated orthology resource based on 5090 organisms and 2502 viruses. Nucleic Acids Research, 47(D1), D309–D314. 10.1093/nar/gky1085

28. Luan, J.-B., Chen, W., Hasegawa, D.K., Simmons, A.M., Wintermantel, W. M., Ling, K.-S., Fei, Z., Liu, S.-S., and Douglas, A.E. (2015). Metabolic coevolution in the bacterial symbiosis of whiteflies and related plant sap-feeding insects. Genome Biol. Evol. 7, 2635–2647. 10.1093/gbe/evv170

29. Wilson, A. C. C., & Duncan, R. P. (2015). Signatures of host/symbiont genome coevolution in insect nutritional endosymbioses. Proc. Natl. Acad. Sci. USA, 112(33), 10255–10261. 10.1073/pnas.1423305112

30. Zhang, Y. M., & Rock, C. O. (2008). Membrane lipid homeostasis in bacteria. Nature reviews. Microbiology, 6(3), 222–233. 10.1038/nrmicro1839

31. Gidden, J., Denson, J., Liyanage, R., Ivey, D. M., & Lay, J. O. (2009). Lipid compositions in *Escherichia coli* and *Bacillus subtilis* during growth as determined by MALDI-TOF and TOF/TOF mass spectrometry. International journal of mass spectrometry, 283(1-3), 178–184. 10.1016/j.ijms.2009.03.005

32. Houk, E. J., Griffiths, G. W., & Beck, S. D. (1976). Lipid metabolism in the symbiotes of the pea aphid, *Acyrthosiphon pisum*. Comparative biochemistry and physiology. B, Comparative biochemistry, 54(3), 427–431. 10.1016/0305-0491(76)90270-4

33. Jiang, F., Ryan, M. T., Schlame, M., Zhao, M., Gu, Z., Klingenberg, M., Pfanner, N., & Greenberg, M. L. (2000). Absence of cardiolipin in the *crd1* null mutant results in decreased mitochondrial membrane potential and reduced mitochondrial function. Journal of Biological Chemistry, 275(29), 22387–22394. 10.1074/jbc.M909868199

34. Ren, M., Phoon, C. K. L., & Schlame, M. (2014). Metabolism and function of mitochondrial cardiolipin. Progress in Lipid Research, 55, 1–16. 10.1016/j.plipres.2014.04.001

35. Tan, K. X. Y., & Shigenobu, S. (2025). Targeted disruption of the *cls* gene in *Buchnera aphidicola* impairs membrane integrity and host symbiont dynamics. iScience, 28(8), 113178. 10.1016/j.isci.2025.113178

36. Furse, S., & Scott, D. J. (2016). Three-dimensional distribution of phospholipids in gram negative bacteria. Biochemistry, 55(34), 4742–4747. 10.1021/acs.biochem.6b00541

37. Matsumoto K. (2001). Dispensable nature of phosphatidylglycerol in *Escherichia coli*: dual roles of anionic phospholipids. Molecular microbiology, 39(6), 1427–1433. 10.1046/j.1365-2958.2001.02320.x

38. Soltani, M., Metzger, P., & Largeau, C. (2005). Fatty acid and hydroxy acid adaptation in three gram-negative hydrocarbon-degrading bacteria in relation to carbon source. Lipids, 40(12), 1263–1272. 10.1007/s11745-005-1494-x

39. Subczynski, W. K., Pasenkiewicz-Gierula, M., Widomska, J., Mainali, L., & Raguz, M. (2017). High cholesterol/low cholesterol: effects in biological membranes: a review. Cell biochemistry and biophysics, 75(3-4), 369–385. 10.1007/s12013-017-0792-7

40. Mileykovskaya, E., & Dowhan, W. (2009). Cardiolipin membrane domains in prokaryotes and eukaryotes. Biochimica et biophysica acta, 1788(10), 2084–2091. 10.1016/j.bbamem.2009.04.003

41. Yeh, J. I., Chinte, U., & Du, S. (2008). Structure of glycerol-3-phosphate dehydrogenase, an essential monotopic membrane enzyme involved in respiration and metabolism. Proc. Natl. Acad. Sci. USA, 105(9), 3280–3285. 10.1073/pnas.0712331105

42. Li, Y., Park, H., Smith, T. E., Moran, N. A., & Singh, N. (2019). Gene family evolution in the pea aphid based on chromosome-level genome assembly. Molecular Biology and Evolution, 36(10), 2143–2156. 10.1093/molbev/msz138

43. Shigenobu, S., et al. (2022). Genomic and transcriptomic analyses of the subterranean termite *Reticulitermes speratus*: Gene duplication facilitates social evolution. Proc. Natl. Acad. Sci. USA, 119(3), e2110361119. 10.1073/pnas.2110361119

44. Whittle, M., Barreaux, A. M. G., Bonsall, M. B., Ponton, F., & English, S. (2021). Insect-host control of obligate, intracellular symbiont density. Proceedings of the Royal Society B: Biological Sciences, 288(1963), 20211993. 10.1098/rspb.2021.1993

45. Herren, J. K., Paredes, J. C., Schüpfer, F., Arafah, K., Bulet, P., & Lemaitre, B. (2014). Insect endosymbiont proliferation is limited by lipid availability. eLife, 3, e02964. 10.7554/eLife.02964

46. Griffiths, G. W., & Beck, S. D. (1977). In vivo sterol biosynthesis by pea aphid symbiotes as determined by digitonin and electron microscopic autoradiography. Cell and tissue research, 176(2), 179–190. 10.1007/BF00229461

47. Dambekodi, P. C., & Gilliland, S. E. (1998). Incorporation of cholesterol into the cellular membrane of *Bifidobacterium longum*. Journal of dairy science, 81(7), 1818–1824. 10.3168/jds.S0022-0302(98)75751-0

48. Lin, M., & Rikihisa, Y. (2003). *Ehrlichia chaffeensis* and *Anaplasma phagocytophilum* lack genes for lipid A biosynthesis and incorporate cholesterol for their survival. Infection and immunity, 71(9), 5324–5331. 10.1128/IAI.71.9.5324-5331.2003

49. Lin, M., Grandinetti, G., Hartnell, L. M., Bliss, D., Subramaniam, S., & Rikihisa, Y. (2020). Host membrane lipids are trafficked to membranes of intravacuolar bacterium *Ehrlichia chaffeensis*. Proc. Natl. Acad. Sci. USA, 117(14), 8032–8043. 10.1073/pnas.1921619117

50. Nakabachi, A., Shigenobu, S., Sakazume, N., Shiraki, T., Hayashizaki, Y., Carninci, P., Ishikawa, H., Kudo, T., & Fukatsu, T. (2005). Transcriptome analysis of the aphid bacteriocyte, the symbiotic host cell that harbors an endocellular mutualistic bacterium, *Buchnera*. Proc. Natl. Acad. Sci. USA, 102(15), 5477–5482. 10.1073/pnas.0409034102

51. Derré, I., Swiss, R., & Agaisse, H. (2011). The lipid transfer protein CERT interacts with the *Chlamydia* inclusion protein IncD and participates to ER-*Chlamydia* inclusion membrane contact sites. PLoS Pathogens, 7(6), e1002092. 10.1371/journal.ppat.1002092

52. Elwell, C. A., et al. (2011). *Chlamydia trachomatis* co-opts GBF1 and CERT to acquire host sphingomyelin for distinct roles during intracellular development. PLoS Pathogens, 7(9), e1002198. 10.1371/journal.ppat.1002198

53. J. Ren, Q. Wang, X. Zhang, Y. Cao, J. Wu, J. Tian, Y. Yu, Q. Gong, Z. Kong, Control of rhizobia endosymbiosis by coupling ER expansion with enhanced UPR. Adv. Sci. 2025, 12, 2414519. 10.1002/advs.202414519

54. Feng, H., Edwards, N., Anderson, C. M. H., Althaus, M., Duncan, R. P., Hsu, Y. C., Luetje, C. W., Price, D. R. G., Wilson, A. C. C., & Thwaites, D. T. (2019). Trading amino acids at the aphid-*Buchnera* symbiotic interface. Proc. Natl. Acad. Sci. USA, 116(32), 16003–16011. 10.1073/pnas.1906223116

55. Simonet, P., Gaget, K., Balmand, S., Ribeiro Lopes, M., Parisot, N., Buhler, K., Duport, G., Vulsteke, V., Febvay, G., Heddi, A., Charles, H., Callaerts, P., & Calevro, F. (2018). Bacteriocyte cell death in the pea aphid/*Buchnera* symbiotic system. Proc. Natl. Acad. Sci. USA, 115(8), E1819–E1828. 10.1073/pnas.1720237115

56. Nakabachi, A., Yamashita, A., Toh, H., Ishikawa, H., Dunbar, H. E., Moran, N. A., & Hattori, M. (2006). The 160-kilobase genome of the bacterial endosymbiont *Carsonella*. Science (New York, N.Y.), 314(5797), 267. 10.1126/science.1134196

57. Husnik, F., & McCutcheon, J. P. (2016). Repeated replacement of an intrabacterial symbiont in the tripartite nested mealybug symbiosis. Proc. Natl. Acad. Sci. USA, 113(37), E5416–E5424. 10.1073/pnas.1603910113

58. 58. McCutcheon, J. P., & von Dohlen, C. D. (2011). An interdependent metabolic patchwork in the nested symbiosis of mealybugs. Current biology, 21(16), 1366–1372. 10.1016/j.cub.2011.06.051

59. Rao, Q., Rollat-Farnier, P. A., Zhu, D. T., Santos-Garcia, D., Silva, F. J., Moya, A., Latorre, A., Klein, C. C., Vavre, F., Sagot, M. F., Liu, S. S., Mouton, L., & Wang, X. W. (2015). Genome reduction and potential metabolic complementation of the dual endosymbionts in the whitefly *Bemisia tabaci*. BMC genomics, 16(1), 226. 10.1186/s12864-015-1379-6

60. Sakai, R., Goto-Inoue, N., Yamashita, H., Aimoto, N., Kitai, Y., & Maruyama, T. (2023). Smart utilization of betaine lipids in the giant clam *Tridacna crocea*. iScience, 26(7), 107250. 10.1016/j.isci.2023.107250

61. Kanbe, T., and Akimoto, S.i. (2009). Allelic and genotypic diversity in long-term asexual populations of the pea aphid, *Acyrthosiphon pisum*, in comparison with sexual populations. Mol. Ecol. 18, 801–816. 10.1111/j.1365-294X.2008.04077

62. 62. Poliakov, A., Russell, C. W., Ponnala, L., Hoops, H. J., Sun, Q., Douglas, A. E., & van Wijk, K. J. (2011). Large-scale label-free quantitative proteomics of the pea aphid-*Buchnera* symbiosis. Molecular & Cellular Proteomics, 10(6), M110.007039. 10.1074/mcp.M110.007039

63. Houk, E. J., & McLean, D. L. (1974). Isolation of the primary intracellular symbiote of the pea aphid, *Acyrthosiphon pisum*. Journal of Invertebrate Pathology. 10.1016/0022-2011(74)90190-6

64. Tsugawa, H., et al. (2020). A lipidome atlas in MS-DIAL 4. Nature Biotechnology, 38, 1159–1163. 10.1038/s41587-020-0531-2

65. Mohamed, A., Molendijk, J., & Hill, M. M. (2020). Lipidr: A software tool for data mining and analysis of lipidomics datasets. Journal of Proteome Research, 19(7), 2890–2897. 10.1021/acs.jproteome.0c00082

66. Oksanen, J., et al. (2026). vegan: Community Ecology Package. 10.32614/CRAN.package.vegan

67. Harrell, F. E., Jr. (2023). Hmisc: Harrell Miscellaneous. 10.32614/CRAN.package.Hmisc

68. Langfelder, P., Zhang, B., & Horvath, S. (2008). Defining clusters from a hierarchical cluster tree: The Dynamic Tree Cut package for R. Bioinformatics, 24(5), 719–720. 10.1093/bioinformatics/btm563

69. Hahsler, M., Piekenbrock, M., & Doran, D. (2019). dbscan: Fast density-based clustering with R. Journal of Statistical Software, 91(1), 1–30. 10.18637/jss.v091.i01

70. Pedersen, T. L. (2025). ggraph: An implementation of Grammar of Graphics for graphs and networks. 10.32614/CRAN.package.ggraph

71. Csárdi, G., et al. (2026). igraph: Network analysis and visualization. 10.32614/CRAN.package.igraph

72. Wickham, H., and Sievert, C. (2009). ggplot2: Elegant Graphics for Data Analysis (Springer). 10.1007/978-0-387-98141-3

73. Bolger, A. M., Lohse, M., & Usadel, B. (2014). Trimmomatic: A flexible trimmer for Illumina sequence data. Bioinformatics, 30(15), 2114–2120. 10.1093/bioinformatics/btu170

74. S. Andrews, “FastQC: A Quality Control Tool for High Throughput Sequence Data.” Accessed: Jun. 23, 2024. [Online]. Available: http://www.bioinformatics.babraham.ac.uk/projects/fastqc/

75. Ewels, P., Magnusson, M., Lundin, S., & Käller, M. (2016). MultiQC: Summarize analysis results for multiple tools and samples in a single report. Bioinformatics, 32(19), 3047–3048. 10.1093/bioinformatics/btw354

76. Deng, Z. L., Münch, P. C., Mreches, R., & McHardy, A. C. (2022). Rapid and accurate identification of ribosomal RNA sequences via deep learning. Nucleic Acids Research, 50(10), e60. 10.1093/nar/gkac112

77. Patro, R., Duggal, G., Love, M. I., Irizarry, R. A., & Kingsford, C. (2017). Salmon provides fast and bias-aware quantification of transcript expression. Nature Methods, 14(4), 417–419. 10.1038/nmeth.4197

78. Soneson, C., Love, M. I., & Robinson, M. D. (2016). Differential analyses for RNA-seq: Transcript-level estimates improve gene-level inferences. F1000Research, 4, 1521. 10.12688/f1000research.7563.2

79. Robinson, M. D., McCarthy, D. J., & Smyth, G. K. (2010). edgeR: A Bioconductor package for differential expression analysis of digital gene expression data. Bioinformatics, 26(1), 139–140. 10.1093/bioinformatics/btp616

80. Legeai, F., Shigenobu, S., Gauthier, J.-P., Colbourne, J., Rispe, C., Collin, O., Richards, S., Wilson, A. C. C., Murphy, T., & Tagu, D. (2010). AphidBase: A centralized bioinformatic resource for annotation of the pea aphid genome. Insect Molecular Biology, 19(Suppl. 2), 5–12. 10.1111/j.1365-2583.2009.00930.x

81. Young, M. D., Wakefield, M. J., Smyth, G. K., & Oshlack, A. (2010). Gene ontology analysis for RNA-seq: Accounting for selection bias. Genome Biology, 11(2), R14. 10.1186/gb-2010-11-2-r14

82. Grote, S. (2026). GOfuncR: Gene ontology enrichment using FUNC. Bioconductor. 10.18129/B9.bioc.GOfuncR

83. Luo, W., & Brouwer, C. (2013). Pathview: An R/Bioconductor package for pathway-based data integration and visualization. Bioinformatics, 29(14), 1830–1831. 10.1093/bioinformatics/btt285

84. 84. JGraph (2021). diagrams.net (draw.io), version 15.5.2. https://www.diagrams.net/

85. 85. R Core Team (2021). R: A language and environment for statistical computing. https://www.R-project.org/

86. 86. Wickham, H., Hester, J., Chang, W., and Bryan, J. (2022). devtools: Tools to make developing R packages easier. https://devtools.r-lib.org/

87. Signorell, A. (2023). DescTools: Tools for Descriptive Statistics. https://andrisignorell.github.io/DescTools/

88. 88. Wickham, H., François, R., Henry, L., Müller, K., and Vaughan, D. (2023). dplyr: A grammar of data manipulation. https://dplyr.tidyverse.org.

89. 89. Kassambara, A. (2023). ggpubr: ‘ggplot2’ Based Publication Ready Plots. https://rpkgs.datanovia.com/ggpubr/

90. Min, S. H., & Zhou, J. (2021). smplot: An R package for easy and elegant data visualization. Frontiers in Genetics, 12, 802894. 10.3389/fgene.2021.802894

## References

1. 1. R Core Team (2021). R: A language and environment for statistical computing. https://www.R-project.org/

2. 2. Wickham, H., Hester, J., Chang, W., and Bryan, J. (2022). devtools: Tools to make developing R packages easier. https://devtools.r-lib.org/

3. Signorell, A. (2023). DescTools: Tools for Descriptive Statistics. https://andrisignorell.github.io/DescTools/

94. Wickham, H., François, R., Henry, L., Müller, K., and Vaughan, D. (2023). dplyr: A Grammar of Data Manipulation. https://dplyr.tidyverse.org.

5. Wickham, H., and Sievert, C. (2009). ggplot2: Elegant Graphics for Data Analysis (Springer). 10.1007/978-0-387-98141-3

6. 6. JGraph (2021). diagrams.net (draw.io), version 15.5.2. https://www.diagrams.net/

7. Nagele, R. G., Kosciuk, M. C., Wang, S. M., Spero, D. A., & Lee, H. (1985). A method for preparing quick-frozen, freeze-substituted cells for transmission electron microscopy and immunocytochemistry. Journal of Microscopy, 139(3), 291–301. 10.1111/j.1365-2818.1985.tb02645.x

8. Tan, K.X.Y., and Shigenobu, S. (2024). In vivo interference of pea aphid endosymbiont *Buchnera groEL* gene by synthetic peptide nucleic acids. Sci. Rep. 14, 5378. 10.1038/s41598-024-55179-2.

9. Shigenobu, S., Watanabe, H., Hattori, M., Sakaki, Y., and Ishikawa, H. (2000). Genome sequence of the endocellular bacterial symbiont of aphids *Buchnera* sp. APS. Nature 407, 81–86. 10.1038/35024074.

10. Sohlenkamp, C., and Geiger, O. (2016). Bacterial membrane lipids: Diversity in structures and pathways. FEMS Microbiol. Rev. 40, 133–159. 10.1093/femsre/fuv008.

11. Kanehisa, M., Goto, S., Kawashima, S., and Nakaya, A. (2002). The KEGG databases at GenomeNet. Nucleic Acids Res. 30, 42–46. 10.1093/nar/30.1.42

12. Luo, W., & Brouwer, C. (2013). Pathview: An R/Bioconductor package for pathway-based data integration and visualization. Bioinformatics, 29(14), 1830–1831. 10.1093/bioinformatics/btt285

